# More than a flying syringe: Using functional traits in vector-borne disease research

**DOI:** 10.1101/501320

**Authors:** Lauren J. Cator, Leah R. Johnson, Erin A. Mordecai, Fadoua El Moustaid, Thomas R.C. Smallwood, Shannon L. LaDeau, Michael A. Johansson, Peter J. Hudson, Michael Boots, Matthew B. Thomas, Alison G. Power, Samraat Pawar

## Abstract

Many important endemic and emerging diseases are vector-borne. The functional traits of vectors affect not just pathogen transmission rates, but also the fitness and population dynamics of these animals themselves. Increasing empirical evidence suggests that vector traits vary significantly at time scales relevant to transmission dynamics. Currently, an understanding of how this variation in key traits impacts transmission is hindered by a lack of both empirical data and theoretical methods for mechanistically incorporating traits into transmission models. Here, we present a framework for incorporating both intrinsic and environment-driven variation in vector traits into empirical and theoretical vector-borne disease research. This framework mechanistically captures the effect of trait variation on vector fitness, the correlation between vector traits, and how these together determine transmission dynamics. We illustrate how trait-based vector-borne disease modelling can make novel predictions, and identify key steps and challenges in the construction, empirical parameterisation and validation of such models, as well as the organization and prioritization of data collection efforts.

## I. INTRODUCTION

Vector-borne diseases (VBDs) remain a serious threat to health and economic welfare worldwide. At least one-third of the human population is thought to be impacted by mosquito-borne diseases alone (WHO, 2018). Brazil loses an estimated 1.3 billion USD in productivity each year to Chagas disease, which is transmitted by triatomine bugs (Lee *et al*., 2013). VBDs include re-emerging infections, such as dengue (San Martín *et al*., 2010; Dick *et al*., 2012), and new threats, such as Chikungunya, Zika, and Lyme disease (Mead, 2015; CDC, 2016; Faria *et al*., 2016). The impacts of VBDs are not only medical. Many important and emerging diseases of plants (e.g., Tomato yellow leaf curl virus, cassava mosaic virus, and citrus greening virus (Taylor *et al*., 2016)) and livestock (e.g., bluetongue (Wilson & Mellor, 2009)) are transmitted by vectors.

VBD transmission dynamics are driven by a system of interconnected population abundances that vary over time and, usually, also over space. Evidence from a wealth of empirical and theoretical studies indicates that the behaviour and life history of vectors are key determinants of these VBD dynamics (reviewed below). The importance of vector abundance and life history in transmission dynamics is reflected by the fact that almost all VBD models incorporate vector biting rate, competence, adult survival, and density (Jeger *et al*., 2004; Reiner *et al*., 2013). These aspects of vector biology can be described as traits.

A trait is any measurable feature of an individual organism. Specifically, a trait is functional when it directly (e.g., mortality rate/probability) or indirectly (e.g., development time) determines individual fitness (McGill *et al*., 2006; Gibert *et al*., 2015) (henceforth we use “functional trait” and “trait” synonymously). In natural populations, trait variation between individuals, and within individuals over time is ubiquitous and has been shown to alter population and community level processes (Imura, Toquenaga, & Fujii, 2003; McGill *et al*., 2006; Agashe, 2009; Gibert *et al*., 2015). It is widely accepted that vector traits are also important for transmission and they are likely to be temporally and spatially variable (Smith *et al*., 2014). While some models incorporate these traits, most do not accommodate trait variation. Mechanistically incorporating trait variation into transmission models should therefore improve the ability of models to predict transmission through time and space.

Trait variation can change transmission dynamics, either by changing vector abundances on similar time scales as pathogen transmission, or by directly changing transmission potential itself (e.g., vector biting rate or competence). This is partly due to the fact, that the majority of animal and plant disease vectors are arthropods, and therefore have significantly shorter generation times (and thus, operate at faster timescales) than their host (May & Anderson 1979). Furthermore, arthropods are small ectotherms, which means their abundances vary with fluctuations in environmental variables. Work from other fields indicates that both the mean and variation of a trait’s distribution can have strong effects on the dynamics of populations (Norberg *et al*., 2001; Bolnick *et al*., 2011; Gibert *et al*., 2015). Trait variation has now begun to be identified as essential variables that are key for understanding the state and future trajectories of biodiversity and ecosystems (Kissling *et al*., 2018). Disease dynamics are strongly nonlinear, which means that variation in traits over time and across individuals can have compounding effects on transmission (Lloyd-Smith *et al*., 2005; Martin *et al*., 2019). Until we incorporate trait variation into VBD transmission models our ability to predict transmission dynamics at longer temporal and larger spatial scales will remain limited, like any ecological model that lacks a mechanistic basis (Getz *et al*., 2018). To incorporate trait variation robustly, the mechanistic links between biological parameters, and how their effects on vector population fitness can change population dynamics and transmission rates need to be considered.

There are two related challenges to understanding the parameter space in which trait variation could affect transmission. First, for most vector species, we lack basic data on traits underlying these parameters, and how they vary. Researchers are frequently forced to infer parameter values (for example, using the time it takes a mosquito to produce a clutch of eggs to infer biting rate) or use data from related species to parameterise models (for example, Johnson *et al*. 2015, Mordecai *et al*. 2013). Second, while there are many statistical, correlative methods, we lack a theoretical framework to mechanistically incorporate parameter variation (through trait variation) into transmission models.

Here, we propose a trait-based framework as the way forward on both empirical and theoretical fronts for making more accurate VBD transmission dynamic predictions. We begin by reviewing empirical evidence for trait variation in vectors and the degree which it has (or in some cases has not) been incorporated into previous transmission models. We then describe the trait-based modelling framework and provide an example of how this approach can produce novel predictions and allows for investigation into the degree to which different traits drive transmission dynamics and underlying fitness effects. Next we outline how this can be operationalized, both in terms of incorporating traits into transmission models, and data collection efforts. Finally, we discuss further challenges in the way of implementing a trait-based approach in VBD research.

## II. TRAIT VARIATION IN VECTORS

Variation in vector traits can be grouped into three primary types: 1) variation across the lifespan of an individual; 2) variation within a population; and 3) environmentally driven variation (Fig. 1).

**Figure 1.**
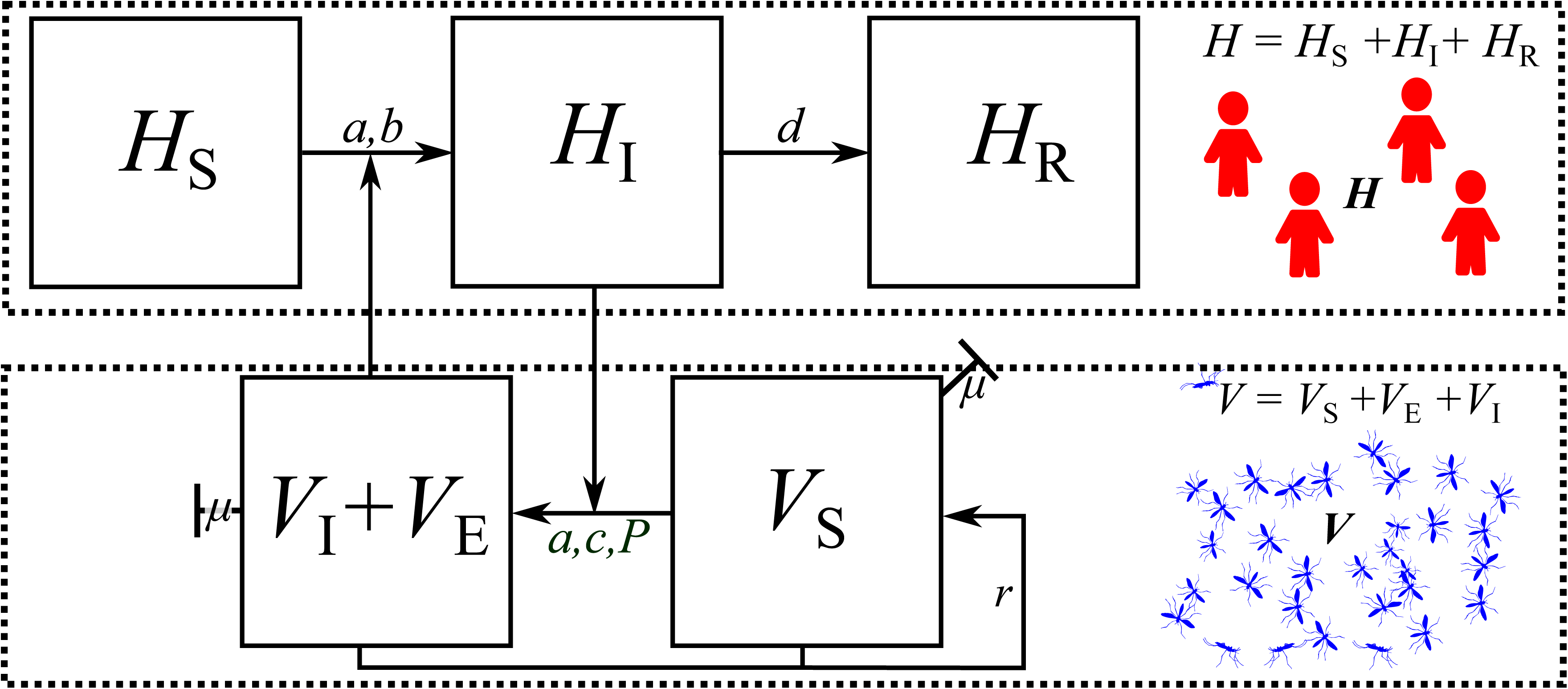
Types of trait variation found in all VBD systems. **A:** Across-individuals: for example, variation in a trait (*z*) within a population at a fixed point in time. The distribution (p(*z*) around the mean (Z̅), represents amount individual variation not measurement error. For example, we could measure the probability of biting of individuals in a population at a particular age (based on data reviewed in Murdock et al. 2017). **B**: Within-individual, over time: for example, biting probability may vary over the vector’s lifespan (based on data reviewed in Murdock et al. 2017). Such variation is quantifiable as a continuous time-dependent function *dz/dt*, where *dz* is the (differential) change in trait variation change with time (*dt*). **C:** Environment-driven: For example, biting rate varies unimodally with temperature (based on Mordecai et al. 2013). Such variation is quantifiable as a continuous environment state-dependent function, *dz*/*dE*. **D:** Combined variation: The three types of trait variation may appear in combination. For example, across-individual trait variation may change over time. The upper line represents change in mean and variance of the trait, while the lower line represents change only in the mean. We use derivatives for quantifying over-time, with-environment and combined types of trait variation to represent the idealized scenario that the trait value function varies continuously and smoothly with respect to another variable. In reality, it may not always be possible to express these as smooth functions for empirical reasons.

### (1) Variation across the lifespan of an individual

For transmission to occur a vector must survive the time it takes from acquiring the parasite to the parasite becoming infectious (extrinsic incubation period or EIP), which can be a large proportion of the vector life span. Therefore, older vector individuals are; 1) more likely to become infected because they are more likely to have been exposed, 2) more likely to be infectious because they are more likely to have survived EIP, and 3) are more likely to transmit the pathogen onwards because they are more likely to bite subsequent hosts after becoming infectious. Therefore, trait values at the end of life disproportionately contribute to transmission. Vector behaviour and life history may vary over the lifetime of an individual also due to intrinsic processes such as aging. There is evidence for age-specific vector competence (Soliman *et al*., 1993), immune function (Hillyer *et al*., 2005; Christensen *et al*., 2005; Laughton, Fan, & Gerardo, 2014), flight performance (Nayar & Sauerman, 1973) and feeding behaviour (Alto, Lounibos, & Juliano, 2003; Den Otter, Tchicaya, & Schutte, 2008; Bohbot *et al*., 2013). When multiple life stages of the vector are involved in transmission, transmission efficiency may vary with stage (Caraco *et al*., 2002; Coletta-Filho *et al*., 2014). All these time-dependent changes in vector traits could lead to significant variation in the number of infectious vectors and their contact rates with hosts.

### (2) Variation within a population

Variation within a population (for example due to intrinsic genetic variation) can lead to subgroups of the vector population having disproportionate effects on the average population fitness and transmission potential. There are multiple examples of this kind of within-population variation in vectors. For example, the nutritional status of vectors can alter both their vector competence and several behavioural parameters linked to VBD transmission (Takken *et al*., 2013; Shapiro *et al*., 2016). Body size varies within populations (Renshaw, Service, & Birley, 1994; De Xue, Edman, & Scott, 1995; Kindlmann & Dixon, 2003) and can drive significant variation in feeding, assimilation, and respiration, and therefore development and mortality rates (Brown *et al*., 2004; Savage, 2004; Amarasekare & Savage, 2012). Inter-individual variation in age-specific mortality is important for transmission (Clements & Paterson, 1981; Harrington *et al*., 2001, 2008; Styer *et al*., 2007). Recent evidence from several different systems shows that vectors infected with a variety of pathogens exhibit altered foraging behaviours (reviewed in (Murdock, Luckhart, & Cator, 2017)). These types of variation between individuals in key traits such as biting rate, host preference, and longevity ultimately impact VBD pathogen transmission rate.

### (3) Environmentally driven variation

The majority of vectors are small ectotherms, so their behaviour and life history are sensitive to their environment. Variation due to environmental drivers, may have both short- or long-term effects on vector traits. At present most of the data on this kind of variation comes from investigations of temperature. Many studies have measured an effect of variation in environmental temperature on vector life history (Kersting, Satar, & Uygun, 1999; Bayoh & Lindsay, 2003; Delatte *et al*., 2009; Ciota *et al*., 2014) and competence (Kramer, Hardy, & Presser, 1983; Murral *et al*., 1996; Dohm, O’Guinn, & Turell, 2002; Paweska, Venter, & Mellor, 2002; Wittmann, Mellor, & Baylis, 2002). Other environmental variables, such as humidity, can also directly affect vector life history (Wittmann *et al*., 2002; Costa *et al*., 2010).

### (4) Traits are mechanistically linked

Not only do vector traits vary but multiple traits also co-vary because they are mechanistically linked. For example, mosquitoes infected with bird malaria parasites exhibit reduced fecundity, which in turn increases longevity (Vézilier *et al*., 2012). These kinds of trait covariances, often appearing in the form of life-history trade-offs (Charnov 1993), have implications for both vector population fitness (and therefore abundance) and transmission rate. Therefore, any theoretical framework that incorporates vector traits and their variation must also account for the mechanistic relationships between traits. There is evidence that this type of incorporation of multiple, mechanistically-linked traits can yield new insights in transmission of VBDs as well as non-VBDs (Mordecai *et al*., 2013, 2017; Molnár *et al*., 2017). For example, recent work has incorporated metabolic traits into micro- and macro-parasite disease transmission (Molnár *et al*., 2017).

In summary, any theoretical framework for VBD should be able to capture changes in transmission dynamics emerging from trait variation (including for those not directly related to transmission), correlations between traits, and the resulting effects on vector population dynamics. We now consider past approaches towards incorporating traits into mathematical VBD models.

## III. CURRENT APPROACHES TO INCORPORATING TRAITS INTO TRANSMISSION DYNAMICS

### (1) Classical compartment models

Classical compartment models focus on the frequency of different (Susceptible, Exposed, Infected, Recovered) sub-populations of the host and vector, often ignoring fluctuations in absolute abundances of the two species. For example, the Ross-Macdonald type model for malaria transmission by a mosquito (Fig. 2, SI Section 2), on which many current VBD models (including non-mosquito vectors) are based (MacDonald, 1957; May & Anderson, 1979; Jeger *et al*., 2004; Smith *et al*., 2012; Reiner *et al*., 2013) focuses exclusively on the parameters governing transmission rate of the pathogen between susceptible and infected vector and host subpopulations, most of which are mosquito traits. It yields a relatively simple equation for the basic reproduction number of the disease (*R*_0_)—the number of new infectious cases that would arise from a single infectious case introduced into a fully susceptible host population—which quantifies its transmission potential or risk (MacDonald, 1957; Smith *et al*., 2012) (see SI section 2 for derivation):

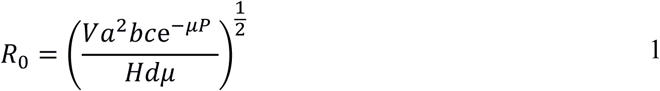

**Figure 2.**
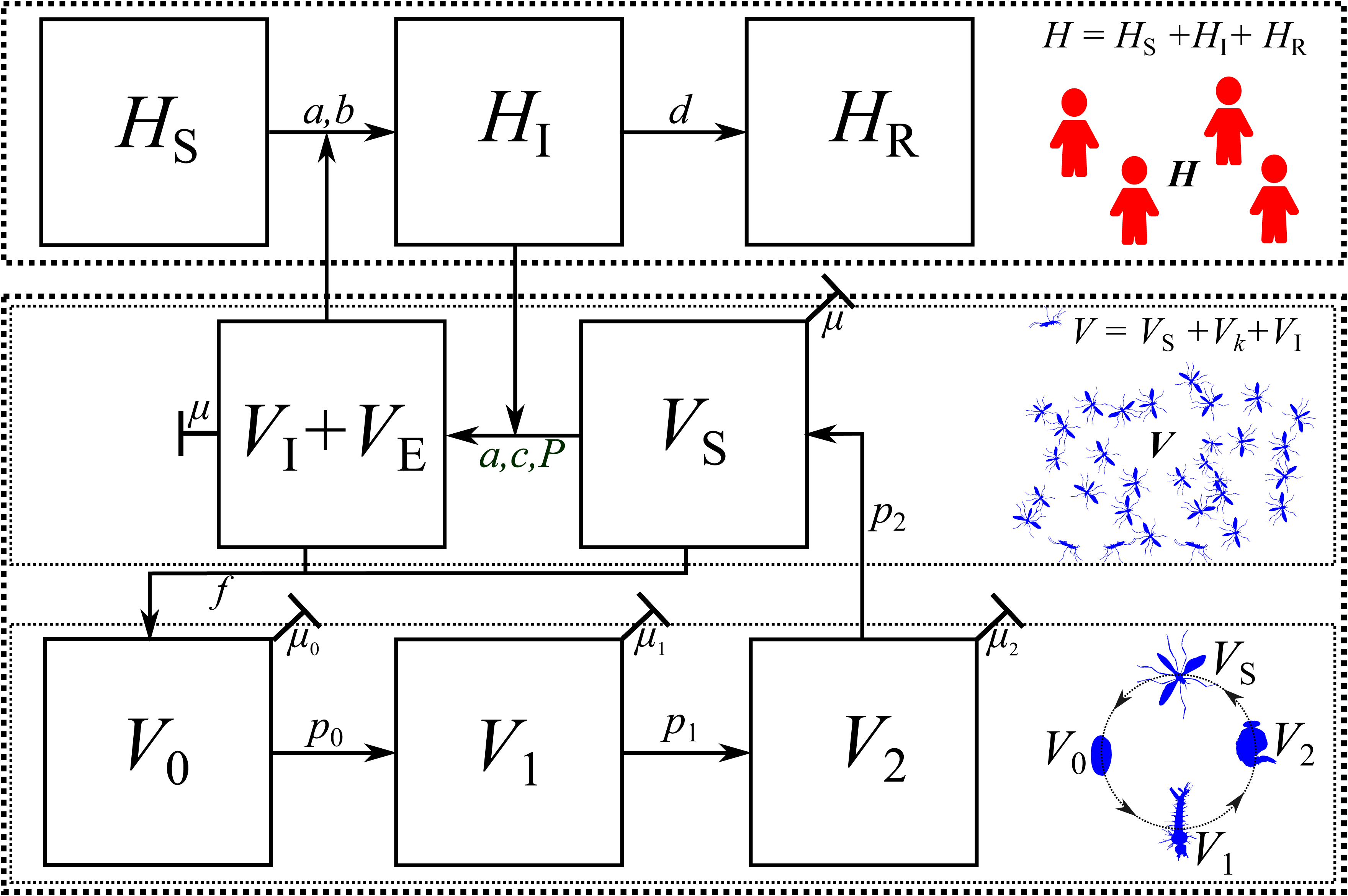
Traits in classical compartment models. As an illustration, we show here a Ross-Macdonald type model for malaria transmission by a mosquito vector. Most of the vector parameters in this model — *a, b, c, P*, *μ* — are directly measurable vector traits (*a*, *μ*), or can be derived from underlying vector and parasite traits (*b, c, P*). The corresponding equations and parameter definitions are shown in SI Section 2. *H* is the total host density which is made up of susceptible (*H_S_*), Infected (*H_I_*) and Recovered (*H_S_*) sub-groups. Total vector density (*V*) is made up for Infected (*V_I_*), Exposed (*V_E_*) and Susceptible (*V_S_*) sub-groups.

Here, *V* is vector density, *a* is per-vector biting rate, *b* is the proportion of the bites by infective mosquitoes that produce infection in susceptible humans, *c* is the proportion of bites by susceptible mosquitoes on infectious humans that infect mosquitoes (thus, *bc* is vector competence), *μ* is adult vector mortality rate, *P* is the extrinsic incubation period of the pathogen within the vector, *H* is host density, and *d* is the rate at which infected hosts recover and acquire immunity (Fig 2).

Compartment models do implicitly incorporate traits. The vector (*a, b, c, P*, *μ*) and host (*d*) parameters used in this approach are all essentially traits that are either directly-measurable properties of the vector itself (*a*, *μ*) or which can be derived from underlying traits of the vector and parasite (*b, c, P*). However, in these models, it is assumed that vector (and host) traits do not affect total vector or host population size and that these traits do not vary across individuals or over time (Fig. 1). Furthermore, all of the vector’s biology and ecology are represented by its adult (often female) biting rate, survival, extrinsic incubation period, competence, and abundance. All of these traits are assumed to be independent of each other despite the fact that they are known to trade-off and feedback on each other with potentially compounding effects on transmission. For example, biting rate (*a*) is assumed to have no effect on mortality (*μ*) or vector population density (*V*) and does not vary across individuals or over time. All individual hosts and vectors are modelled as having an identical risk of infection that remains constant over time and vectors interact exclusively with the target host. This effectively reduces the vector to a homogeneous vehicle to transmit pathogens– a “flying syringe”.

### (2) Compartment models with vector population dynamics

Classical compartment models have been extended to incorporate vector population dynamics, by adding vector life-stage compartments (May & Anderson, 1979; Anderson & May, 1979; Hoshi, Higa, & Chaves, 2014; Johnson *et al*., 2018; Ng *et al*., 2018) (Fig. 3). This introduces additional parameters for the vector’s life history, all of which are directly measurable vector traits (also called demographic rates, e.g., mortality, fecundity), or parameters which can be derived from underlying traits (e.g. the transition probabilities (*p_i_*’s) in Fig. 3 can be derived from stage-specific survivorship and development time). These models do relax the assumption that vector population dynamics are constant, but do not incorporate trait variation, mechanistic links between traits, or allow trait variation to drive vector population dynamics.

**Figure 3.**
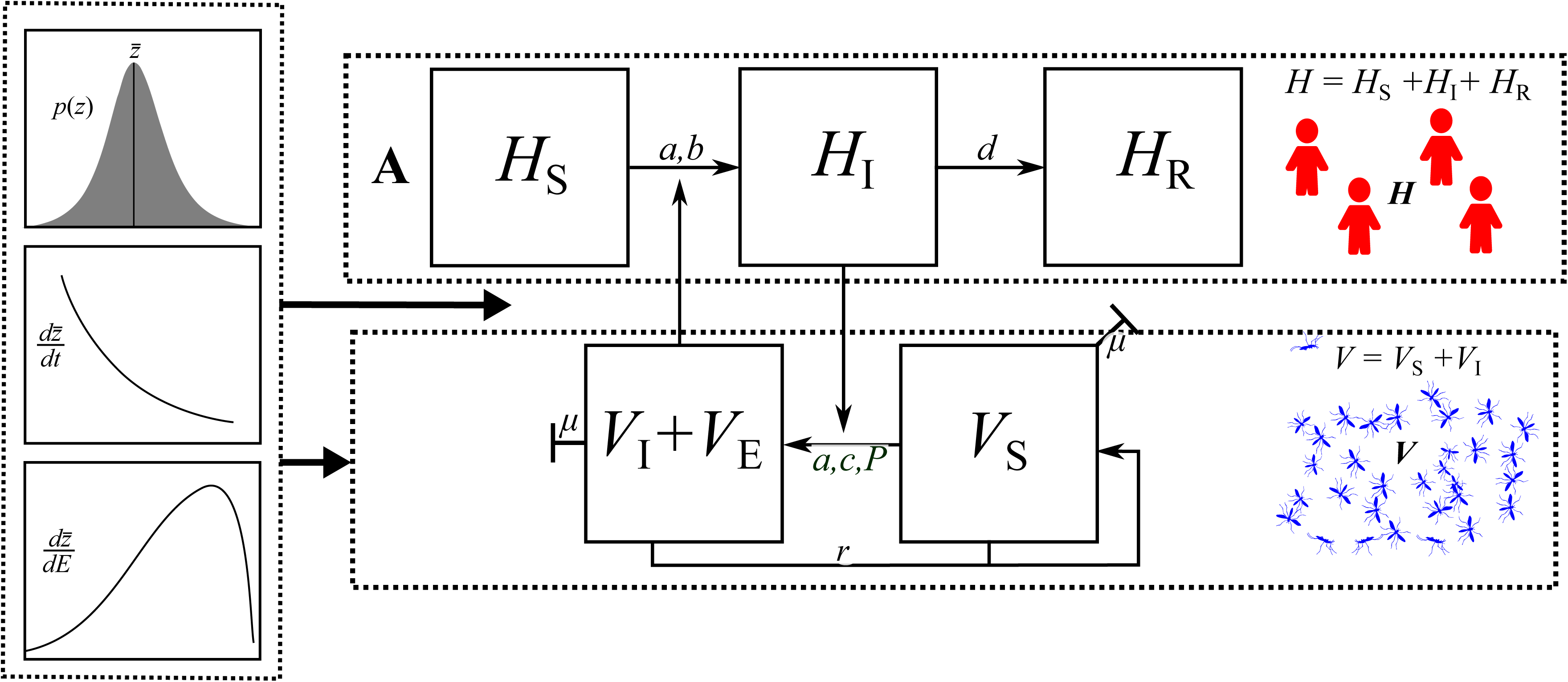
Traits in the classical compartment model with vector population dynamics included (Anderson-May framework). In addition to the host and vector transmission compartments, the Anderson-May framework includes the vector life-history compartments. As an illustration, we extend the classical Ross-MacDonald type models model for malaria (Fig. 2) to include mosquito life history. This introduces additional parameters (vector life stage-specific *μ*’s, fecundity *f*, and stage transition probabilities *p_i_’s*), all of which are directly measurable vector traits (the *μ’s*, *f)*, or which can be derived from underlying traits (the *p_i_’s*). The corresponding equations and parameter definitions are shown in SI Section 2.

### (3) Adding in trait variation

Subsequent studies have incorporated trait variation into the compartment as well as Anderson-May models (Fig. 4). Reiner et al (2013) found that while extensions of the Ross-MacDonald model to incorporate parasite latency in malaria vectors are commonplace, efforts to include variation in vector traits such as heterogeneity in host-vector contact or variation other vector (and host) attributes are rare. Several studies have shown that variation in single traits such as age-specific vector mortality drives changes in the predicted sensitivity of *R*_0_ to vector control (Styer *et al*., 2007; Bellan, 2010; Novoseltsev *et al*., 2012). Incorporating variation in single life history traits associated with infection (McElhany, Real, & Power, 1995; Koella, 2005; Lefèvre & Thomas, 2008) or nutrition (Shapiro *et al*., 2016) has also been found to affect key VBD parameters such as biting rate. In some cases, components of vector ecology have also been added to Ross-MacDonald type models, including environmental drivers (Beck-Johnson *et al*., 2013) or species interactions (Depinay *et al*., 2004; Nakazawa, Yamanaka, & Urano, 2012). While these approaches incorporate trait variation, they don’t necessarily link this to vector population dynamics or link between traits.

**Figure 4.**
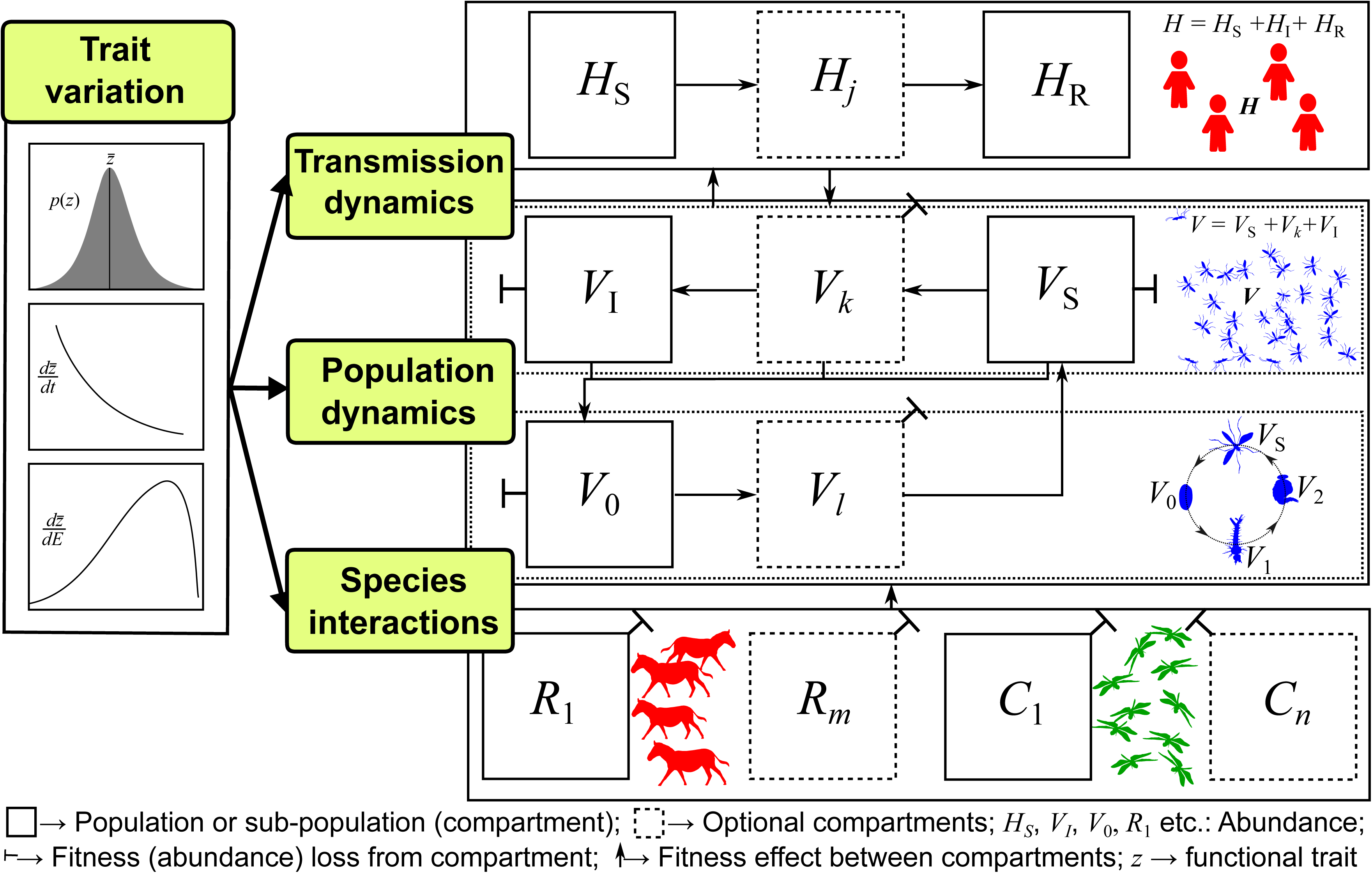
Trait variation in compartment models. In some cases traits and trait variation are used to drive specific components of the classic compartment model (Fig. 2) directly. In some cases traits are individually driven by a common driver (in most examples temperature). Traits are still not related to each other mechanistically, and do not drive the transmission process dynamically through changes in population abundances (e.g., as would happen in the Anderson-May model framework).

Other studies have derived transmission parameters as a function of traits without mechanistically linking traits. For example, Brand, Rock, & Keeling (2016) determined the number of infectious bites delivered by midges by combining the probability of survival through bluetongue virus EIP, age specific-biting rate, and mortality. They demonstrate that determining model parameters with traits can dramatically change the interpretation of *R*_0_ as well as the estimated impact of vector control for bluetongue. However, while the traits drive a common transmission parameter, they are not mechanistically linked to one another. In a similar example, Brady *et al*. (2016) incorporated adult female mosquito blood feeding, egg laying, larval ecology and how these affect vectorial capacity. They found that this increased the relative importance of vector control methods relative to those that specifically targeted adults.

Most recently, trait variation has been incorporated directly into compartment models by incorporating environment driven trait variation. In this case multiple traits are determined by the same driver. For example, Parham & Michael (2010) derived an equation for mean, asymptotic vector population size as a function of traits that are then allowed to be functions of environmental conditions. Mordecai et al. (2013, 2017) built upon this approach to include empirically derived unimodal thermal responses for life-history and transmission related traits. Brand et al. (2017) took a similar approach for allowing biting and EIP parameters to depend on temperature, but modelled the vector population using a statistical approach. All these studies have provided interesting insights into possible effects of trait variation, but they do not mechanistically link traits. We have not found examples of studies that have systematically incorporated trait variation, mechanistic linkages between traits (e.g., co-variation in biting rate and survival), and allowed traits to drive population and transmission dynamics.

## IV. A TRAIT-BASED FRAMEWORK FOR VBD RESEARCH

Here, we present a general trait-based research framework that unifies these previous approaches, and can provides a foundation for systematically tackling the challenge of incorporating trait variation into VBD dynamics. An overview of the framework is given in Fig. 5 (SI section 1, contains a more detailed framework description). In general, a fully trait-based VBD system’s specification must include:

1. **Transmission compartments for each focal host and vector species:** For example, the SIR compartments as presented in the Ross-MacDonald type models (e.g., *H*_S_, *H*_R_) and may have additional host sub-compartments (*H_j_*, where *j ≥* 0) tailored to a particular disease.
2. **Vector life history compartments:** These would include the commonly used infected susceptible vector sub populations (*V*_S,_ *V*_I_), but additionally include the vectors’ juvenile life stage subpopulations, starting at birth (*V*_0_) and followed by immature stages (*V_l_*, where *l ˃* 0). In adult stages, we include the potential for additional stages (*V*_k_) leading to infectious adults (*V*_I_).
3. **Species interaction compartments:** These will depend on the VBD system, but typically at least one consumer-resource interaction would be necessarily incorporated, because all vectors experience significant mortality from predation at one or more life stages. This would require compartments for a resource species consumed by the vector (*R*_1_) and predator species that consume the vector (*C*_1_). In more complex systems, additional consumer and resource compartments (*C_n_, R*_m_) could be added.
4. **Trait Variation**: A suite of trait (e.g., vector mortality, fecundity and biting rates) to parameter mappings that determine fitness effects, and at least a single type of trait variation, such as with an environmental factor (d*z*/d*E*; e.g., temperature) (Fig. 1, 5). As we explain earlier, under this framework “fitness” has a more general meaning in its evolutionary connotation: it is the change in abundance value of any compartment (population) between time steps or spatial locations. Finally, note that a fitness effect in the transmission component (Fig. 5) changes abundances of the host’s infected sub-population through changes in transmission rate (e.g., through biting rate) and reflects a change in fitness of the pathogen’s (population) fitness. While many features of individuals and populations are measureable and may be variable, this framework specifies that these features are only important for transmission when they affect demographic, interaction, or transmission rates that drive population and transmission dynamics. We incorporate three key classes of functional traits; life history, interaction, and transmission traits (Table 1).
5. **Mechanistic Links Between Traits**: Traits should be mechanistically linked. This may be accomplished by explicitly modelling how multiple traits together affect a fitness parameter through shared bio-mechanical and metabolic constraints (see Section VI.1.a).

**Figure 5.**
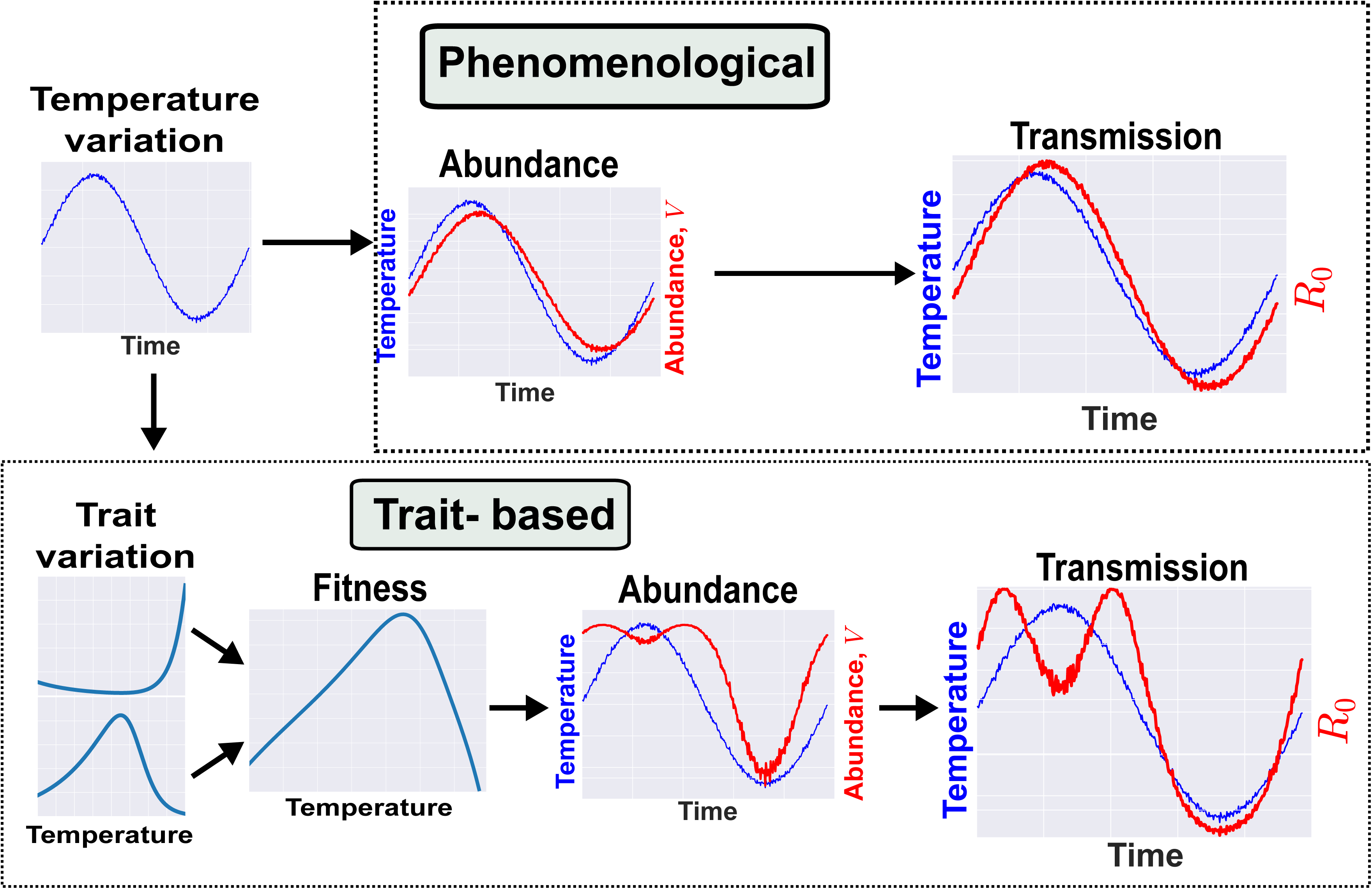
The trait-based framework for vector-borne disease systems. For illustration, we have used a mosquito-borne disease, but this framework can be applied to any VBD with distinct stage or age classes (further details in SI). Arrows between panels represent parameters (potentially with underlying traits) that determine population or transmission dynamics through (absolute) fitness effects. *Transmission dynamics compartments*: Number of Susceptible (S), Infected (I), and Recovered (R) hosts (*H*_S_, *H*_I_, *H*_R_, respectively; additional compartments, *H_j_*, can be added) and number of Susceptible, Infected, and Exposed (E) vectors (*V*_S_, *V*_I_, *V*_E_ respectively). *Vector population dynamics compartments*: number of vector individuals at egg, larval pupal and adult stages (*V*_0_, *V*_1_, *V*_2_, *V*_S_ respectively; additional compartments *V_k_* can be added); *Species interaction compartments*: Abundance of a single resource species (*R*_1_) that is the primary energy source of the vector population (may actually be the host itself, so *R_1_* = *H*_S_), and a single consumer species (*C*_1_) that is the primary source of mortality for the vector population; *Trait variation*: A suite of trait to parameter mappings that determine fitness (e.g., vector mortality, fecundity and biting rates), and a single type of trait variation, such as variation with an environmental factor (d*z*/d*E*; e.g., temperature; see Fig. 3). For developing a mathematical model of such a system, the most common tool would be ordinary differential equation (ODE) systems, as illustrated in SI Section 1.

We do not show explicit linkages between trait variation, consumer-resource, and life history sub-compartments in Fig. 5 because these will vary with VBD system. For example, in the case of most aphid-transmitted diseases, the resource (*R*) and host (*H*) are often the same. For other vectors, such as *Anopheles* mosquitoes, the transmission relevant hosts (*H*) may make up only a proportion of the resources (*R*) that regulate growth and reproduction (e.g. (LoGiudice *et al*., 2003; Donnelly *et al*., 2015)).

## V. AN EXAMPLE: A TRAIT-BASED MODEL OF MOSQUITO ABUNDANCE AND DISEASE TRANSMISSION

We now illustrate how a trait-based approach (Fig. 5) can lead to novel predictions about vector population dynamics and therefore transmission using the effect of temperature on transmission as an example. Temperature is a major source of environment-driven trait variation (Fig. 2), and affects variation in both adult and juvenile traits in vectors. To incorporate trait variation into transmission, we model the effects of temperature-driven life-stage specific trait variation on vector population density, *V*, through the population’s intrinsic growth rate, *r*_m_. Full details of this worked example and the model are provided in SI section 3. Briefly, *r*_m_ is a function of adult peak fecundity (*b*_pk_), age-related fecundity decline rate (*κ*), adult mortality rate (*μ*) and juvenile development time (*α*) and juvenile mortality (*μ*_J_). Variation in each of these traits across temperature is characterized by the thermal performance curve of each trait. By incorporating such environment-driven trait variation into a vector population abundance model we can derive the transmission dynamics over time (Fig. 6).

**Figure 6.**
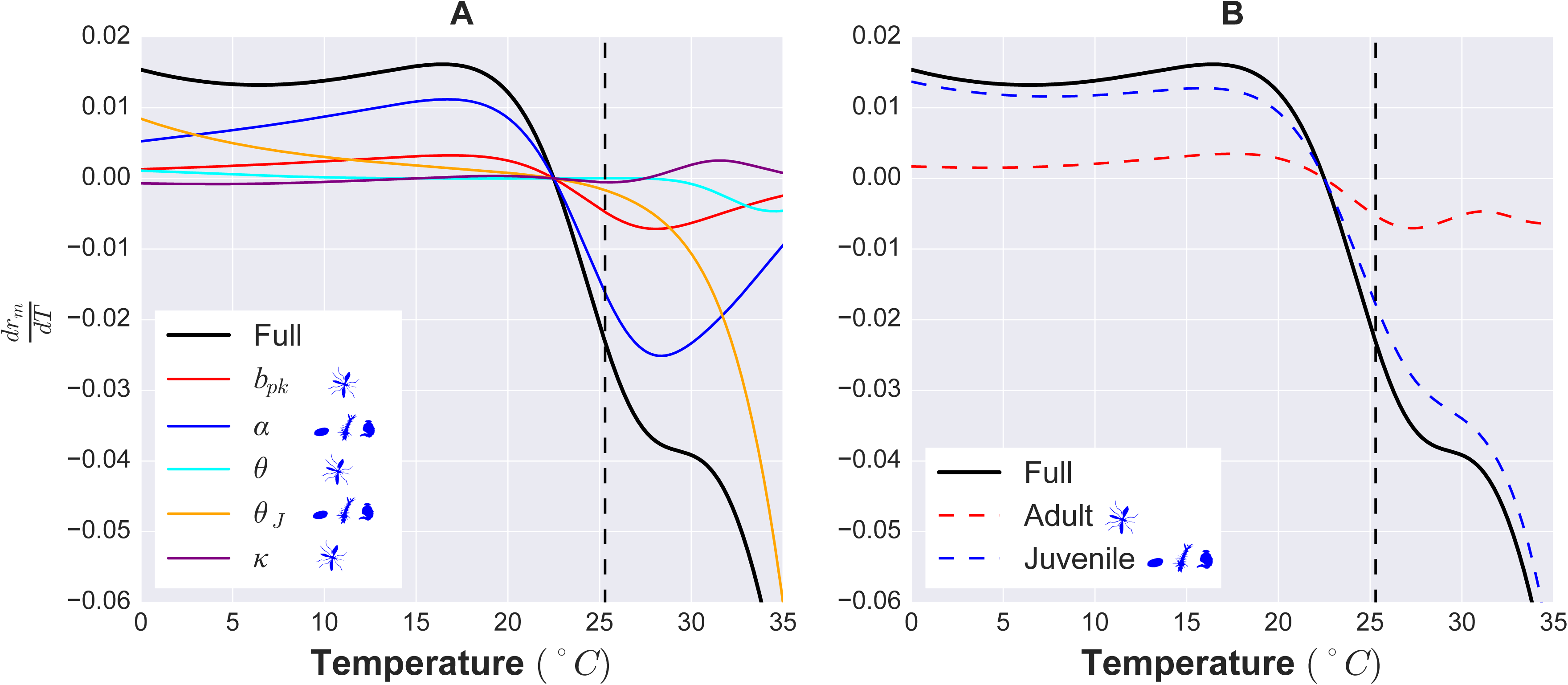
An example trait-based model for malaria transmission. We illustrate here the contrast in models and resulting dynamics produced from phenomenological vs. trait-based approaches. Both models cover a time scale of one year and seek to predict the fluctuation in transmission risk or rate (*R*_0_) during that period. Full details of both models can be found in SI section 3.

We contrast the trait-based approach with a phenomenological one that has been used in previous studies (Hoshi *et al*., 2014; Johnson *et al*., 2018; Ng *et al*., 2018), where abundance (*V*) is directly associated with temperature by fitting a time-series model or where abundance is assumed, a priori, to follow a sinusoidal function (Bacaër & Guernaoui, 2006; Bacaër 2007; Bacaër & Ouifki, 2007; Bacaër & Ait Dads, 2012). This results in a transmission dynamic where *R_0_* tracks temperature variation (Fig. 6), potentially with a time-lag. This result contrasts strongly with that from the trait-based approach that maps traits through parameters to vector population size. Specifically, we observe key differences in predictions of both abundance dynamics and the knock-on effects on transmission: the trait-based model predicts that vector populations will emerge earlier in the year and persist later into the cooler late summer season with a dip in the warmest period of the summer. These differences in *V* feed through to a longer period of annual transmission with an early and late summer peak. The trait-based model predicts a longer transmission season than the phenomenological model, and reveals a decrease in transmission risk in the warmest period. The latter result in particular is surprising given the general “warmer is better” view, but is consistent with the results of Mordecai et al (2013) which incorporated variation (SI section 2.2). Interestingly, when metabolic theory was used to mechanistically model of infection of an endothermic host with an arctic nematode parasite, the continuous spring-to-fall transmission season morphed into two distinct transmission seasons as climate warmed was also observed (Molnár, 2013). The similarity in predicted transmission dynamics across these two very different systems suggests that mechanistically incorporating trait variation can reveal general constraints on VBD systems — in this case, the effect of temperature on VBD dynamics through life-history traits. Additionally, a trait-based approach enables us to determine which traits matter the most for determining thermal sensitivity of transmission dynamics (through trait sensitivity analyses; see next section).

The example we have developed here also illustrates a key theoretical point we raised at the start. If vector traits (eqn 1; assumption 2) change at the same or shorter timescales (here, driven by within-year temperature change) than the rate of pathogen transmission, the classical approach will fail to capture important aspects of contemporary transmission dynamics (Anderson & May, 1981; Heesterbeek & Roberts, 1995; Bacaër, 2007). One approach towards relaxing the constant vector population assumption of classical transmission models has been to build phenomenological models for time varying vector populations, for example by assuming that the vector population oscillates sinusoidally (Heesterbeek & Roberts, 1995; Bacaër, 2007). Although this approach is relatively simple, it cannot quantify or predict how variation in key vector traits or parameters (e.g., *a, b, c, μ*) interact to drive transmission dynamics over time if the relationship between abundance and temperature is non-linear. A trait-based approach provides new mechanistic insights and hypotheses, and predicts non-trivial dynamics that would likely inform public health and control decisions.

### (1) The importance of trait sensitivity analyses

In addition to producing novel predictions, a trait-based approach allows investigation of the degree to which different traits drive transmission dynamics and underlying fitness effects. For example, a local trait sensitivity analysis of the population fitness component of the above trait-based model allows us to investigate the relative importance of juvenile versus adult traits in determining effects of temperature on abundance (and therefore transmission) (Fig. 7).

**Figure 7.**
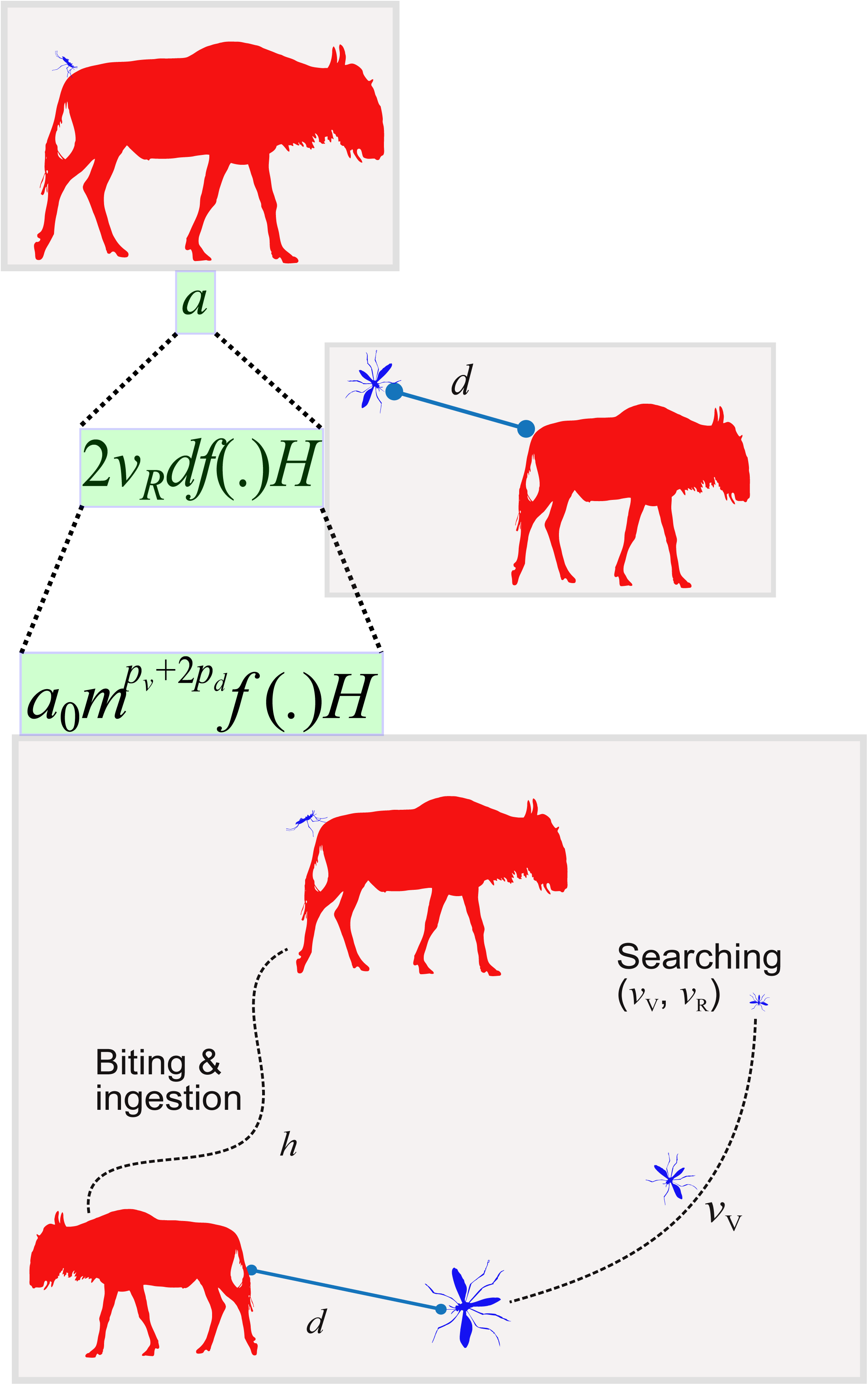
An example trait sensitivity analysis. Analyzing the influence of individual fitness traits on the thermal sensitivity of population fitness of mosquito vectors reveals that juvenile traits have a stronger influence than adult traits in shaping population fitness (*r_m_*) across temperatures-(*b*_pk_), age-related fecundity decline rate (*κ*), adult mortality rate (*μ*) and juvenile development time (*α*) and juvenile mortality (*μ*_J_). The vertical dashed line marks the thermal optimum of fitness. The sensitivity of *r_m_* to the thermal dependence can be assessed by deviation of *dr_m_/dT* from zero. When *dr_m_/dT* is positive this means that population fitness increases at this temperature. **A.** The sensitivity of *r_m_* to the thermal dependence of each parameter can be assessed by deviation of each evaluated partial derivative with respect to each parameter (the differently coloured lines) from zero and compared to the full model (black). **B.** Traits can be combined by life stage, with development rate and juvenile mortality categorised as juvenile traits and adult mortality, fecundity, and rate of loss of fecundity as adult traits. This reveals the relative contribution of the thermal performance curve of each life stage to the thermal performance curve of *r_m_*, again with greater deviation from zero indicating a greater contribution. Full details of the trait sensitivity analysis are in SI Section 4.

This leads to a key insight: juvenile traits are expected to play a major role in determining vector fitness across temperatures, and therefore abundance, and ultimately transmission. In particular, the trait sensitivity analysis adds further insight to *why* the trait-based approach yields very different predictions for population abundance and *R*_0_ compared to a phenomenological approach. The thermal sensitivity of abundance (*V*) and the underlying population fitness (*r_m_*) is driven by the variation of temperature-driven variation in larval stage traits. These predictions and insights provide quantitative targets for validation using field data. Sensitivity analyses of transmission measures with respect to traits also allow key traits to be identified, guiding further empirical and theoretical work on the contributions of traits to VBD system dynamics.

## VI. A TRAIT-BASED FRAMEWORK ROADMAP

Our relatively simple worked example in the previous section gives an idea of the steps one would need to take in order to use a trait-based approach. A full trait-based study would involve making the following links:

1. Trait→Parameter
2. Trait-Variation→Fitness Variation
3. Fitness→Population Dynamics
4. Population Dynamics→Transmission Dynamics

Research programs may focus on all or a portion of the following four sequential components or steps. In our worked example, we only attempt to tackle step 2 and 3. Each step requires a relationship or mapping (→) to be quantified through empirical studies coupled with mathematical modelling. To be clear, useful progress and insights can be gained at each step. We now explain each of these steps, followed by an example.

### (1) Trait→Parameter

A key component of any trait-based framework is the mapping of trait values onto mathematical VBD model parameters (Fig. 5). In this framework a parameter represents a process that would have a fitness effect within the model (e.g., it can change the abundance of one of the model compartments). For example, mortality rate decreases population abundance (*V*_S_) and fecundity increases abundance (*V*_S_). The trait/parameter delineation is not exclusive. Mortality rate and fecundity could also be interpreted as traits because they are directly measurable. In contrast, vectorial capacity is not a trait as it is a derived measure and cannot be directly measured. The distinction between “trait” and “parameter” is a matter of context; when a quantity is being used to drive another parameter it is a trait and when is directly affecting fitness it is a parameter. Deconstruction of parameters into their underlying traits bounds the parameter’s feasible range (parameter space), and also reveals how different parameters are linked, and therefore interact or trade-off with one another to impact compartment fitness (Charnov, 1993) and ultimately, transmission dynamics.

Biomechanical and metabolic approaches that link physical and performance traits could be very useful for tackling this challenge (Brown *et al*., 2004; McGill *et al*., 2006; Amarasekare & Savage, 2012; Pawar, Dell, & Savage, 2015a). These approaches use a common currency (energy) to map parameters onto traits. This inevitably establishes mechanistic links between traits. For example, body size drives not just adult vector biting rate, but also its fecundity and mortality rates. Recent advances in metabolic modelling offer an opportunity to determine encounter rate parameters between vectors and hosts (Dell, Pawar, & Savage, 2011; Pawar, Dell, & Savage, 2012; Dell, Pawar, & Savage, 2014; Gilbert *et al*., 2014; Pawar *et al*., 2015a; Rizzuto, Carbone, & Pawar, 2018) (see example below) and even capture within-host parasite dynamics (Kirk *et al*., 2018). Empirical studies that validate trait-fitness models for specific vectors and VBD systems are of equal importance. Ideally, such studies should measure multiple traits within individuals (for example, vector competence, fecundity, and survival) and how they relate to individual fitness (Ohm *et al*., 2016).

#### (a) A Worked Example: Deconstruction of Biting Rate

To illustrate Trait***→*** Parameter mapping, we use recent advances in mechanistic, metabolic trait-based theory for species interactions (Pawar *et al*., 2012; Gibert *et al*., 2015; Pawar *et al*., 2015a) to develop a mathematical model that deconstructs biting rate (a parameter in the model) into its underlying functional traits and show how it is ultimately linked to the vector’s adult body size. Heterogeneity in biting behaviour is considered important for determining transmission dynamics (Dye & Hasibeder, 1986; Smith *et al*., 2014). This key transmission parameter in all VBD systems, that also underpins vector fitness, can be derived from body size (a physical trait) (Fig. 8, SI section 1.2)(De Leo & Dobson, 1996). Adult vector size is a key physical trait as it determines many functional traits, and also itself responds strongly and predictably to external factors such as environmental temperature and resource availability (e.g., larval nutrition).

**Figure 8.**
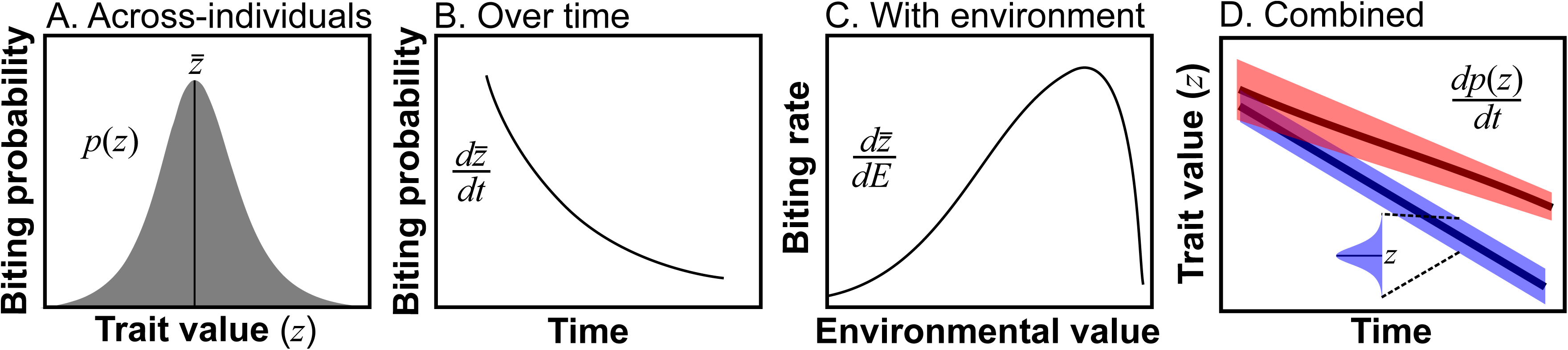
Illustration of a Trait-Parameter mapping. We develop a mechanistic, trait-based model for biting rate (a key transmission parameter; eqn. 1) of vectors by deconstructing it into component functional traits (shown above). Biting rate (*a*) can be decomposed into relative velocity (*v_r_*), distance between vectors and hosts (*d*), risk (*f(.)*) and host density (*H*). These parameters can be further decomposed so that f(.) is determined by handling time (*h*) and the mass-specific velocities of the vector and host (*V_V_*, *V_R_*) as in eq. 2. The model derivation is in SI section 1.2.

Consider a flying vector such as a mosquito, fly, or midge, seeking larger, warm-blooded hosts (e.g., a terrestrial mammal, which is also the energy resource for the vector). A general field encounter rate model for such vector-host interactions in two-dimensional Euclidean space (2*D*) is (Pawar *et al*., 2012),

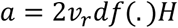

Here, *a* is biting rate (bites × time^−1^) for a given host density *H* (individuals × area^−1^), *v*_r_ is relative velocity between the vector and host, *d* is the reaction distance (minimum distance at which the vector detects and reacts to host), and *f*(.) is the host “risk” function that determines the (biting) functional response of the vector. That is, *f*(.) encapsulates the constraints on the vector’s handling time (*h*) (the post-encounter phase in Fig. 8), which would be determined by its ability to land on and complete the biting action successfully. Assuming each encounter between vector and host results in one bite, we can derive biting rate as a function of body size (as physical trait) two key functional traits (vector’s flight velocity and reaction distance) (details in SI section 1.2):

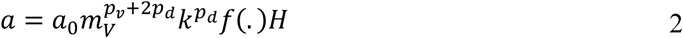

where *a*_0_ is a normalization constant that also includes the effect of environmental temperature, *m*_V_ is the vector’s body mass, *p*_v_ and *p*_d_ are scaling exponents for the size-dependence of vector flight velocity and reaction distance (respectively) *k* is the vector-host body mass-ratio (a measure of their relative size difference). In the case where hosts are relatively rare, *f*(.) = 1, but in the case where hosts are common (e.g., mosquito populations are often localized near human villages), it can be replaced with a more complex function that includes the effect of host handling time (physical limits on vector’s biting & ingestion rate; Fig. 2), which itself is likely to be vector size and temperature dependent (Dell *et al*., 2011; Pawar *et al*., 2012).

Breaking down biting rate into its component traits recovers known patterns: that smaller flying vectors will have a lower mean or maximum biting rate relative to larger ones (e.g., midges vs. mosquitoes), that biting rates will be lower for walking vectors compared to flying ones (e.g., ticks vs. mosquitoes), and explains how temperature (through the scaling constants *a*_0_) affects biting rate (e.g., (Dell *et al*., 2011; Mordecai *et al*., 2013; Dell *et al*., 2014), providing a mechanistic basis for determining the effect of temperature on vector fitness as well as vectorial capacity. This Trait→Parameter mapping approach can be modified to accommodate vectors with different moving or sensory modes (e.g., ticks vs. mosquitoes), and for the same host density (say, humans).

### (2) Trait-Variation→Fitness

A population’s fitness is essentially a distribution of fitness values across its individuals. Therefore, the second key step in a trait-based VBD framework is to use Trait-Parameter mappings to quantify how variation in functional traits generates variation in vector fitness and transmission. That is, after identifying key functional traits and building trait–parameter models, the effect of trait variation on the parameter, and eventually, fitness distribution in each class of compartment needs to be quantified. The same trait can affect the fitness of more than one compartment. For example, variation in biting rate or intrinsic mortality would affect both vector population fitness (e.g., *r*_max_) and transmission fitness (e.g., *R*_0_). Incorporating or mapping any of the above three types of trait variation (Fig. 1) onto vector population or transmission parameter requires the re-definition of the parameters as functions (e.g., *p*(*z*), *dz*/*dt*, *dz*/*dE*; Fig. 3), which are better constructed mechanistically using Trait-Parameter mappings. In our worked example, we explicitly derive population fitness using environment-driven traits.

### (3) Fitness→Population Dynamics

The third step is to quantify how trait variation determines vector population abundance or dynamics over time through effects on fitness. This requires the construction of stage-structured population dynamic models (Fig. 5). There are two key challenges here. The first is quantification of the impact of trait variation (Fig. 3) on the vector’s stage-structured population dynamics. One approach for doing so is trait-driver theory, which provides methods inherited from quantitative genetics to study how trait variation drives abundance dynamics (Norberg *et al*., 2001; Webb *et al*., 2010; Enquist *et al*., 2015). This challenge may be made more difficult because most vectors have stage-structured life cycles underlying population dynamics, which may have transgenerational effects (e.g., maternal effects) or carryover effects across life stages (Lorenz & Koella, 2011). Initial progress can be made by simply measuring trait variation and “plugging” it into population dynamics models (through Trait-Parameter models) with variation to allow deviations from the standard exponential assumptions (for example see (Brand *et al*., 2016)) to determine how sensitive abundance is to trait variation. In the worked example, we mapped trait-based fitness effects onto vector abundance using one type of trait variation (environment-driven, specifically temperature). Because body-size is a key physical trait that also changes with life stage (over time), integral projection models may be another way to incorporate traits into VBDs (Coulson, 2012; Rees, Childs, & Ellner, 2014; Metcalf *et al*., 2016).

There is increasing interest in incorporating species interactions into VBD transmission dynamics (Keesing *et al*., 2010). This is an area of ongoing investigation not just in VBD research, but in ecology in general. Species interactions impact life history traits, especially fecundity and mortality, by shifting them from the baseline, interaction-independent values (Roux *et al*., 2015). For example, in consumer-resource interactions, fecundity increases with availability of the vector’s resources (vector-resource or vector-host interaction), and mortality with the vector’s consumers (vector-predator interaction). Tackling this challenge will require mathematical models (and complementary empirical studies) that tractably include the impacts of species interactions on baseline (or idealized) life history (demographic) parameters (typically measured in the laboratory), especially fecundity and mortality. One relatively simple way to make progress in this direction is to re-define the otherwise idealized life history parameter (e.g., development rate, fecundity, and mortality) functions to include losses or gains due to species interactions. This approach could circumvent the additional complexity of explicitly adding consumer-resource dynamics to vector population and transmission dynamics.

### (4) Population Dynamics→Transmission Dynamics

The final step is to quantify transmission dynamics using trait-based vector population dynamics and resulting vector-host interaction rates (Fig. 5). To achieve this, two key challenges on the theoretical front need to be tackled. First, how trait variation determines the timescale of fluctuations in vector population sizes and vector-host interaction traits relative to the timescale of transmission dynamics needs to be modeled and validated. The trait-based approach, by deriving the timescales of population fluctuations mechanistically, would “naturally” determine whether and when the separation of the timescales of population and transmission dynamics, implicit in classical (compartment-type) VBD models, is valid (see example below). Second, the appropriate level of biological complexity (essentially, number of parameters or traits) needed to capture the effect of trait variation through population dynamics on transmission dynamics needs to be determined. For example, trait variation in both juveniles and adults may need to be incorporated simultaneously into transmission models (Fig 7). In addition to vector traits that affect its population dynamics, models and data on traits that determine the ability of the vector to transmit a pathogen (e.g., *P* and *bc* in eqn. 1) and interact with the host (e.g., *a* in eqn. 1) will be needed. In many cases, these transmission-relevant traits will be the same as those determining fitness. For example, both, encounter rate with host and with the vector’s resources (or predators) are determined by body size and velocity (Fig. 8). Indeed, the host is the primary or sole resource in many vectors (e.g., aphids) which links transmission parameters directly to the vector’s fitness though biting and feeding rate.

## VII. KEY CHALLENGES

The four components for building trait-based models that we have outlined above will each face four challenges to differing degrees; *data*: how to prioritize experiments and report data; *parameterisation*: how to link model components to empirical data; *validation*: how to test the trait – fitness, trait variation – fitness, population dynamics, and transmission models; *selection*: how to pick the most parsimonious model.

### (1) Data

Experiments to quantify vector traits involve taking individual measures on small insects that are often logistically difficult to work with due to ethical and biosafety concerns. As we have described above, theoretical work can help to prioritize which traits we should expect to have the greatest impact on fitness and transmission. New data collection efforts are currently underway in several disease systems, but the limited accessibility of data that have already been collected is a major issue that needs to be addressed. It is critical to get as much information as possible from these time-and labor-intensive experiments by reporting data at the most disaggregated level possible. Many studies report point estimates of trait means/demographic rates across populations, across lifespans, and over environmental gradients rather than presenting distributions or individual values. Individual-level data are critical for accurately estimating within-population variation in trait values. Beyond individual studies, consolidating datasets with individual measurements into common formats would allow for the identification of gaps and coordinated data collection efforts to specifically target the traits and conditions that are data-poor. We have recently launched a hub for storing and accessing vector trait data (www.vectorbyte.org) and a platform for coordinating data collection efforts (www.vectorbite.org). Current data can be used to parameterise mechanistic models, including trait distributions that accurately represent uncertainty and variation within populations. This would allow us to identify which traits and types of variation in these traits are not well characterized and may play an important role in transmission.

### (2) Parameterisation

Accurately quantifying the distribution of trait values within populations is a major barrier to developing and refining Trait → Parameter and Trait Variation → Population Fitness models. First, better identification of how individual measurements link to modeled quantities needs to be more consistently addressed. For example, many models assume a single fixed value of the extrinsic incubation period (EIP – the time it takes for an exposed vector to become infectious). However, studies may report any number of variations in EIP – such as EIP_50_ (time to 50% of vectors being infectious) of EIP_90_ (time to 90% infectious) (Ohm *et al*., 2018). Correctly identifying and incorporating empirical measurements from the literature to match with the meaning of the trait or parameters or transformations is necessary. Then the variation and uncertainty in the traits must be quantified. This involves the use of statistical methods for separating true biological variation from measurement error, and thereby quantifying uncertainty in estimates of trait values (e.g., the mean or variance of a trait). Using sophisticated approaches that allow quantification and propagation of uncertainty (e.g., from Trait→Parameter to population dynamics modelling), such as Bayesian inference, with or without prior information, will be key here (Clark, 2007; Johnson *et al*., 2015). In addition, parameter sensitivity analyses in trait-based models (Fig. 7) are crucial, and can provide biological insights into the traits driving variation in transmission (Johnson *et al*., 2015).

### (3) Model Validation and Selection

A fundamental goal of trait-based VBD research should be to determine the conditions under which vector traits drive significant variation in realized transmission rate. This requires validation of models at each level or compartment of the framework (ideally) with data on the spatial or temporal distribution of traits/drivers as predictors. In contrast to inference or calibration of a model, validation is the process of assessing how well a parameterised model can replicate data that was not used for parameter inference/calibration (i.e., out-of-sample prediction) (Hooten & Hobbs, 2015). For example, Trait→Parameter models need to be validated with data on population growth rates, the subsequent population dynamics models need to be validated with data on abundance variation over space or time, and the transmission models need to be validated using disease incidence data over space and time. However, when the data available for validation are very different in nature from the data used for model fitting or model outputs do not correspond to easily measured quantities, it can be difficult to determine when a model captures sufficient detail to adequately represent reality. Efforts to develop hierarchical model validation methods for the types of dynamic, trait-based models described here are ongoing (e.g., (LaDeau *et al*., 2011; Johnson, Pecquerie, & Nisbet, 2013; Johnson *et al*., 2014; Sun, Lee, & Hoeting, 2015). These will include Bayesian methods, which allow quantification of uncertainty and the incorporation of prior data (Clark, 2007; Hooten & Hobbs, 2015). Not only do the statistical methods for (VBD) dynamical systems need to be refined, but as they are developed it is important that these methods are made accessible for non-statisticians doing research in this area. This requires training a new generation of researchers in both the new modeling techniques (so they can develop models that include details such as behaviour) as well as statistical techniques appropriate for parameterizing and validating the models as they are developed.

It is inevitable and useful that multiple models will be built to address the same question within any of the compartments of a trait-based framework (Johnson & Omland, 2004). For example, there are multiple ways to predict fitness from underlying life history traits (Amarasekare & Coutinho, 2013). Comparing the predictions from multiple models allows us to identify which models are most should make validation data sets publically available and accessible, and standardized metrics of goodness-of-fit or similar should be reported for all models against validation sets. Second, publishing the code used to generate model outputs as a standard practice would facilitate comparing transmission models of all types—not just those incorporating trait variation. These steps would enable model comparison and multi-model ensembles to be used for future predictions.

## VIII. WHY NOW?

This is the ideal time to undertake the development of fully trait-based approaches in VBD research. Recent public health crises have spurred government agencies to support the collection of large amounts of data on these systems. This, combined with innovations in empirical data collection and sharing, means that the necessary data for parameterizing and validating trait-based models are now becoming available. At the same time, the movement towards reproducible research has resulted in open models and computational approaches that allow more direct comparison of models and easier expansion of existing methods.

At the same time, the broader field of trait-based research is maturing across ecological systems, with both the theory and experimental methods growing apace (McGill *et al*., 2006; Pawar, Woodward, & Dell, 2015b). This provides VBD trait-based approaches with a solid foundation. There are many other areas of ecology that are currently striving to mechanistically incorporate trait variation to understand emergent (especially community and ecosystem level) processes. Methods developed for VBDs would have utility in solving problems in these other systems, such as predicting changes in ecosystem services like primary production (Díaz *et al*., 2007; Blanchard *et al*., 2012).

Building a fully trait-based approach to modeling VBD dynamics is not the “quick and easy path” (Kershner & Lucas, 1980). It is data-hungry and requires extensive efforts to building models that integrate knowledge about processes from the individual to populations and beyond. However, we argue that in comparison to phenomenological approaches (e.g., regressions or other correlative approaches) taking a more mechanistic approach, in general, provides a better way to extrapolate dynamics across time or space (Bayarri *et al*., 2009). This kind of predictive power is becoming increasingly critical as these pathogens continue to emerge and expand their ranges in the context of a globalized and rapidly changing environment. Beyond that, incorporating traits with this bottom up approach should allow us to better understand *why* we see the patterns of transmission that we do, and potentially inform efforts at control. By explicitly modelling the fitness of a given trait and its effect on population dynamics and fitness, trait-based approaches could be used to incorporate trait evolution into transmission models. For instance, although we have focused on traits that directly affect vector population fitness in idealized conditions, the addition of other traits that are mediated by human intervention, such as insecticide resistance, is also possible within this framework. The evolution of insecticide resistance is arguably the largest challenge to sustainable management of vector-borne diseases. A trait-based approach has the potential to better understand the implication of both current (e.g., chemical pesticides) and future control measures (e.g., genetically altered vectors) that inherently alter traits, while suggesting innovative and nuanced ways to apply control in a way that to anticipates changes driven by the inherent complexities of these systems.

## IX. CONCLUSION

1. Mounting evidence shows that vector functional traits vary over lifespan, within populations, and across environmental gradients. This variation in key parameters is expected to alter disease transmission dynamics.
2. Mathematical models of Vector-Borne Disease (VBD) dynamics to date have not mechanistically incorporated trait variation, and are therefore unable to capture changes in transmission dynamics that may emerge from different types and magnitudes of trait variation, the correlations between traits, and resulting vector population dynamics.
3. A fully trait-based VBD approach would involve explicitly modelling trait variation and its effect on transmission, vector life history, species interactions and also allowing feedback between traits. This requires mechanistic Trait-Parameter mapping, and the incorporation of trait variation (and co-variation) into the fitness of vector population and disease transmission compartments.
4. Using a trait-based VBD approach can lead to novel predictions for transmission dynamics, and allows analyses of the relative importance of different traits in transmission dynamics. Incorporating trait variation into VBD transmission models will enhance our ability to predict transmission dynamics at longer temporal and larger spatial scales.
5. Increasing research efforts in the development of trait-based theory and data, combined with increasing investment in vector population surveillance makes this an ideal time for the community to systematically and strategically incorporate trait variation into VBD research.

## Acknowledgements

We would like to thank the members of the VectorBiTE RCN (www.vectorbite.org) for their energy and stimulating discussions around these ideas. This work was funded by NIH grant 1R01AI122284-01 and BBSRC grant BB/N013573/1 as part of the joint [NIH-NSF-USDA-BBSRC] Ecology and Evolution of Infectious Diseases program. EAM and LRJ were funded by NSF grant DEB-1518681.

## Author Contributions

LJC, SP, LRJ, EM and PJH conceived the study. LJC and SP wrote the manuscript with inputs from MBT, AGP, MB, SLL, MAJ, LRJ, EM, and PJH. SP, LRJ, TS, and FEM developed the mathematical models and worked examples.

## References

Agashe, D. (2009) The stabilizing effect of intraspecific genetic variation on population dynamics in novel and ancestral habitats. American Naturalist 174, 255–267.

Alto, B.W., Lounibos, L.P. & Juliano, S.A. (2003) Age-dependent bloodfeeding of *Aedes aegypti* and Aedes albopictus on artificial and living hosts. Journal of the American Mosquito Control Association 19, 347–352.

Amarasekare, P. & Savage, V. (2012) A framework for elucidating the temperature dependence of fitness. The American Naturalist 179, 178–191.

Amarasekare, P. & Coutinho, R.M. (2013) The intrinsic growth rate as a predictor of population viability under climate warming. Journal of Animal Ecology 82, 1240–1253.

Anderson, R.M. & May, R.M. (1979) Population biology of infectious diseases: Part I. Nature 280, 361–367.

Anderson, R.M. & May, R.M. (1981) The population dynamics of microparasites and their invertebrate hosts. Philosophical Transactions of the Royal Society of London B 291, 451–524.

Bacaër, N. (2007) Approximation of the basic reproduction number R0 for vector-borne diseases with a periodic vector population. Bulletin of Mathematical Biology 69, 1067–1091.

Bacaër, N. & Ait Dads, E. (2012) On the biological interpretation of a definition for the parameter R0 in periodic population models. Journal of Mathematical Biology 65, 601–621.

Bacaër, N. & Guernaoui, S. (2006) The epidemic threshold of vector-borne diseases with seasonality. Journal of Mathematical Biology 53, 421–436.

Bacaër, N. & Ouifki, R. (2007) Growth rate and basic reproduction number for population models with a simple periodic factor. Mathematical Biosciences 210, 647–658.

Bayarri, M., Calder, E., Lunagomez, S., Pitman, E., Berger, J., Dalbey, K., Patra, A., Spiller, E. & Wolpert, R. (2009) Using statistical and computer models to quantify volcanic hazards. Technometrics 51, 401–413.

Bayoh, M.N. & Lindsay, S.W. (2003) Effect of temperature on the development of the aquatic stages of *Anopheles gambiae* sensu stricto (Diptera: Culicidae). Bulletin of Entomological Research 93, 375–381.

Beck-Johnson, L.M., Nelson, W.A., Paaijmans, K.P., Read, A.F., Thomas, M.B. & Bjørnstad, O.N. (2013) The effect of temperature on *Anopheles* mosquito population dynamics and the potential for malaria transmission. PLoS ONE 8, e79276.

Bellan, S. (2010) The importance of age dependent mortality and the extrinsic incubation period in models of mosquito-borne disease transmission and control. PLoS One 5, e10165.

Blanchard, J.L., Jennings, S., Holmes, R., Harle, J., Merino, G., Allen, J.I., Holt, J., Dulvy, N.K. & Barange, M. (2012) Potential consequences of climate change for primary production and fish production in large marine ecosystems. Philosophical Transactions of the Royal Society of London B 367, 2979–2989.

Bohbot, J., Durand, N., Vinyard, B. & Dickens, J. (2013) Functional development of the octenol response in *Aedes aegypti*. Frontiers in Physiology.

Bolnick, D.I., Amarasekare, P., Araújo, M.S., Bürger, R., Levine, J.M., Novak, M., Rudolf, V.H.W., Schreiber, S.J., Urban, M.C. & Vasseur, D. (2011) Why intraspecific trait variation matters in community ecology. Trends in ecology & evolution 26, 183–192.

Brady, O.J., Godfray, H.C.J., Tatem, A.J., Gething, P.W., Cohen, J.M., Mckenzie, F.E., Perkins, T.A., Reiner, R.C., Tusting, L.S., Sinka, M.E., Moyes, C.L., Eckhoff, P.A., Scott, T.W., Lindsay, S.W., Hay, S.I., ET AL. (2016) Vectorial capacity and vector control: reconsidering sensitivity to parameters for malaria elimination. Transactions of the Royal Society of Tropical Medicine and Hygiene 110, 107–117.

Brand, S., Rock, K. & Keeling, M. (2016) The interaction between vector life history and short vector life in vector-borne disease control. PLoS computational biology 12, e1004837.

Brown, J.H., Gillooly, J.F., Allen, A.P., Savage, V.M. & West, G.B. (2004) Toward a metabolic theory of ecology. Ecology 85, 1771–1789.

Caraco, T., Glavanakov, S., Chen, G., Flaherty, J.E., Ohsumi, T.K. & Szymanski, B.K. (2002) Stage-structured infection transmission and a spatial epidemic: a model for Lyme disease. The American Naturalist 160, 348–359.

CDC (2016) Lyme disease: data and statistics. www.cdc.gov/lyme/stats/ [accessed 31 May 2016].

Charnov, E.L. (1993) Life History Invariants: Some Explorations of Symmetry in Evolutionary Ecology. Oxford University Press.

Christensen, B.M., Li, J., Chen, C.-C. & Nappi, A.J. (2005) Melanization immune responses in mosquito vectors. Trends in Parasitology 21, 192–199.

Ciota, A.T., Matacchiero, A.C., Kilpatrick, A.M. & Kramer, L.D. (2014) The effect of temperature on life history traits of *Culex* mosquitoes. Journal of medical entomology 51, 55–62.

Clark, J. (2007) Models for Ecological Data: An Introduction. Princeton University Press, Princeton N.J.

Clements, A.N. & Paterson, G.D. (1981) The analysis of mortality and survival rates in wild populations of mosquitoes. Journal of Applied Ecology 18, 373–399.

Coletta-Filho, H.D., Daugherty, M.P., Ferreira, C. & Lopes, J.R. (2014) Temporal progression of *Candidatus Liberibacter asiaticus* infection in citrus and acquisition efficiency by *Diaphorina citri*. Phytopathology 104, 416–421.

Costa, E.A.P. DE A., Santos, E.M. DE M., Correia, J.C. & Albuquerque, C.M.R. DE (2010) Impact of small variations in temperature and humidity on the reproductive activity and survival of *Aedes aegypti* (Diptera, Culicidae). Revista Brasileira de Entomologia 54, 488–493.

Coulson, T. (2012) Integral projections models, their construction and use in posing hypotheses in ecology. Oikos 121, 1337–1350.

Crutsinger, G., Collins, M., Fordyce, J., Gompert, Z., Nice, C. & Sanders, N. (2006) Plant genotypic diversity predicts community structure and governs an ecosystem process. Science 313, 966–968.

De Leo, G.A. & Dobson, A.P. (1996) Allometry and simple epidemic models for microparasites. Nature 379, 720–722.

De Xue, R., Edman, J.D. & Scott, T.W. (1995) Age and body size effects on blood meal size and multiple blood feeding by *Aedes aegypti* (Diptera: Culicidae). Journal of Medical Entomology 32, 471–474.

Delatte, H., Gimonneau, G., Triboire, A. & Fontenille, D. (2009) Influence of temperature on immature development, survival, longevity, fecundity, and gonotrophic cycles of *Aedes albopictus*, vector of chikungunya and dengue in the Indian Ocean. Journal of Medical Entomology 46, 33–41.

Dell, A.I., Pawar, S. & Savage, V.M. (2011) Systematic variation in the temperature dependence of physiological and ecological traits. Proceedings of the National Academy of Sciences 108, 10591–10596.

Dell, A.I., Pawar, S. & Savage, V.M. (2014) Temperature dependence of trophic interactions are driven by asymmetry of species responses and foraging strategy. Journal of Animal Ecology 83, 70–84.

Den Otter, C., Tchicaya, T. & Schutte, M. (2008) Effect of age, sex and hunger on the antennal olfatory sensitivity of tsetse flies. Physiological Entomology 16, 173–182.

Depinay, J.M.O., Mbogo, C.M., Killeen, G., Knols, B., Beier, J., Carlson, J., Dushoff, J., Billingsley, P., Mwambi, H., Githure, J. & Others (2004) A simulation model of African *Anopheles ecology* and population dynamics for the analysis of malaria transmission. Malaria Journal 3, 29.

Díaz, S., Lavorel, S., De Bello, F., Quétier, F., Grigulis, K. & Robson, T.M. (2007) Incorporating plant functional diversity effects in ecosystem service assessments. Proceedings of the National Academy of Sciences 104, 20684–20689.

Dick, O.B., San Martín, J.L., Montoya, R.H., Del Diego, J., Zambrano, B. & Dayan, G.H. (2012) The history of dengue outbreaks in the Americas. The American Journal of Tropical Medicine and Hygiene 87, 584–593.

Dohm, D.J., O’guinn, M.L. & Turell, M.J. (2002) Effect of environmental temperature on the ability of *Culex pipiens* (Diptera: Culicidae) to transmit West Nile Virus. Journal of Medical Entomology 39, 221–225.

Donnelly, B., Berrang-Ford, L., Ross, N.A. & Michel, P. (2015) A systematic, realist review of zooprophylaxis for malaria control. Malaria Journal 14, 313.

Dye, C. & Hasibeder, G. (1986) Population dynamics of mosquito-borne disease: effects of flies which bite some people more frequently than others. Transactions of the Royal Society of Tropical Medicine and Hygiene 80, 69–77.

Enquist, B.J., Norberg, J., Bonser, S.P., Violle, C., Webb, C.T., Henderson, A., Sloat, L.L. & Savage, V.M. (2015) Scaling from traits to ecosystems: developing a general trait driver theory via integrating trait-based and metabolic scaling theories. In Advances in Ecological Research pp. 249–318. Elsevier.

Faria, N.R., Azevedo, R. Do S. Da S., Kraemer, M.U.G., Souza, R., Cunha, M.S., Hill, S.C., Thézé, J., Bonsall, M.B., Bowden, T.A., Rissanen, I., Rocco, I.M., Nogueira, J.S., Maeda, A.Y., Vasami, F.G. Da S., Macedo, F.L. De L., ET AL. (2016) Zika virus in the Americas: Early epidemiological and genetic findings. Science 352, 345–349.

Getz, W.M., Marshall, C.R., Carlson, C.J., Giuggioli, L., Ryan, S.J., Romañach, S.S., Boettiger, C., Chamberlain, S.D., Larsen, L., D’odorico, P. & O’sullivan, D. (2018) Making ecological models adequate. Ecology Letters 21, 153–166.

Gibert, J.P., Dell, A.I., Delong, J.P. & Pawar, S. (2015) Scaling-up trait variation from individuals to ecosystems. In Advances in Ecological Research (ed G.W. AND A.I.D. Samraat Pawar), pp. 1–17. Academic Press.

Gilbert, B., Tunney, T.D., Mccann, K.S., Delong, J.P., Vasseur, D.A., Savage, V., Shurin, J.B., Dell, A.I., Barton, B.T. & Harley, C.D. (2014) A bioenergetic framework for the temperature dependence of trophic interactions. Ecology Letters 17, 902–914.

Harrington, L.C., Buonaccorsi, J.P., Edman, J.D., Costero, A., Kittayapong, P., Clark, G.G. & Scott, T.W. (2001) Analysis of survival of young and old *Aedes aegypti* (Diptera: Culicidae) from Puerto Rico and Thailand. Journal of medical entomology 38, 537–547.

Harrington, L.C., Françoisevermeylen, Null, Jones, J.J., Kitthawee, S., Sithiprasasna, R., Edman, J.D. & Scott, T.W. (2008) Age-dependent survival of the dengue vector *Aedes aegypti* (Diptera: Culicidae) demonstrated by simultaneous release-recapture of different age cohorts. Journal of Medical Entomology 45, 307–313.

Heesterbeek, J. & Roberts, M. (1995) Threshold quantities for infectious diseases in periodic environments. Journal of biological systems 3, 779–787.

Hillyer, J.F., Schmidt, S.L., Fuchs, J.F., Boyle, J.P. & Christensen, B.M. (2005) Age-associated mortality in immune challenged mosquitoes (*Aedes aegypti*) correlates with a decrease in haemocyte numbers. Cellular Microbiology 7, 39–51.

Hooten, M. & Hobbs, N. (2015) A guide to Bayesian model selection for ecologists. Ecological Monographs 85, 3–28.

Hoshi, T., Higa, Y. & Chaves, L.F. (2014) *Uranotaenia novobscura ryukyuana* (Diptera: Culicidae) population dynamics are denso-dependent and autonomous from weather fluctuations. Annals of the Entomological Society of America 107, 136–142.

Imura, D., Toquenaga, Y. & Fujii, K. (2003) Genetic variation can promote system persistence in an experimental host-parasitoid system. Population Ecology 45, 205–212.

Jeger, M.J., Holt, J., Van Den Bosch, F. & Madden, L.V. (2004) Epidemiology of insect-transmitted plant viruses: modelling disease dynamics and control interventions. Physiological Entomology 29, 291–304.

Johnson, J. & Omland, K. (2004) Model selection in ecology and evolution. Trends in Ecology & Evolution 19, 101–108.

Johnson, L., Pecquerie, L. & Nisbet, R. (2013) Bayesain inference for bioenergetic models. Ecology 94, 882–889.

Johnson, L., Lafferty, K., Mcnally, E., Mordecai, E., Paaijmans, K., Pawar, S. & Ryan, S. (2014) Mapping the distribution of malaria: current approaches and future directions. Pages 191–211 in D. Chen, B. Moulin, and J. Wu, editors. In Analyzing and modeling spatial and temporal dynamics of infectious diseases. John Wiley & Sons, Hoboken, New Jersey, USA. pp. 191–211. John Wiley & Sons, Hoboken, USA.

Johnson, L.R., Ben-Horin, T., Lafferty, K.D., Mcnally, A., Mordecai, E., Paaijmans, K.P., Pawar, S. & Ryan, S.J. (2015) Understanding uncertainty in temperature effects on vector-borne disease: a Bayesian approach. Ecology 96, 203–213.

Johnson, L.R., Gramacy, R.B., Cohen, J., Mordecai, E., Murdock, C., Rohr, J., Ryan, S.J., Stewart-Ibarra, A.M. & Weikel, D. (2018) Phenomenological forecasting of disease incidence using heteroskedastic Gaussian processes: A dengue case study. The Annals of Applied Statistics 12, 27–66.

Keesing, F., Belden, L.K., Daszak, P., Dobson, A., Harvell, C.D., Holt, R.D., Hudson, P., Jolles, A., Jones, K.E. & Mitchell, C.E. (2010) Impacts of biodiversity on the emergence and transmission of infectious diseases. Nature 468, 647.

Kershner, I. & Lucas, G. (1980) The Empire strikes back. Twentieth Century-Fox Film Corporation.

Kersting, U., Satar, S. & Uygun, N. (1999) Effect of temperature on development rate and fecundity of apterous *Aphis gossypii* Glover (Hom., Aphididae) reared on *Gossypium hirsutum* L. Journal of Applied Entomology 123, 23–27.

Kindlmann, P. & Dixon, A.F.G. (2003) Insect predator–prey dynamics and the biological control of aphids by ladybirds. In First international symposium on biological control of arthropods. USDA Forest Service, USA pp. 118–124.

Kirk, D., Jones, N., Peacock, S., Phillips, J., Molnár, P.K., Krkošek, M. & Luijckx, P. (2018) Empirical evidence that metabolic theory describes the temperature dependency of within-host parasite dynamics. PLOS Biology 16, e2004608.

Kissling, W.D., Walls, R., Bowser, A., Jones, M.O., Kattge, J., Agosti, D., Amengual, J., Basset, A., Van Bodegom, P.M. & Cornelissen, J.H. (2018) Towards global data products of Essential Biodiversity Variables on species traits. Nature ecology & evolution 2, 1531–1540.

Koella, J.C. (2005) Malaria as a manipulator. Behavioural processes 68, 271–273.

Kramer, L.D., Hardy, J.L. & Presser, S.B. (1983) Effect of temperature of extrinsic incubation on the vector competence of *Culex tarsalis* for Western Equine Encephalomyelitis Virus. The American Journal of Tropical Medicine and Hygiene 32, 1130–1139.

Ladeau, S.L., Glass, G.E., Hobbs, N.T., Latimer, A. & Ostfeld, R.S. (2011) Data-model fusion to better understand emerging pathogens and improve infectious disease forecasting. Ecological Applications: A Publication of the Ecological Society of America 21, 1443–1460.

Laughton, A.M., Fan, M.H. & Gerardo, N.M. (2014) The combined effects of bacterial symbionts and aging on life history traits in the pea aphid, *Acyrthosiphon pisum*. Applied and environmental microbiology 80, 470–477.

Lloyd-Smith, J.O., Schreiber, S.J., Kopp, P.E. & Getz, W.M. (2005) Superspreading and the effect of individual variation on disease emergence. Nature 438, 355–359.

Lee, B.Y., Bacon, K.M., Bottazzi, M.E. & Hotez, P.J. (2013) Global economic burden of Chagas disease: a computational simulation model. The Lancet Infectious Diseases 13, 342–348.

Lefèvre, T. & Thomas, F. (2008) Behind the scene, something else is pulling the strings: emphasizing parasitic manipulation in vector-borne diseases. Infection, genetics and evolution 8, 504–519.

Logiudice, K., Ostfeld, R.S., Schmidt, K.A. & Keesing, F. (2003) The ecology of infectious disease: effects of host diversity and community composition on Lyme disease risk. Proceedings of the National Academy of Sciences 100, 567–571.

Lorenz, L. & Koella, J. (2011) Maternal environment shapes the life history and susceptibility to malaria of *Anopheles gambiae* mosquitoes. Malaria Journal 10, 1–8.

Macdonald, G. (1957) The epidemiology and control of malaria. Oxford University Press, London.

Martin, L.B., Addison, B., Bean, A.G., Buchanan, K.L., Crino, O.L., Eastwood, J.R., Flies, A.S., Hamede, R., Hill, G.E. & Klaassen, M. (2019) Extreme Competence: Keystone Hosts of Infections. Trends in Ecology & Evolution.

May, R.M. & Anderson, R.M. (1979) Population biology of infectious diseases: Part II. Nature 280, 455–461.

Mcelhany, P., Real, L.A. & Power, A.G. (1995) Vector preference and disease dynamics: a study of barley yellow dwarf virus. Ecology, 444–457.

Mcgill, B.J., Enquist, B.J., Weiher, E. & Westoby, M. (2006) Rebuilding community ecology from functional traits. Trends in Ecology & Evolution 21, 178–185.

Mead, P.S. (2015) Epidemiology of Lyme disease. Infectious Disease Clinics of North America 29, 187–210.

Metcalf, C.J.E., Graham, A.L., Martinez-Bakker, M. & Childs, D.Z. (2016) Opportunities and challenges of Integral Projection Models for modelling host–parasite dynamics. Journal of Animal Ecology 85, 343–355.

Molnár, P. (2013) Metabolic approaches to understanding climate change impacts on seasonal host-macroparasite dynamics. Ecology Letters 16, 9–21.

Molnár, P.K., Sckrabulis, J.P., Altman, K.A. & Raffel, T.R. (2017) Thermal performance curves and the metabolic theory of ecology—a practical guide to models and experiments for parasitologists. Journal of Parasitology 103, 423–439.

Mordecai, E.A., Cohen, J.M., Evans, M.V., Gudapati, P., Johnson, L.R., Lippi, C.A., Miazgowicz, K., Murdock, C.C., Rohr, J.R., Ryan, S.J., Savage, V., Shocket, M.S., Stewart Ibarra, A., Thomas, M.B. & Weikel, D.P. (2017) Detecting the impact of temperature on transmission of Zika, dengue, and chikungunya using mechanistic models. PLoS neglected tropical diseases 11, e0005568.

Mordecai, E.A., Paaijmans, K.P., Johnson, L.R., Balzer, C., Ben-Horin, T., De Moor, E., Mcnally, A., Pawar, S., Ryan, S.J., Smith, T.C. & Lafferty, K.D. (2013) Optimal temperature for malaria transmission is dramatically lower than previously predicted. Ecology Letters 16, 22–30.

Murdock, C.C., Luckhart, S. & Cator, L.J. (2017) Immunity, host physiology, and behaviour in infected vectors. Current Opinion in Insect Science 20, 28–33.

Murral, D.J., Nault, L.R., Hoy, C.W., Madden, L.V. & Miller, S.A. (1996) Effects of temperature and vector age on transmission of two Ohio strains of aster yellows phytoplasma by the aster leafhopper (Homoptera: Cicadellidae). Journal of Economic Entomology 89, 1223–1232.

Nakazawa, T., Yamanaka, T. & Urano, S. (2012) Model analysis for plant disease dynamics co-mediated by herbivory and herbivore-borne phytopathogens. Biology Letters 8, 685–688.

Nayar, J.K. & Sauerman, D.M. (1973) A comparative study of flight performance and fuel utilization as a function of age in females of Florida mosquitoes. Journal of insect physiology 19, 1977–1988.

Ng, K.C., Chaves, L.F., Tsai, K.H. & Chuang, T.W. (2018) Increased adult *Aedes aegypti* and *Culex quinquefasciatus* (Diptera: Culicidae) abundance in a dengue transmission hotspot, compared to a coldspot, within Kaohsiung City, Taiwan. Insects 9, 1–16.

Norberg, J., Swaney, D.P., Dushoff, J., Lin, J., Casagrandi, R. & Levin, S. (2001) Phenotyptic diversity and ecosystem functioning in chaning environments: a theoretical framework. PNAS 98, 11376–11381.

Novoseltsev, V.N., Michalski, A.I., Novoseltseva, J.A., Yashin, A.I., Carey, J.R. & Ellis, A.M. (2012) An age-structured extension to the vectorial capacity model. PLOS ONE 7, e39479.

Ohm, J.R., Baldini, F., Barreaux, P., Lefevre, T., Lynch, P.A., Suh, E., Whitehead, S.A. & Thomas, M.B. (2018) Rethinking the extrinsic incubation period of malaria parasites. Parasites & Vectors 11, 178.

Ohm, J.R., Teeple, J., Nelson, W.A., Thomas, M.B., Read, A.F. & Cator, L.J. (2016) Fitness consequences of altered feeding behaviour in immune-challenged mosquitoes. Parasites & Vectors 9, 113.

Parham, P.E. & Michael, E. (2010) Modeling the effects of weather and climate change on malaria transmission. Environmental Health Perspectives 118, 620–626.

Pawar, S., Dell, A.I. & Savage, V. (2015a) From Metabolic Constraints on Individuals to the Dynamics of Ecosystems. Aquatic Functional Biodiversity: An Ecological and Evolutionary Perspective, 3–36.

Pawar, S., Dell, A.I. & Savage, V.M. (2012) Dimensionality of consumer search space drives trophic interaction strengths. Nature 486, 485.

Pawar, S., Woodward, G. & Dell, A. (2015b) Trait-Based Ecology-from structure to function, 1st edition. Elsevier Academic Press, Oxford, UK.

Paweska, J.T., Venter, G.J. & Mellor, P.S. (2002) Vector competence of South African *Culicoides* species for bluetongue virus serotype 1 (BTV-1) with special reference to the effect of temperature on the rate of virus replication in *C. imicola* and *C. bolitinos*. Medical and Veterinary Entomology 16, 10–21.

Reiner, R.J., Perkins, T., Barker, C., Niu, T., Chaves, L., Ellis, A., George, D., Le Manach, A., Pulliam, J., Bisanzio, D., Buckee, C., Chiyaka, C., Cummings, D., Garcia, A., Gatton, M., ET AL. (2013) A systemic review of mathematical models of mosquito-bourne pathogen transmission: 1970:2010. J R Soc Interface 10, 20120921.

Renshaw, M., Service, M.W. & Birley, M.H. (1994) Size variation and reproductive success in the mosquito *Aedes cantans*. Medical and Veterinary Entomology 8, 179–186.

Rizzuto, M., Carbone, C. & Pawar, S. (2018) Foraging constraints reverse the scaling of activity time in carnivores. Nature ecology & evolution 2, 247.

Roux, O., Vantaux, A., Roche, B., Yameogo, K.B., Dabiré, K.R., Diabaté, A., Simard, F. & Lefèvre, T. (2015) Evidence for carry-over effects of predator exposure on pathogen transmission potential. Proceedings of the Royal Society B 282, 20152430.

San Martín, J.L., Brathwaite, O., Zambrano, B., Solórzano, J.O., Bouckenooghe, A., Dayan, G.H. & Guzmán, M.G. (2010) The epidemiology of dengue in the Americas over the last three decades: a worrisome reality. The American Journal of Tropical Medicine and Hygiene 82, 128–135.

Savage, V.M. (2004) Improved approximations to scaling relationships for species, populations, and ecosystems across latitudinal and elevational gradients. Journal of Theoretical Biology 227, 525–534.

Shapiro, L.L.M., Murdock, C.C., Jacobs, G.R., Thomas, R.J. & Thomas, M.B. (2016) Larval food quantity affects the capacity of adult mosquitoes to transmit human malaria. Proceedings of the Royal Society B 283, 20160298.

Smith, D.L., Battle, K.E., Hay, S.I., Barker, C.M., Scott, T.W. & Mckenzie, F.E. (2012) Ross, Macdonald, and a theory for the dynamics and control of mosquito-transmitted pathogens. PLoS Pathog 8, e1002588.

Smith, D.L., Perkins, T.A., Reiner, R.C., Barker, C.M., Niu, T., Chaves, L.F., Ellis, A.M., George, D.B., Menach, A.L., Pulliam, J.R.C., Bisanzio, D., Buckee, C., Chiyaka, C., Cummings, D.A.T., Garcia, A.J., ET AL. (2014) Recasting the theory of mosquito-borne pathogen transmission dynamics and control. Transactions of The Royal Society of Tropical Medicine and Hygiene 108, 185–197.

Soliman, B., Abo Ghalia, A., Shoukry, A. & Merdan, A. (1993) Mosquito age as a factor influencing the transmission of *Wuchereria bancrofti*. Journal of the Egyptian Society of Parasitology 23, 717–722.

Styer, L.M., Carey, J.R., Wang, J.-L. & Scott, T.W. (2007) Mosquitoes do senesce: departure from the paradigm of constant mortality. The American Journal of Tropical Medicine and Hygiene 76, 111–117.

Sun, L., Lee, C. & Hoeting, J.A. (2015) Parameter inference and model selection in deterministic and stochastic models via approximate Bayesian computation: modelling a wildlife epidemic. Environmentrics 26, 451–462.

Takken, W., Smallegange, R.C., Vigneau, A.J., Johnston, V., Brown, M., Mordue-Luntz, A. & Billingsley, P.F. (2013) Larval nutrition differentially affects adult fitness and Plasmodium development in the malaria vectors *Anopheles gambiae* and *Anopheles stephensi*. Parasites & Vectors 6, 345.

Taylor, R., Mordecai, E., Gilligan, C., Rohr, J. & Johnson, L. (2016) Mathematical models are a powerful method to understand and control the spread of Huanglongbing. PeerJ 4, e2642.

Vézilier, J., Nicot, A., Gandon, S. & Rivero, A. (2012) *Plasmodium* infection decreases fecundity and increases survival of mosquitoes. Proceedings of the Royal Society B 279, 4033–4041.

Webb, C.T., Hoeting, J.A., Ames, G.M., Pyne, M.I. & Leroy Poff, N. (2010) A structured and dynamic framework to advance traits-based theory and prediction in ecology. Ecology Letters 13, 267–283.

Wilson, A.J. & Mellor, P.S. (2009) Bluetongue in Europe: past, present and future. Philosophical Transactions of the Royal Society B: Biological Sciences 364, 2669–2681.

Wittmann, E.J., Mellor, P.S. & Baylis, M. (2002) Effect of temperature on the transmission of orbiviruses by the biting midge, *Culicoides sonorensis*. Medical and Veterinary Entomology 16, 147–156.

WORLD HEALTH ORGANIZATION (2017) Vector-borne diseases factsheet. www.who.int/mediacentre/factsheets/fs387/en/ [accessed 14 April 2017].

