## Supplemental Materials for "More than a flying syringe: Using functional traits in vector-borne disease research"

In [190]:

```
# Load some modules and settings
%matplotlib inline
import matplotlib.pyplot as plt
import matplotlib.ticker as ticker
from sympy import *
import scipy as sc
import numpy as np
init_printing()
from IPython.display import SVG, display
# print(plt.style.available)
```

In [191]:

```
plt.style.use('seaborn-darkgrid')
```

### *Supplementary Information for*:

### More than a flying syringe: using functional traits in Vector Borne Disease research

### Table of Contents


- 1 The trait-based framework for VBD research
  - 1.1 The meaning of "fitness effect"
  - 1.2 A trait-fitness parameter mapping example
    - 1.2.1 The mathematical model
- 2 Relationship of the trait-based framework to current approaches for incorporating trait variation into VBD dynamics
  - 2.1 Classical VBD models
    - 2.1.1 Derivation of $R_0$
  - 2.2 The classical compartment model with vector population dynamics
  - 2.3 The "plugging in" approach to incorporate trait variation into transmission models
- 3 Example of a trait-based model for VBD dynamics
  - 3.1 The vector's fitness
  - 3.2 From fitness to abundance
  - 3.3 Temperature-dependent trait variation
    - 3.3.1 Thermal performance curve parameterizations
  - 3.4 The effect of temperature-driven trait varation on vector abundance
    - 3.4.1 Temperature-driven abundance dynamics
  - 3.5 Temperature dependence of $R_0$
  - 3.6 Comparison with a phenomenological (non-trait-based) approach
- 4 Trait sensitivity analyses
  - 4.1 An example trait sensitivity analysis
    - 4.1.1 A numerical sensitivity analysis
- 5 References

### The trait-based framework for VBD research

Here we present details of the trait-based framework for studying Vector Borne Disease (VBD) systems. The general framework is illustration in the following Figure.

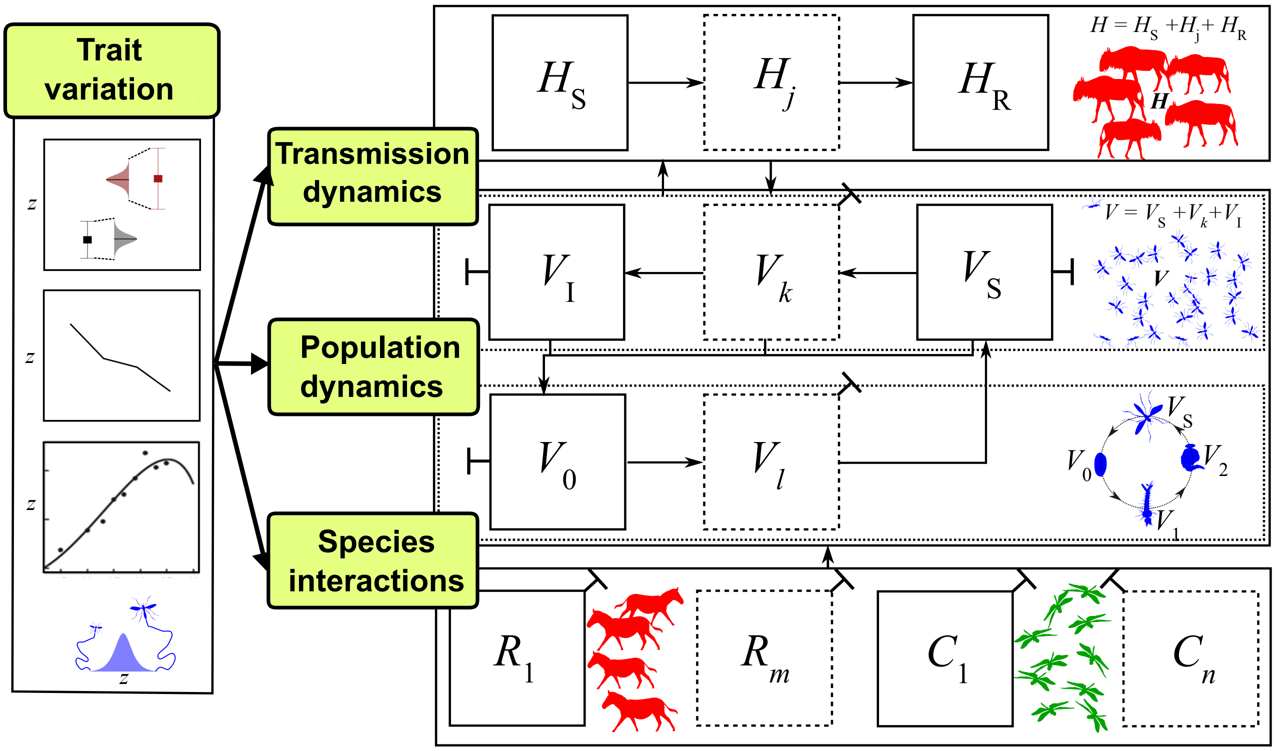


□→ Population or sub-population (compartment); ⬚→ Optional compartments; $H_S$, $V_I$, $V_0$, $R_1$ etc.: Abundance; T → Fitness (abundance) loss from compartment; ↑→ Fitness effect between compartments; $z$ → functional trait

Essentially, any specific trait-based VBD system's framework must be made up of:

- **Transmission compartments**:
  - Suceptible and Recovered host subpopulations (compartments) with $j\ (\geq 0)$ additional sub-populations to correctly model the transmission dynamics
  - Suceptible and Infected vector subpopulations with $k\ (\geq 0)$ additional sub-populations to correctly model the transmission dynamics
- **Vector life history compartments**: The vector's juvenile life stage subpopulations, starting with the stage at birth $(V_0)$ followed by $l\ (\geq 0)$ additional stages leading to the (susceptible) adult stage
- **Species interaction compartments**:
  - The Vector's $m\ (\geq 1)$ resource populations $(R_{1,2,\ldots m})$, which may be vector life stage-specific
  - The Vector's $n\ (\geq 1)$ consumer populations $(C_{1,2,\ldots m})$, which may also be vector life stage-specific
- **Trait Variation**
  - A suite of trait $(z)$ to parameter mappings that determine fitness effects (e.g., vector mortality, fecundity and biting rates)
  - A type of trait variation (one of $p(z)$, $dz/dt$, $dz/dE$, or a combination of these (main text Fig 3)) ))

Thus, the framework can be broken down into four (sequential) components or steps, each a mapping $(\rightarrow)$ that needs to be quantified through empirical studies coupled with mathematical modelling:

1. Trait → Fitness parameter
2. Trait-variation → Population Fitness
3. Population Fitness → Population Dynamics
4. Population Dynamics → Transmission Dynamics

For example, in the case of mosquito-borne arboviruses, the simplest possible specification would be:

- **The transmission compartments**: Number of Suceptible (S), Infected (I), and Recovered (R) hosts ($H_S$, $H_I$, $H_R$, respectively) and number of Suceptible, Infected, and Exposed (E) vectors ($V_S$, $V_I$ , $V_E$ respectively)
- **Vector life history compartments**: Number of vector individuals at egg, larval pupal and adult stages ($V_0$, $V_1$, $V_2$, and $V_S$ respectively)
- **Species interaction compartments**: Abundance of a single Resource species ($R_1$) that is the primary energy source of the vector population (which may actually be the host itself, so $R_1 = H_S$), and a single Consumer species ($C_1$) that is the primary source of mortality for the vector population (presumably only impacting certain vector life stages).
- **Trait Variation**: A suite of trait to parameter mappings that determine fitness effects (e.g., vector mortality, fecundity and biting rates), and a single type of trait variation, such as variation with an environmental factor ($dz/dE$; e.g., temperature)

For developing a mathematical model of such a system, the most common tool would be systems of ordinary differential equations (ODEs), as illustrated in the following sections.

#### The meaning of "fitness effect"

We define a fitness effect in a general way -- any parameter that changes the abundance value of any of the compartments in the trait-based VBD framework. So, for example, mortality rate (decreases abundance of any of the populations or sub-populations), fecundity (increases abundance), and biting rate (changes transmission rate, or the "fitness" of the disease transmission dynamic itself), all exert fitness effects. In the context of the transmission component, fitness essentially means that abundance of infected cases depends on the parasite's individual and population fitness (determined by the vector's traits, as this is a vector-focused framework).

### A trait-fitness parameter mapping example

Here we detail the trait to fitness mapping example summarized in Box 1 of the main text. The example uses recent advances in mechanistic, trait-based theory for species interactions [Pawar2012b, Dell2014a, Gilbert2014c, Pawar2015b, Rizzuto2017], using body size as a physical trait that is mapped through intermediate functional traits (metabolic rate, velocity, reaction distance) onto its biting rate, a functional trait and parameter that determines both vector abundance and transmission rate (two different types of fitness under our general definition of fitness). This approach has been empirically validated for consumer-resource interactions (especially, predation and herbivory) outside the context of VBDs, proving very effective in yielding feasible estimates of attack, biting, and consumption rates in vertebrates as well as invertebrates, including insects [Pawar2012b, Rizzuto2017].

The general schema of the trait-fitness parameter model is shown in the following figure.

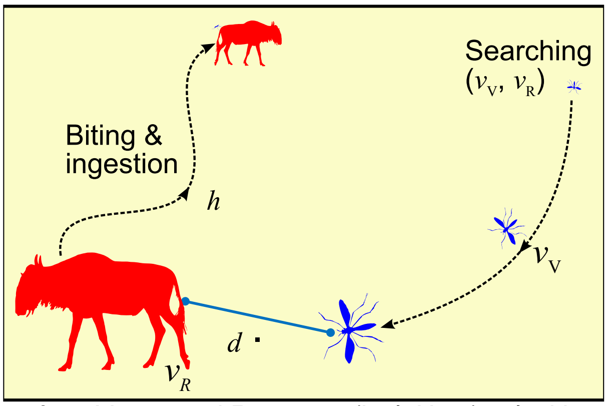


$v_V$ & $v_R$: Vector and Resource (incl. Host) velocities;
$h$ : Handling time; $d$ : Detection distance

#### The mathematical model

We consider a flying vector such as mosquitoes, flies or midges, seeking larger, warm-blooded hosts (a terrestrial mammal, such as a wildebeest (illustrated) or a human, which is also its energy resource). A general field encounter/consumption rate model for $2D$ (two-dimensional euclidean space) vector-host interactions is [Pawar2012b],

$$a = 2v_r d f(.) H \tag{1}$$

Here, $a$ is biting rate (bites $\cdot$ time$^{-1}$) for a given host density $H$ (individuals $\cdot$ area$^{-1}$), $v_r$ is relative velocity between the vector and host, $d$ is the reaction distance (minimum distance at which the vector detects and reacts to host), $f(.)$ is the host risk function that determines the (biting) functional response of the vector. This functional response encapsulates the constraints on the vector's handling time ($h$) (the post-encounter phase in the above figure), which would be determined by its ability to land on and complete the biting action successfully. We note that this interaction is $2D$ because even a flying vector effectively searches a two-dimensional plane (humans can't fly). The interaction between a walking or crawling vector such as tick and its mammalian host would also therefore be $2D$, but with a different velocity and reaction distance component (see below). Assuming each encounter between vector and host results in one bite, we now derive biting rate as a function of underlying traits.

Assuming random movement of vector and host individuals in the landscape, relative velocity $v_r$ is the square root of the sum of the average velocities of vector and the host individuals ($v_r = \sqrt{v_V^2 + v_H^2}$) [Pawar2012b, Dell2014a]. Assuming terrestrial vectors tend to move around widely in their search for hosts which are usually much larger than themselves and also restricted to patches (e.g., villages), the hosts can be considered relatively "sessile" with respect to their vector such that $v_V >> v_R$ in the search phase of the vector-host interaction. That is, $v_r \approx v_V$, effectively a "grazing" interaction [Pawar2012b, Pawar2015b, Dell2014a]). Thus, a critical component of biting rate hinges purely on the a vector functional trait, flight velocity, $v_V$. This in turn, depends on the vector's body size, because flight velocity scales with body mass due to biomechanical constraints:

$$v_r = v_V = v_0 m_V^{p_v}, \tag{2}$$

where $m_V$ is the vector's body mass, $v_0$ is a scaling exponent that depends partly on temperature and partly on flying mode (e.g., flying vs walking), and $p_v$ is a scaling exponent (usually between 0.1 and 0.3).

Furthermore, reaction distance $d$, is also expected to scale with body size because of the size-dependence of both, the vector's sensory organs and the host's ability to broadcast a signal (both in terms of volume and distance) [Pawar2012b]:

$$d = d_0 m_V^{2p_d} k^{p_d} \tag{3}$$

where $d_0$ is as in the case of velocity a scaling constant that may depend on temperature, $p_d$ a exponent respectively, and $k = \frac{m_H}{m_V}$, is the host-host body size-ratio.

Substituting Eqns.2, and 3 into1 and simplifying, we get

$$a = 2v_0 d_0 m_V^{p_v+2p_d} k^{p_d} f(.)H \tag{4}$$

In the case where hosts are relatively rare under field conditions, $f(.) = 1$, because handling time is not a dominant constraint (e.g., a Type I functional response is relevant). In cases where the hosts are relatively common (e.g., vectors live in the vicinity of the hosts, such is often the case in human villages), handling time would pay a bigger role, and a Type II functional response would come into play [Pawar2012b]. Previous work suggests that handling time $h$ itself also depends on size and temperature, and models can be built to make this relationship explicit and empirically testable [Vucic-Pestic2010b, Pawar2012b].

This trait to fitness parameter mapping can lead new insights and hypotheses, illustrating the value of a trait-based approach. In particular,

- It can be modified to accommodate vectors with different moving or sensory modes (e.g., ticks vs mosquitoes), and for the same host density (say, humans) predicts a lower maximal biting rate for smaller flying vectors relative to larger ones (e.g., midges vs. mosquitoes) and also lower biting rates for walking vectors compared to flying ones (e.g., ticks vs. mosquitoes).
- If temperature effects on vector velocity and detection were included (through the scaling constants $v_0$ and $d_0$), the effect of long or short term climatic or micro-climatic temperature changes can be mechanistically incorporated into biting rate, and thus, ultimately, vector abundance and disease transmission dynamics. Based on previous work, because metabolic rate and therefore search velocity increases approximately exponentially with temperature, this would predict a similar (approximately exponential) increase in biting rate with temperature [Dell2011c, Dell2014a], which has indeed been shown to be the case in mosquitoes [Mordecai2012, Mordecai2017]. Below we develop a trait-based model that includes temperature-driven biting rate for mosquitoes, which does not actually include a mechanistic trait-parameter mapping. The next step would be to include a mechanistic temperature-driven trait to biting rate model as we have done here.
- It provides a quantitative basis for understanding and predicting the effects of temperature-driven (developmental) body size decreases (the "size-temperature rule"; [Forster2012]) on biting rate. Based on eqn 4, temperature-driven decreases in body size are expected to decrease vector biting rate in a non-linear (allometric power law) way, but this may be counter-balanced by the increase in velocity with temperature - an exciting avenue for future work.

Such trait-fitness parameter mappings can derived for other key life-history and transmission parameters as well, such as vector fecundity and mortality using bio-mechanical, metabolic trait-based theory.

### Relationship of the trait-based framework to current approaches for incorporating trait variation into VBD dynamics

#### Classical VBD models

Omitting both trait variation and vector population (life history) dynamics from the general framework yields the classical compartment (SIR or SEIR) models for VBDs. As an illustration, we analyze such a model for malarial transmission by a mosquito vector (a Ross-Macdonald type model):

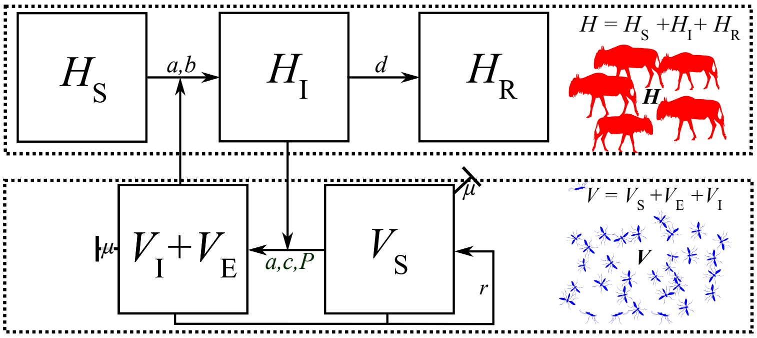


Similar models can and have been used for other vector borne diseases. The model can be specified mathematically by the following system of of coupled ODEs:

$$\begin{aligned}
\frac{dH_S}{dt} &= -\frac{ab}{H} V_I H_S \\
\frac{dH_I}{dt} &= \frac{ab}{H} V_I(t-\tau) H_S(t-\tau) - dH_I \\
\frac{dH_R}{dt} &= dH_I \\
\frac{dV_S}{dt} &= rV - \frac{ac}{H} H_I V_S - \mu V_S \\
\frac{dV_E}{dt} &= \frac{ac}{H} H_I V_S - \frac{ace^{-\mu P}}{H} H_I(t-P) V_S(t-P) - \mu V_E \\
\frac{dV_I}{dt} &= \frac{ace^{-\mu P}}{H} H_I(t-P) V_S(t-P) - \mu V_I,
\end{aligned} \tag{5}$$

where

$$\begin{aligned} H &= H_S + H_I + H_R \\ V &= V_S + V_E + V_I \end{aligned}$$

That is, the host population ($H$) is divided into susceptible ($H_S$), infected ($H_I$), and recovered (or immune) ($H_R$) classes, while the vector population ($V$) is divided into susceptible ($V_S$), Exposed ($V_E$), and infected ($V_I$) classes. Both host ($H$) and vector $V$ populations are assumed (and are by definition of the model) constant. The Model's parameters are:

| Parameter | Definition | Units |
|---|---|---|
| $d$ | Recovery rate of infected hosts | 1/time |
| $\tau$ | Exposure period in hosts | time |

| Parameter | Definition | Units |
| --- | --- | --- |
| $r$ | Vector population's reproduction or birth rate (may be maximal population growth rate, $r_m$) | 1/time |
| $a$ | Vector biting rate | bites / time |
| $b$ | Proportion of the bites by infective mosquitoes that produce infection in susceptible humans | dimensionless |
| $c$ | Proportion of bites by susceptible mosquitoes on infectious humans that infect mosquitoes (thus, $bc$ is vector competence) | dimensionless |
| $\mu$ | Mortality rate of adult vectors | 1/time |
| $P$ | Parasite's extrinsic incubation period | time |

Note that the parameters $\tau$, the time that a susceptible (S) takes to become infected after receiving a bite from infected vector ($V_I$), and $P$, the period that an exposed vector $V_E$ takes to become infectious (be able to infect a susceptible host (S)), introduce time-delays, making this a system of delay-differential equations.

#### Derivation of $R_0$

To calculate the basic reproductive number of the disease, we can use the next generation matrix method [Diekmann2010]. Given our model equations, we define the transmission matrix $\mathscr{T}$ and the transition matrix $\mathbf{\Sigma}$ as follows:

$$\mathscr{T} = \begin{bmatrix} 0 & 0 & \frac{abH_S}{H} \\ \frac{acV_S}{H} & 0 & 0 \\ 0 & 0 & 0 \end{bmatrix}, \mathbf{\Sigma} = \begin{bmatrix} -d & 0 & 0 \\ -\frac{acV_S}{H}e^{-\mu P} & -\mu & 0 \\ \frac{acV_S}{H}e^{-\mu P} & 0 & -\mu \end{bmatrix} \tag{6}$$

We consider a disease-free steady state, i.e., where both host and vector populations are entirely made up of susceptible individuals: $V_S = V$ and $H_S = H$. Then, $R_0$ is the square root of the largest eigenvalue of the product matrix $-\mathscr{T}\mathbf{\Sigma}^{-1}$:

$$-\mathscr{T}\mathbf{\Sigma}^{-1} = \begin{bmatrix} 0 & 0 & ab \\ \frac{acV}{H} & 0 & 0 \\ 0 & 0 & 0 \end{bmatrix} \times \begin{bmatrix} \frac{1}{d} & 0 & 0 \\ -\frac{Vac}{H\mu d}e^{-\mu P} & \frac{1}{\mu} & 0 \\ \frac{Vac}{H\mu d}e^{-\mu P} & 0 & \frac{1}{\mu} \end{bmatrix} = \begin{bmatrix} \frac{Va^2bc}{H\mu d}e^{-\mu P} & 0 & \frac{ab}{\mu} \\ \frac{Vac}{Hd} & 0 & 0 \\ 0 & 0 & 0 \end{bmatrix}$$

That is,

$$R_0 = \left( \frac{Va^2bce^{-\mu P}}{Hd\mu} \right)^{1/2} \tag{8}$$

This is equation 1 of the main text.

### The classical compartment model with vector population dynamics

The vector's population dynamics can be included in the the classical compartment VBD model [May1979a, Anderson1979]. This essentially entails making explicit the life history dynamics underlying the vector fitness parameter ($r$). For example, the above specification of the classical model for mosquito-borne arboviruses can be extended as follows:

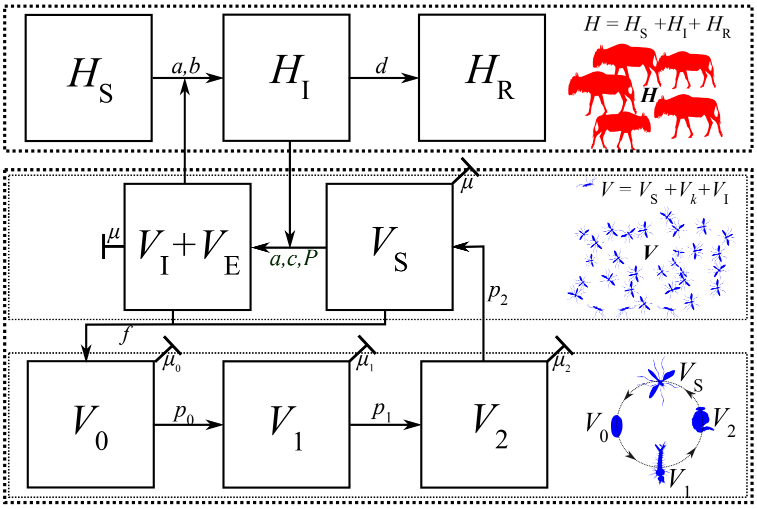


Mathematically, this can be achieved by defining an additional set of ODEs that determine the rate of production (fitness $r$; Eqn [5]) of new (susceptible) adult vectors ($V$).

$$\begin{aligned}\frac{dV}{dt} &= p_2V_2 - (f + \mu)V \\ \frac{dV_2}{dt} &= p_1V_1 - (p_2 + \mu_2)V_2 \\ \frac{dV_1}{dt} &= p_0V_0 - (p_1 + \mu_1)V_1 \\ \frac{dV_0}{dt} &= fV - (p_0 + \mu_0)V_0\end{aligned} \qquad (9)$$

where $p_i$ is the recruitment of each stage into the $i + 1^{\text{th}}$ stage, $f$ is adult fecundity, and $\mu_i$ is the mortality of the $i^{\text{th}}$ juvenile stage while $\mu$ is adult mortality. Each recruitment parameter may in fact be a function that depends on time delays (e.g., development time) [Beck-Johnson2013, Amarasekare2013]. Adding abundance dynamics to a classical compartment models is essentially a step towards the full trait-based framework.

### The "plugging in" approach to incorporate trait variation into transmission models

A few more recent studies have sought to incorporate trait variation directly into the classical VBD model by "plugging in" trait variation. This entails simply defining some of the parameters of the classical model as functions in terms of traits, as illustrated below.

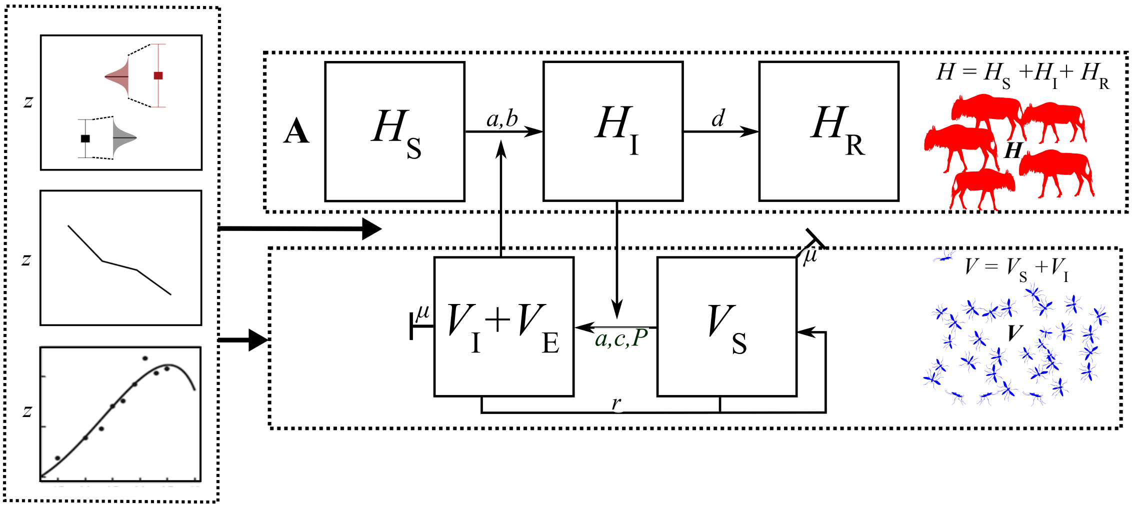


To illustrate this, consider the approach used by Mordecai et al [Mordecai2012, Johnson2015b, Mordecai2017]), who used the $R_0$ equation (Eqn. 8). The steps they took are:

1. Redefine $P$ as inverse of the parasite development rate (PDR), i.e., $P = \dfrac{1}{PDR}$
2. Define vector population size to be $V = \dfrac{\lambda}{\mu}$, where $\lambda$ is the *total* birth rate of adult mosquitoes in the *whole* population (adults/day), and $\mu$ is per-capita adult mortality rate (1/day). This is based on Parham & Michael [Parham2010], who derive the expression phenomenologically by treating $V$ as a random variable.Thus, $\lambda$ is equivalent to $rV$ in eqn 4 above.
3. Define the total vector population birth rate as $\lambda = \dfrac{EFD\ p_J\ MDR}{\mu}$ where $EFD$ is the number of eggs produced by all females in the population per day, $p_J$ is the probability that an egg will hatch and the larva will survive to adult, and $MDR$ is the average egg - to - adult development rate (inverse of development time). Then, the abundance of the vector becomes,

$$V = \frac{EFD\ p_J\ MDR}{\mu^2}$$

Substituting this into Eqn 8 yields a $R_0$ equation where traits underlying its parameters have been plugged in:

$$R_0 = \left( \frac{EFD\ p_J\ MDR\ a^2\ b\ c\ e^{-\mu/PDR}}{H\ d\ \mu^3} \right)^{1/2} \tag{10}$$

### Example of a trait-based model for VBD dynamics

We now develop a trait-based model for vector fitness ($r$) and the resulting transmission rate $R_0$ (leading to the results shown in Figure 3 of the main text). We compare this model's predictions with a more phenomenological approach. The model covers a subset of the full trait-based framework as illustrated below (the darker components shown).

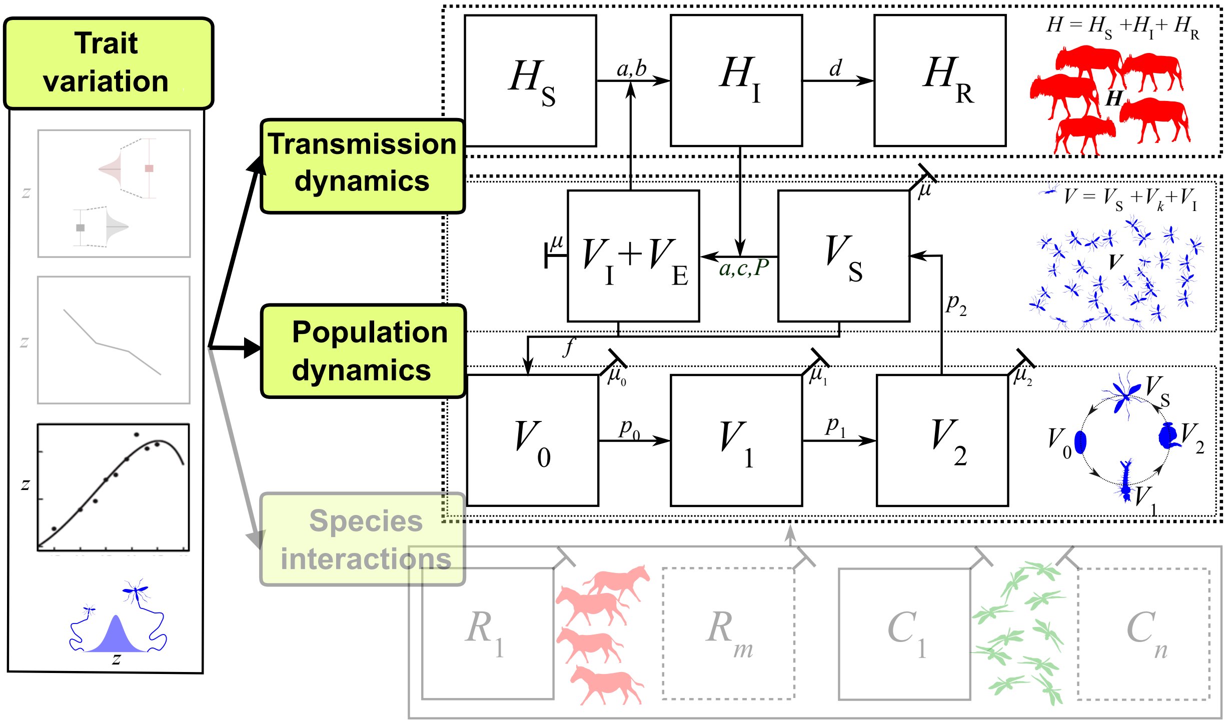


That is, we focus on one type of trait variation (environment-driven, with temperature as the variable) and ignore Consumer-Resource interactions (the bottom compartments of the full framework).

#### The vector's fitness

For simplicity, instead of directly simulating or analyzing the ODE system shown in Eqns [7], we focus on the asymptotic fitness from the discretized, matrix representation of the stage-structured life-history dynamics. In age/stage-structured populations, the expected lifetime reproductive success of a newborn individual in a population growing at the rate $r_m$ that has achieved a stable age distribution (SAD) can be described using the continuous form of the Euler-Lotka equation:

$$\int_{\alpha}^{\infty} \mathrm{e}^{-r_m x} l_x b_x \, \mathrm{d}x = 1 \qquad (11)$$

where $r_m$ is the (asymptotic) population growth rate, $a$ is the age of first reproduction, $l_x$ is the age-specific survivorship, and $b_x$ is the age-specific fecundity. Solving this equation gives the population growth rate, $r_m$ as a function of life-history parameters/traits. The Euler-Lotka equation is derived under the assumption of a SAD (proportions of the population in each age class constant over time), but if temperatures fluctuate sufficiently, the population may never reach a SAD ([Amarasekare2013]). Exploring this is beyond the scope of our model for illustrating a trait-based VBD approach, but would be an important area of future research.

In [192]:

```
x, l_a, b_pk, alpha, mu, mu_J, kappa, T, V_0, K, t = var('x l_a b_pk alpha mu mu
_J kappa T V_0 K t',real=True, positive = true) #assign symbolic variables

r_m  = var('r_m', real = True) # r_m can be negative
```

As arthropods and especially disease vectors are expected to have a Type III survivorship curve, given an instantaneous mortality rate, $z$, age-specific survivorship, $l_x$, declines exponentially with age and can be modeled as:

$$l_x = l_\alpha \mathrm{e}^{-\mu(x-\alpha)} \tag{12}$$

Here, $l_\alpha$ is the proportion of eggs surviving to adulthood (age $\alpha$), which, assuming a fixed instantaneous mortality rate across all juvenile stage, $\mu_J$, can be modelled as:

$$l_\alpha = \mathrm{e}^{-\int_0^\alpha \mu_J \, \mathrm{d}x} = \mathrm{e}^{-\mu_J \alpha} \tag{13}$$

In [193]:

```
l_alpha = exp(-integrate(mu_J,(x,0,alpha))); l_alpha
```

Out[193]:

$$e^{-\alpha\mu_J} \tag{14}$$

This, when substituted into the $l_x$ equation then gives

$$l_x = \mathrm{e}^{-(\mu_J\alpha + \mu(x-\alpha))} \tag{15}$$

The figures below show the shape of the $l_x$ model. This model can be easily adapted for a variable mortality rate, for example due to senescence, often modelled using a gompertz-type equation.

In [194]:

```
l_x = l_alpha * exp(-mu*(x - alpha)); simplify(l_x)
```

Out[194]:

$$e^{-\alpha\mu_J + \mu(\alpha - x)} \tag{16}$$

(where $x \geq \alpha$). Next, age-specific fecundity $b_x$ is expected to reach a peak, $b_{pk}$, shortly after maturation and then decline gradually with age. We model this using an exponential function

$$b_x = b_{pk} \mathrm{e}^{\kappa(\alpha - x)} \tag{17}$$

where we assume that fecundity peaks at the age of first reproduction, which occurs immediately at the time of adult emergence ($\alpha$), and $\kappa$ is a shape parameter (units of day$^{-1}$; can be interpreted as fecundity loss rate) that controls the spread of the fecundity schedule, that is, the rate of decline in fecundity after its peak. It is also possible that there is a systematic temperature-dependence of the delay between $\alpha$ and the age of peak fecundity. This can be modelled using a shifted exponential or gompertz equation, but is outside the scope of the current study. The figures below show the shape of the $b_x$ model.

In [195]:

```
b_x = b_pk * exp(-kappa*(x - alpha)); simplify(b_x)
```

Out[195]:

$$b_{pk} e^{\kappa(\alpha - x)} \tag{18}$$

In [196]:

```
#assign parameter values
V_0_par = 1 # Starting value for abundance - arbitrary
t_par = 30 # time for abundance calculation (approx value of development time at
 10degC)
K_par = 100 # carrying capacity for abundance - arbitrary

mu_J_par = .05 #1/day
mu_par = .03 #1/day
alp_par = 25. #days
b_pk_par = 10. #individuals/(individual * day)
kap_par = .1 #1/day
```

In [197]:

```
#Numerically evaluate
x_vec = np.arange(0, 60, 0.1) #vector of ages

# use lambidfy to speed up:
l_x_lam = lambdify((mu_J, mu, alpha,x), l_x, np)
l_x_vec = l_x_lam(mu_J_par, mu_par, alp_par, x_vec[x_vec>alp_par])

fig = plt.figure(); ax = fig.add_subplot(111)
ax.plot(x_vec[x_vec > alp_par], l_x_vec)
ax.set_xlabel('Age (days)', fontsize=14, fontweight = 'bold')
ax.set_ylabel('Survivorship ($l_x$)', fontsize=14, fontweight = 'bold')
ax.text(sc.mean(x_vec[x_vec>alp_par]), sc.amax(l_x_vec),
        '$\mu_J = ' + str(mu_J_par)+'$ \n' +
        '$\mu = ' + str(mu_par)+'$ \n' +
        r'$\alpha = ' + str(alp_par)+'$ \n',
        horizontalalignment='left', verticalalignment='top', fontsize=15)
plt.savefig('Results/lxModel.pdf')
```

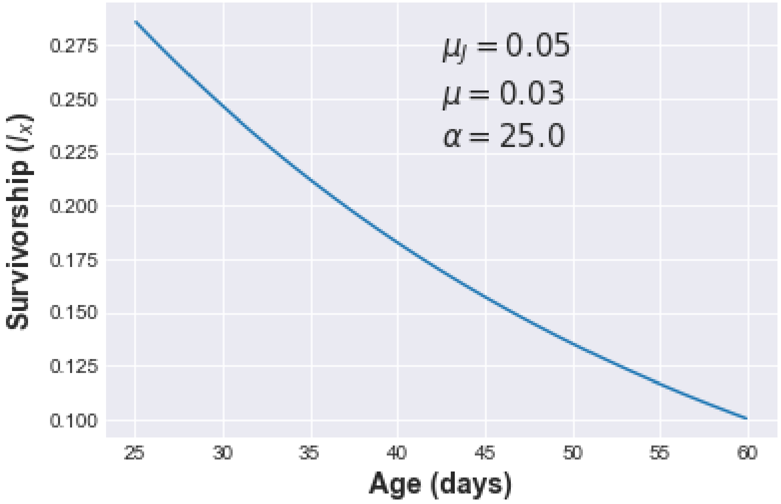

In [198]:

```
fig = plt.figure(); ax = fig.add_subplot(111)
#b_x_vec = np.array([b_x.evalf(subs = {b_pk:b_pk_par, x:age, alpha:alp_par, kapp
a:kap_par}) for age in x_vec])#use lambidy to speed up
b_x_lam = lambdify((b_pk, kappa, alpha, x), b_x, np)
b_x_vec = b_x_lam(b_pk_par, kap_par, alp_par, x_vec[x_vec>alp_par])

ax.plot(x_vec[x_vec>alp_par], b_x_vec);
ax.set_xlabel('Age (days)', fontsize=14, fontweight = 'bold');
ax.set_ylabel('Fecundity ($b_x$)', fontsize=14, fontweight = 'bold')
ax.text(sc.mean(x_vec[x_vec>alp_par]), sc.amax(b_x_vec)/2,
        r'$\kappa = ' + str(kap_par)+'$ \n'+
        r'$\alpha = ' + str(alp_par)+'$ \n' +
        r'$b_{pk} = ' + str(b_pk_par)+'$ \n',
        horizontalalignment='left', verticalalignment='top', fontsize=15)
plt.savefig('Results/bxModel.pdf')
```

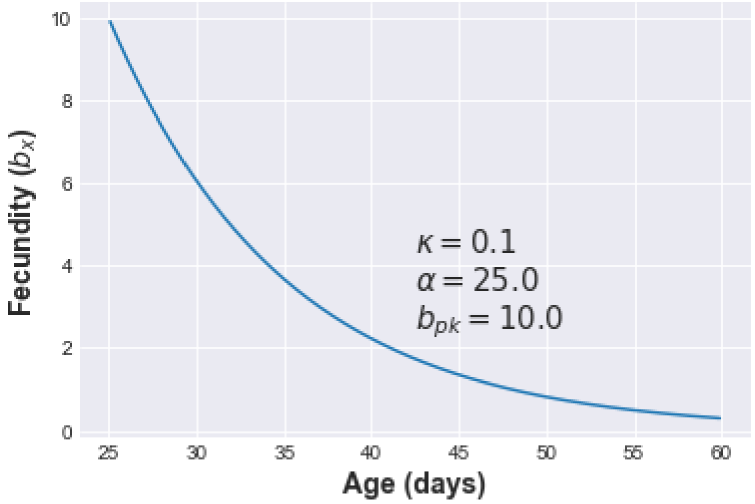


In [199]:

```
fig = plt.figure(); ax = fig.add_subplot(111) #now plot lxmx

ax.plot(x_vec[x_vec>alp_par], b_x_vec*l_x_vec);
ax.set_xlabel('Age (days)', fontsize=14, fontweight = 'bold');
ax.set_ylabel('$l_x b_x$', fontsize=14, fontweight = 'bold')
plt.savefig('Results/lxbxModel.pdf')
```

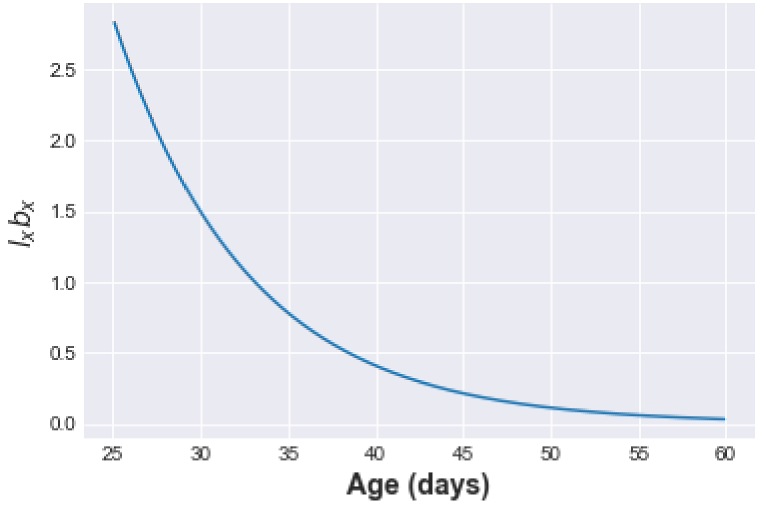

Substituting the $l_x$ and $b_x$ models into the Euler-Lotka Equation, we get

$$\int_{\alpha}^{\infty} b_{pk} \mathrm{e}^{-\alpha\mu_J+\kappa(\alpha-x)-rx+\mu(\alpha-x)}\,\mathrm{d}x = 1 \tag{19}$$

i.e.,

$$b_{pk}\mathrm{e}^{-\alpha\mu_J}\int_{\alpha}^{\infty} \mathrm{e}^{-rx+(\kappa+\mu)(\alpha-x)}\,\mathrm{d}x = 1 \tag{20}$$

In [200]:

```
EuLo = exp(-r_m * x) * l_x * b_x; simplify(EuLo)
```

Out[200]:

$$b_{pk}e^{-\alpha\mu_J+\kappa(\alpha-x)+\mu(\alpha-x)-r_m x} \tag{21}$$

In [201]:

```
# integrate(EuLo,(x,alpha,oo)) #doesn't work - parameters need to be constraine
d/bounded
# So integrate EuLo on sagemath using integrate(EuLo, x, alpha, infinity), with
 positivity constraints on all parameters, which gives:
EuLo_int = b_pk*exp(-alpha*mu_J)/((kappa + r_m)*exp(alpha*r_m) + mu*exp(alpha*r_
m)); simplify(EuLo_int)
```

Out[201]:

$$\frac{b_{pk}e^{-\alpha\left(\mu_J+r_m\right)}}{\kappa+\mu+r_m} \tag{22}$$

And evaluating the integral produces after simplification:

$$\frac{b_{pk}e^{-\alpha\left(r+\mu_J\right)}}{\kappa+r+\mu} = 1 \tag{23}$$

Solving this for $r_m$ gives:

In [202]:

```
r_SP = solve(EuLo_int-1,r_m); r_SP = simplify(r_SP[0]);r_SP
```

Out[202]:

$$\frac{1}{\alpha}\left(-\alpha\left(\kappa+\mu\right)+\operatorname{LambertW}\left(\alpha b_{pk}e^{\alpha\left(\kappa+\mu-\mu_J\right)}\right)\right) \tag{24}$$

The LambertW function part is difficult to use for sensitivity analyses (and also hard to interpret), so instead, we approximate this by taking log of the solution to the integral above and perform a power series expansion around $r_m = 0$ of just the term $\ln(\kappa + r + z)$:

In [203]:

```
series(ln(r_m + kappa + mu),r_m, 0,3) #
# That is, exp(log(kappa + z) + r/(kappa + z))
```

Out[203]:

$$\log(\kappa+\mu)+\frac{r_m}{\kappa+\mu}-\frac{r_m^2}{2\kappa^2+4\kappa\mu+2\mu^2}+\mathcal{O}\left(r_m^3\right) \tag{25}$$

That is,

$$\frac{b_{pk}e^{-\alpha\mu_J}}{\mu e^{\alpha r_m}+(\kappa+r_m)\,e^{\alpha r_m}}\approx\log(\kappa+\mu)+\frac{r_m}{\kappa+\mu}-\frac{r_m^2}{2\kappa^2+4\kappa\mu+2\mu^2}+\mathcal{O}\left(r_m^3\right) \tag{26}$$

Then, substituting the first two terms back into the integral solution and solving for $r_m$ gives:

In [204]:

```
#substitute # exp(log(kappa + z) + r/(kappa + z))
EuLo_int_app = b_pk*exp(-alpha*(r_m + mu_J))/ exp(log(kappa + mu) + r_m/(kappa +
 mu))

r_SP_app = solve(EuLo_int_app-1, r_m); r_SP_app = simplify(r_SP_app[0]); r_SP_ap
p
```

Out[204]:

$$-\frac{1}{\alpha\kappa+\alpha\mu+1}\left(\alpha\kappa\mu_J+\alpha\mu\mu_J+\log\left(\frac{(\kappa+\mu)^{\kappa+\mu}}{b_{pk}^{\kappa+\mu}}\right)\right) \tag{27}$$

Thus the approximation is:

$$r_m\approx\frac{-\alpha\kappa\mu_J-\alpha\mu\mu_J+\kappa\log\left(\frac{b_{pk}}{\kappa+\mu}\right)+\mu\log\left(\frac{b_{pk}}{\kappa+\mu}\right)}{\alpha(\kappa+\mu)+1} \tag{28}$$

that is,

$$r_m\approx\frac{(\kappa+\mu)\left(\log\left(\frac{b_{pk}}{\kappa+\mu}\right)-\alpha\mu_J\right)}{\alpha(\kappa+\mu)+1} \tag{29}$$

**Accuracy of the $r_m$ approximation**

We now show that this approximation is very good as long as $r_m$ is relatively small ($<1$). This range of $r_m$ values is typically where most insect growth rates lie (https://www.journals.uchicago.edu/doi/full/10.1086/506977) [Frazier2006].

In [205]:

```
from scipy.special import lambertw # need this for the lambertw

bpk_vec = np.arange(.1, 1000, 0.1).astype(float) #vector of b_pk's

#r_SP_app_vec = np.array([r_SP_app.evalf(subs = {b_pk:b_pkVal,z_J:z_J_par, alph
a:alp_par, z:z_par, kappa:kap_par}) for b_pkVal in bpk_vec])
#use lambidy to speed up:
r_SP_app_lam = lambdify((b_pk, mu_J, mu, kappa, alpha), r_SP_app, np)
r_SP_app_vec = r_SP_app_lam(bpk_vec, mu_J_par, mu_par, kap_par, alp_par)

# have to specify r_SP with lambertw:
# r_SP_vec = (-a*(kappa + z) + lambertw((a_par*bpk_vec*exp(a_par*(kap_par + z_pa
r - z_J_par))).astype('float')).real)/a_par
tmp = lambertw((alp_par*bpk_vec*exp(alp_par*(kap_par + mu_par - mu_J_par))).asty
pe('float')).real
r_SP_vec = -(kap_par + mu_par) + tmp/alp_par

fig = plt.figure(); ax = fig.add_subplot(111)
ax.plot(r_SP_vec, r_SP_app_vec)
ax.plot(np.arange(-1,5,0.1), np.arange(-1,5,0.1), "k--")
ax.set_xlim([min(r_SP_vec),max(r_SP_vec)])
ax.set_ylim([min(r_SP_app_vec),max(r_SP_app_vec)])
ax.set_xlabel('$r$ (exact)', fontsize=14, fontweight = 'bold');
ax.set_ylabel('$r$ (approximation)', fontsize=14, fontweight = 'bold')
plt.savefig('Results/rapprox0.pdf')
```

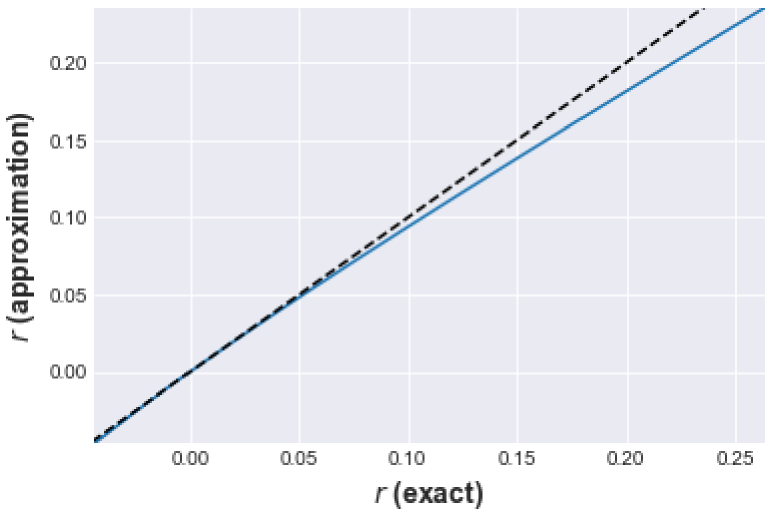

In [206]:

```
fig = plt.figure(); ax = fig.add_subplot(111)
ax.plot(r_SP_vec, r_SP_vec/r_SP_app_vec)
ax.set_xlabel('$r$ (exact)', fontsize=14, fontweight = 'bold');
ax.set_ylabel('Approximation as $\%$ of exact value', fontsize=14, fontweight =
'bold')
plt.savefig('Results/rapprox1.pdf')
```

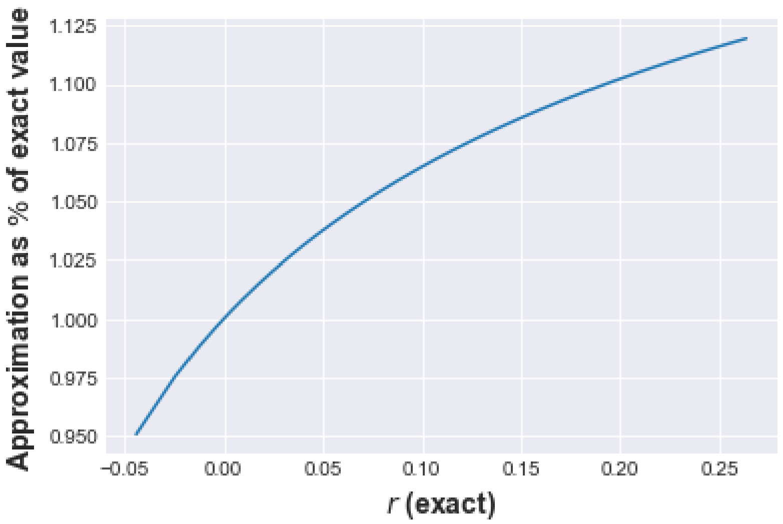


The parameters underlying fitness and their units are summarized in the following table:

| Parameter | Units | Description |
|---|---|---|
| $a$ | day | Age of maturation (Juvenile to adult development time) |
| $b_{pk}$ | individuals (eggs) × individual (female) × day $^{-1}$ , i.e., day$^{-1}$ | Peak reproductive rate |
| $\mu$ | day$^{-1}$ | Adult mortality rate |
| $\mu_J$ | day$^{-1}$ | Mortality rate across all juvenile stages |
| $\kappa$ | day$^{-1}$ | Fecundity loss rate (inverse of spread of the fecundity schedule) |

### From fitness to abundance

Finally, $r_m$ needs to be converted into vector population abundance ($V$). For this, it is reasonable to assume that the population grows logistically:

$$\frac{\mathrm{d}V}{\mathrm{d}t} = r_m V\left(1 - \frac{V}{K}\right) \tag{30}$$

where $K$ is the carrying capacity of the environment (maximum possible number of adult female mosquitoes). This has the time-dependent solution:

$$V_t = \frac{N_0 K \mathrm{e}^{r_m t}}{K + V_0(\mathrm{e}^{r_m t} - 1)} \tag{31}$$

Note that the effect of precipitation and temperature on resource (e.g., breeding site availability in mosquitoes) can be incorporated into $K$. However, for simplicity, and because the focus of our study is to determine the sensitivity of $V$ (and not it's absolute value) to life history traits we assume that $K$ is both precipitation and temperature-independent.

In [207]:

```
V = V_0*K*exp(r_m * t)/(K + V_0*(exp(r_m * t) -1))
```

### Temperature-dependent trait variation

We describe the thermal performance curve for each life history parameter/trait using the Sharpe-Schoolfield model, or its inverse:

In [208]:

```
# Assign functions
B_0, E, E_D, T_pk, k, T, T_ref = var('B_0 E E_D T_pk k T T_ref')
```

In [209]:

```
B = B_0 * exp(-E * ((1/(k*(T))) - (1/(k*T_ref)))) /(1 + (E / (E_D - E)) * exp((E
_D / k)* ((1/T_pk) - (1/T)))); simplify(B)
```

Out[209]:

$$-\frac{B_0\,(E - E_D)\,e^{\frac{E\left(T - T_{ref}\right)}{T T_{ref} k}}}{E e^{\frac{E_D\left(T - T_{pk}\right)}{T T_{pk} k}} - E + E_D} \tag{32}$$

In [210]:

```
B_inv = simplify(1/B); B_inv
```

Out[210]:

$$\frac{e^{-\frac{E\left(T-T_{ref}\right)}{TT_{ref}k}}}{B_0\left(E-E_D\right)}\left(-Ee^{\frac{E_D\left(T-T_{pk}\right)}{TT_{pk}k}}+E-E_D\right) \tag{33}$$

The Sharpe-Schoolfield model provides a mechanistic interpretation of the thermal performance curves of biological traits but could be replaced with alternative unimodal models, including statistical model such as Briere-type or polynomial models, without having a qualitative impact on the results.

In [211]:

```
# Assign generic parameter values
k_par = 8.617 * 10**-5
E_par = .55
E_D_par = 4
T_pk_par = 25.3
T_ref_par = 10+273.15
B_0_par = 0.5
```

In [212]:

```
T_vec = 273.15+np.arange(0, 40, 0.05) #Vector of temperatures

B_lam = lambdify((B_0, E, T_pk, T_ref, E_D,k,T), B, np)
B_vec = B_lam(B_0_par,E_par,T_pk_par+273.15,T_ref_par,E_D_par,k_par,T_vec)

B_inv_lam = lambdify((B_0, E, T_pk, T_ref, E_D,k,T), B_inv, np)
B_inv_vec = B_inv_lam(B_0_par,E_par,T_pk_par+273.15,T_ref_par,E_D_par,k_par,T_ve
c)

fig = plt.figure(figsize=(10, 4)); ax = fig.add_subplot(121)

ax.plot(T_vec-273.15, B_vec);

ax.annotate(s='', xy=(T_vec[250]-273.15,B_vec[250]*1.05), xytext=(T_vec[350]-27
3.15,B_vec[350]*1.05), arrowprops=dict(arrowstyle='<-'))
ax.text(T_vec[300]-273.15, B_vec[300]*1.1, ' $E_A$',  horizontalalignment='righ
t', verticalalignment='bottom', fontsize=13)

ax.annotate(s='', xy=(T_vec[610]-273.15,B_vec[610]*1.1), xytext=(T_vec[660]-273.
15,B_vec[660]*1.1), arrowprops=dict(arrowstyle='<-'))
ax.text(T_vec[635]-273.15, B_vec[635]*1.15, ' $E_D$',  horizontalalignment='lef
t', verticalalignment='bottom', fontsize=13)

ax.plot([T_pk_par, T_pk_par], [0,ax.get_ylim()[1]], color='k', linestyle='--', l
inewidth=1)
ax.text(T_pk_par, B_0_par/2, ' $T_{pk}$',  horizontalalignment='left', verticala
lignment='center', fontsize=13)

ax.plot([0,10], [B_0_par, B_0_par], color='k', linestyle='--', linewidth=1)
ax.plot([10, 10], [0,B_0_par], color='k', linestyle='--', linewidth=1)
ax.text(5, B_0_par, '$B_0$',  horizontalalignment='center', verticalalignment='b
ottom', fontsize=13)
ax.text(10, B_0_par/2, ' $T_{ref}$',  horizontalalignment='left', verticalalignm
ent='center', fontsize=13)

ax.set_title('A: Sharpe-Schoolfield model', fontsize=13, fontweight = 'bold');
ax.set_xlabel('Temperature $(^\circ C)$', fontsize=14, fontweight = 'bold');
ax.set_ylabel('Trait Value ($B$)', fontsize=14, fontweight = 'bold')

ax = fig.add_subplot(122)
ax.plot(T_vec-273.15, B_inv_vec);
ax.plot([T_pk_par, T_pk_par], [0,ax.get_ylim()[1]], color='k', linestyle='--', l
inewidth=1)
ax.set_title('B: Inverse Sharpe-Schoolfield model', fontsize=13, fontweight = 'b
old');
ax.set_xlabel('Temperature $(^\circ C)$', fontsize=14, fontweight = 'bold');
fig.tight_layout()

plt.savefig('Results/TPCModel.pdf')
```

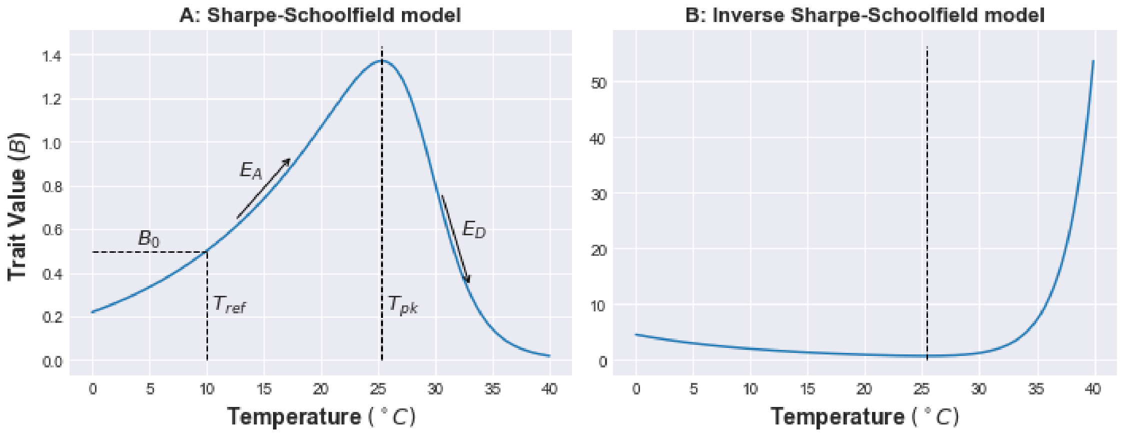


#### Thermal performance curve parameterizations

We incorporate trait variation into fitness by including thermal performance curves (TPCs) of each trait. These TPCs were parametrised to have equal sensitivity to temperature by assigning all traits the same values for thermal optimum $(T_{pk})$, activation energy $(E)$ and deactivation energy $(E_D)$.

The thermal optimum and activation energy were parameterized using the median values for the data from Mordecai et al [Mordecai2012, Johnson2015b]. These are also consistent with the median values found by Dell et al. [Dell2011c] in an meta-analysis of thermal performance curves across a range of taxa for these categories of traits.

The normalisation constant $(B_0)$ was parametrised for each trait in order that the trait value at the thermal optimum is of a scale consistent with empirical observations of these traits in disease vectors. For this too, we used the thermal perfomance curve parameters obtained by Mordecai et al. [Mordecai2012, Mordecai2017] for mosquitoes.

In addition to the traits underlying vector fitness and abundance, calculation of $R_0$ requires three more temperature-dependent parameters (eqn 8): vector biting rate $a$, vector competence $bc$, and the parasite's extrinsic incupbation period (which we will assume is the inverse of parasite development rate (PDR), following Mordecai et al [Mordecai2012, Johnson2015b]. Also, following Mordecai et al, we assume these are all unimodal functions of temperature, also modeled using the Sharpe-Schoolfield equation.

All these parameterizations are summarized in the following table.

| Model parameter | Normalization constant ($B_0$) | Thermal optimum ($T_{pk}$) | Thermal Sensitivity ($E$) | $E_D$ |
|---|---|---|---|---|
| $\alpha$ | 25 | 22.5 | 0.65 | 3 |
| $b_{pk}$ | 10 | 22.5 | 0.65 | 3 |
| $\mu$ | 0.03 | 22.5 | 0.65 | 3 |
| $\mu_J$ | 0.05 | 22.5 | 0.65 | 3 |
| $\kappa$ | 0.1 | 22.5 | 0.65 | 3 |
| $a$ | .15 | 22.5 | 0.65 | 3 |
| $bc$ | 24 | 22.5 | 0.65 | 3 |
| PDR | .05 | 22.5 | 0.65 | 3 |

**Parameter values for the thermal performance curves of traits underlying vector abundance and $R_0$.** We use the approximate mean of the thermal optima across traits reported in Mordecai et al [Mordecai2012, Johnson2015b]. For simplicity we assume a common thermal optimum as well as activation and deactivation energy across all traits based on the median values from Mordecai et al's data. This serves the purpose of illustrating the trait-based VBD framework, but exploring the effects of variation in these TPC parameters across traits and species is a potentially rich area for future work.

The resulting TPCs are plotted below.

In [213]:

```
#Assign trait-specific TPC values:

B_0_alp =  1/alp_par#days; inverse because we are using inverse of the Sharpe-Sc
hoolfield model
T_pk_alp = 22.5
E_alp = .55
E_D_alp = 3

B_0_bpk = b_pk_par #individuals/(individual * day)
T_pk_bpk = 22.5
E_bpk = .55
E_D_bpk = 3

B_0_mu = 1/mu_par  #1/day; inverse because we are using inverse of the S-S TPC m
odel
T_pk_mu = 22.5
E_mu = .55
E_D_mu = 3

B_0_muJ = 1/mu_J_par  #1/day; inverse because we are using inverse of the S-S TP
C model
T_pk_muJ = 22.5
E_muJ = .55
E_D_muJ = 3

B_0_kap = kap_par #1/day
T_pk_kap = 22.5
E_kap = .55
E_D_kap = 3

# Additional R_0 parameters:
a_par = .2
bc_par = 35
PDR_par = .05

B_0_a = a_par #1/day
T_pk_a = 22.5
E_a = .55
E_D_a = 3

B_0_bc = bc_par
T_pk_bc = 22.5
E_bc = .55
E_D_bc = 3

B_0_PDR = PDR_par
T_pk_PDR = 22.5
E_PDR = .55
E_D_PDR = 3
```

In [214]:

```
#Calculate the TPCs

alp_vec = B_inv_lam(B_0_alp,E_alp,T_pk_alp+273.15,T_ref_par,E_D_alp,k_par,T_vec)
bpk_vec = B_lam(B_0_bpk,E_bpk,T_pk_bpk+273.15,T_ref_par,E_D_bpk,k_par,T_vec)
mu_vec = B_inv_lam(B_0_mu,E_mu,T_pk_mu+273.15,T_ref_par,E_D_mu,k_par,T_vec)
muJ_vec = B_inv_lam(B_0_muJ,E_muJ,T_pk_muJ+273.15,T_ref_par,E_D_muJ,k_par,T_vec)
kap_vec = B_lam(B_0_kap,E_kap,T_pk_kap+273.15,T_ref_par,E_D_kap,k_par,T_vec)

a_vec = B_lam(B_0_a,E_a,T_pk_a+273.15,T_ref_par,E_D_a,k_par,T_vec)

bc_vec = B_lam(B_0_bc,E_bc,T_pk_bc+273.15,T_ref_par,E_D_bc,k_par,T_vec)
PDR_vec = B_lam(B_0_PDR,E_PDR,T_pk_PDR+273.15,T_ref_par,E_D_PDR,k_par,T_vec)

fig = plt.figure(figsize=(12, 5))

ax = fig.add_subplot(241)
ax.plot(T_vec-273.15, alp_vec);
ax.plot([T_pk_alp, T_pk_alp], [0,ax.get_ylim()[1]], color='k', linestyle='--', l
inewidth=1)
ax.set_xlabel('Temperature $(^\circ C)$', fontsize=14, fontweight = 'bold');
ax.set_ylabel(r'$\alpha$', fontsize=14, fontweight = 'bold')

ax = fig.add_subplot(242)
ax.plot(T_vec-273.15, bpk_vec);
ax.plot([T_pk_bpk, T_pk_bpk], [0,ax.get_ylim()[1]], color='k', linestyle='--', l
inewidth=1)
ax.set_xlabel('Temperature $(^\circ C)$', fontsize=14, fontweight = 'bold');
ax.set_ylabel('$b_{pk}$', fontsize=14, fontweight = 'bold')

ax = fig.add_subplot(243)
ax.plot(T_vec-273.15, mu_vec);
ax.plot([T_pk_mu, T_pk_mu], [0,ax.get_ylim()[1]], color='k', linestyle='--', lin
ewidth=1)
ax.set_xlabel('Temperature $(^\circ C)$', fontsize=14, fontweight = 'bold');
ax.set_ylabel('$\mu$', fontsize=14, fontweight = 'bold')

ax = fig.add_subplot(244)
ax.plot(T_vec-273.15, muJ_vec);
ax.plot([T_pk_muJ, T_pk_muJ], [0,ax.get_ylim()[1]], color='k', linestyle='--', l
inewidth=1)
ax.set_xlabel('Temperature $(^\circ C)$', fontsize=14, fontweight = 'bold');
ax.set_ylabel('$\mu_J$', fontsize=14, fontweight = 'bold')

ax = fig.add_subplot(245)
ax.plot(T_vec-273.15, kap_vec);
ax.plot([T_pk_kap, T_pk_kap], [0,ax.get_ylim()[1]], color='k', linestyle='--', l
inewidth=1)
ax.set_xlabel('Temperature $(^\circ C)$', fontsize=14, fontweight = 'bold');
ax.set_ylabel(r'$\kappa$', fontsize=14, fontweight = 'bold')

ax = fig.add_subplot(246)
ax.plot(T_vec-273.15, a_vec);
ax.plot([T_pk_a, T_pk_a], [0,ax.get_ylim()[1]], color='k', linestyle='--', linew
idth=1)
ax.set_xlabel('Temperature $(^\circ C)$', fontsize=14, fontweight = 'bold');
ax.set_ylabel(r'$a$', fontsize=14, fontweight = 'bold')

ax = fig.add_subplot(247)
ax.plot(T_vec-273.15, bc_vec);
```

```
ax.plot([T_pk_bc, T_pk_bc], [0,ax.get_ylim()[1]], color='k', linestyle='--', linewidth=1)
ax.set_xlabel('Temperature $(^\circ C)$', fontsize=14, fontweight = 'bold');
ax.set_ylabel(r'$bc$', fontsize=14, fontweight = 'bold')

ax = fig.add_subplot(248)
ax.plot(T_vec-273.15, PDR_vec);
ax.plot([T_pk_PDR, T_pk_PDR], [0,ax.get_ylim()[1]], color='k', linestyle='--', linewidth=1)
ax.set_xlabel('Temperature $(^\circ C)$', fontsize=14, fontweight = 'bold');
ax.set_ylabel(r'PDR', fontsize=14, fontweight = 'bold')


fig.tight_layout()

plt.savefig('Results/TraitTPCs.pdf')
```

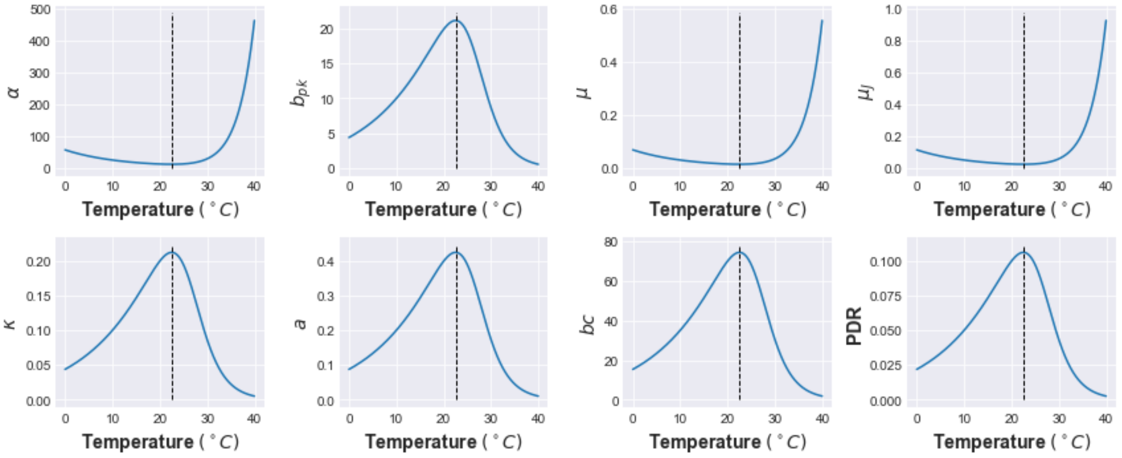


### The effect of temperature-driven trait varation on vector abundance

To determine the temperature dependence of the vector population growth rate, we substitute the thermal performance curves of the life history traits into the approximation of $r_m$ derived above:

In [215]:

```
r_m_vec = r_SP_app_lam(bpk_vec, muJ_vec, mu_vec, kap_vec, alp_vec)

fig = plt.figure(); ax = fig.add_subplot(111)
ax.plot(T_vec-273.15, r_m_vec);
ax.plot(T_vec-273.15, [0]*len(T_vec), color='k', linestyle='--', linewidth=1)
ax.set_xlabel('Temperature $(^\circ C)$', fontsize=14, fontweight = 'bold');
ax.set_ylabel('$r_m$', fontsize=14, fontweight = 'bold')
plt.ylim((-.1,r_m_vec.max()+.01))

plt.savefig('Results/r_TPC.pdf')
```

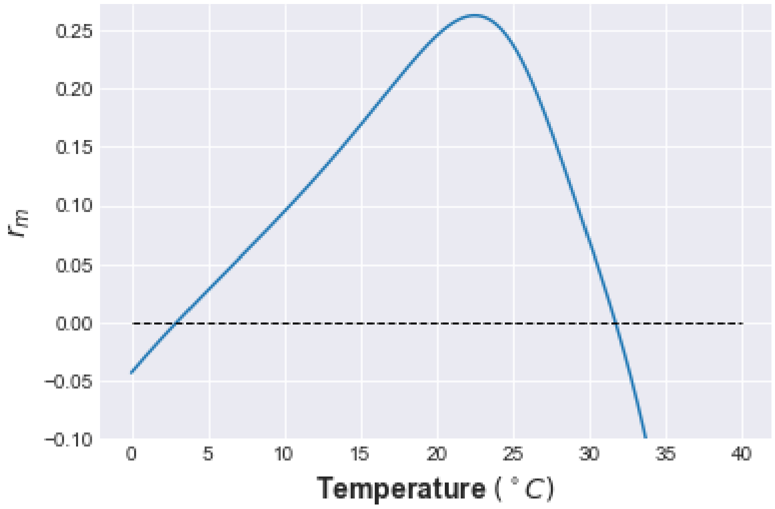


The effect of temperature-driven trait variation on population growth rate, $r_m$. The horizontal line delineates the the range of temperatures for with population growth rate is greater than 0.

Finally, we can substitute the temperature-dependent $r_m$ into the population growth model (the logistic equation) to determine the temperature dependence of vector population density:

In [216]:

```
V_lam = lambdify((r_m, K, V_0, t), V, np)

V_vec = V_lam(r_m_vec, K_par, V_0_par, t_par)

fig = plt.figure(); ax1 = fig.add_subplot(111)
ax1.plot(T_vec-273.15, V_vec);
ax1.plot([T_vec[V_vec==V_vec.max()]-273.15, T_vec[V_vec==V_vec.max()]-273.15], [
ax1.get_ylim()[0],ax1.get_ylim()[1]], color='k', linestyle='--', linewidth=1)
ax1.set_xlabel('Temperature $(^\circ C)$', fontsize=14, fontweight = 'bold');
ax1.set_ylabel('$V$', fontsize=14, fontweight = 'bold')
ax1.xaxis.set_major_locator(ticker.MultipleLocator(10))
ax1.yaxis.set_major_locator(ticker.MultipleLocator(20))

left, bottom, width, height = [0.18, 0.67, 0.2, 0.2]
ax2 = fig.add_axes([left, bottom, width, height])
ax2.plot(T_vec-273.15, r_m_vec, color='red')
ax2.plot(T_vec-273.15, [0]*len(T_vec), color='k', linestyle='--', linewidth=1)
ax2.set_ylabel('$r_m$', fontweight = 'bold',fontsize=14)
ax2.set_xlabel('Temperature $(^\circ C)$', fontweight = 'bold',fontsize=10)
ax2.tick_params(axis='both', which='both', bottom='off', top='off', labelbottom=
'off', right='off', left='off', labelleft='off')
plt.ylim((-.1,r_m_vec.max()+.01))

plt.savefig('Results/V_TPC.pdf')
```

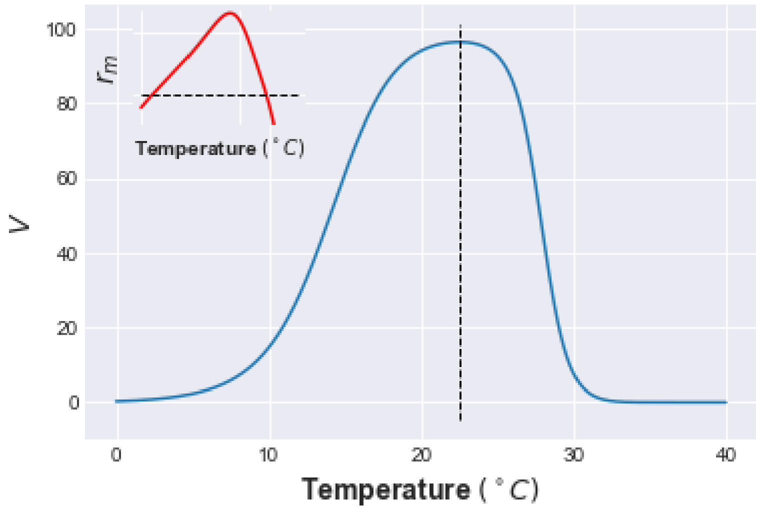

#### Temperature-driven abundance dynamics

To model environmental fluctuations that have dominant periodic or cyclical behavior (such as seasonal fluctuations), the following model can be used to generate a stationary time series' with the desired properties:

$$T_t = T_0 + A \sin \frac{2\pi t}{L} \tag{34}$$

where $T$ is temperature at time $t$ (so $T_t$ is temperature at time-point $t$), while A and L are the amplitude and length of the temperature cycles, respectively. If $T_t$ is further defined to be an autocorrelated or uncorrelated Gaussian random variable with a finite mean and variance, smaller scale fluctuations can also be simulated. Thus this this simple model can generate a wide range of time series' with fluctuations ranging from predominantly cyclical, to serially uncorrelated or correlated random noise.

Here is an example times series:

In [217]:

```
tlen = 365 # months
T_0 = 20
A = 5
L = 365
sigma = .1
t_vec = sc.arange(1.,tlen)

np.random.seed(0)

T_t = (T_0 + A * sc.sin(2 * sc.pi * t_vec / L)) + (sc.randn(len(t_vec))*sigma)

fig = plt.figure(figsize=(4, 3)); ax = fig.add_subplot(111)
ax.plot(t_vec, T_t)
ax.set_xlabel('Time (days)', fontsize=12, fontweight = 'bold');
ax.set_ylabel('Temperature, $T (^\circ$C)', fontsize=12, fontweight = 'bold')
ax.tick_params(axis='both', which='major', labelsize=11)
ax.xaxis.set_major_locator(ticker.MultipleLocator(tlen/5))
ax.yaxis.set_major_locator(ticker.MultipleLocator(2))

plt.savefig('Results/T_fluct.pdf')
```

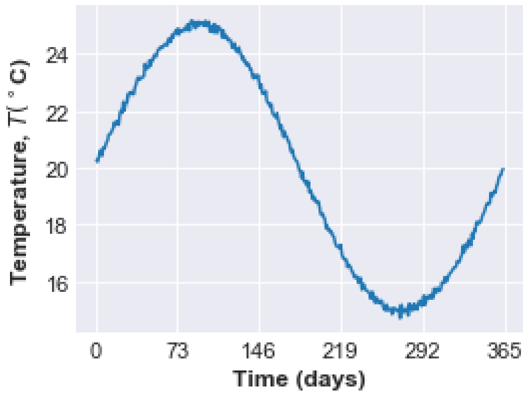


Figure xx: Time series over 5 years generated by eqn. 34, for monthly variation in temperature ($L$ = 12 months, $t$ = 120 months). Here $T_t$ is modeled as an uncorrelated Gaussian random variable. The other parameter values are $T_0 = 10^\circ$C, $A = 1.5^\circ$C, $\sigma = 1^\circ$C.

Now we can calculate the effect of a seasonally varying temperature on trait fitness:

In [218]:

```
alp_vec = B_inv_lam(B_0_alp,E_alp,T_pk_alp+273.15,T_ref_par,E_D_alp,k_par,T_t+ 273.15)
bpk_vec = B_lam(B_0_bpk,E_bpk,T_pk_bpk+273.15,T_ref_par,E_D_bpk,k_par,T_t+ 273.15)
mu_vec = B_inv_lam(B_0_mu,E_mu,T_pk_mu+273.15,T_ref_par,E_D_mu,k_par,T_t+ 273.15)
muJ_vec = B_inv_lam(B_0_muJ,E_muJ,T_pk_muJ+273.15,T_ref_par,E_D_muJ,k_par,T_t+ 273.15)
kap_vec = B_lam(B_0_kap,E_kap,T_pk_kap+273.15,T_ref_par,E_D_kap,k_par,T_t+ 273.15)

a_vec = B_lam(B_0_a,E_a,T_pk_a+273.15,T_ref_par,E_D_a,k_par,T_t+ 273.15)
bc_vec = B_lam(B_0_bc,E_bc,T_pk_bc+273.15,T_ref_par,E_D_bc,k_par,T_t+ 273.15)
PDR_vec = B_lam(B_0_PDR,E_PDR,T_pk_PDR+273.15,T_ref_par,E_D_PDR,k_par,T_t+ 273.15)

r_m_vec = r_SP_app_lam(bpk_vec, muJ_vec, mu_vec, kap_vec, alp_vec)

V_vec = V_lam(r_m_vec, K_par, V_0_par, t_par)
```

And plot the result:

In [219]:

```
fig, ax1 = plt.subplots(figsize=(4, 3))
color = 'blue'
ax1.set_xlabel('Time (days)', fontsize=16, fontweight = 'bold')
ax1.set_ylabel('Temperature, $T (^\circ$C)', fontsize=16, fontweight = 'bold',co
lor=color)
ax1.plot(t_vec, T_t, color=color, linestyle='-',linewidth=1)
ax1.tick_params(axis='y', labelcolor=color)
ax1.tick_params(axis='both', which='major', labelsize=14, direction = 'in')
ax1.xaxis.set_major_locator(ticker.MultipleLocator(75))
ax1.yaxis.set_major_locator(ticker.MultipleLocator(2))

ax2 = ax1.twinx()  # create a second axes that shares the same x-axis
color = 'red'
ax2.set_ylabel('Abundance, $V$', fontsize=16, fontweight = 'bold',color = color)
ax2.plot(t_vec, V_vec, color=color,linewidth=2)
ax2.tick_params(axis='y', labelcolor=color)
ax2.tick_params(axis='both', which='major', labelsize=14, direction = 'in')
ax2.yaxis.set_major_locator(ticker.MultipleLocator(20))

plt.savefig('Results/V_fluct.pdf')
```

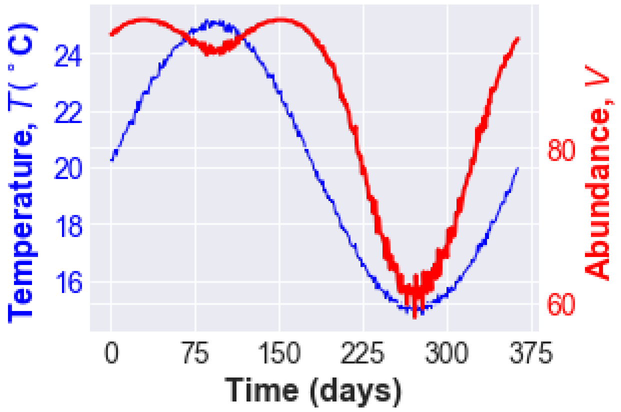


Thus the trait-based abundance model predicts a widening of the temporal niche of high vector fitness (and therefore abundance), with an early-year emergence as well as vector persistence into the cooler late summer season, with a dip in the warmest period of the summer season. This is partly driven by the lower thermal optima of underlying traits (22.5$^{\circ}$C) relative to the thermal maximum for this particular environment (25$^{\circ}$C).

#### Temperature dependence of $R_0$

We now calculate $R_0$ assuming temperature-driven traits underlying its parameters ()

In [220]:

```
H = 1
d = 1
R0_vec = (V_vec * a_vec**2 * bc_vec * sc.exp(-mu_vec*(1/PDR_vec)))/(H * d * mu_v
ec)
R0_vec = R0_vec/np.max(R0_vec)

fig, ax1 = plt.subplots(figsize=(4, 3))
color = 'blue'
ax1.set_xlabel('Time (days)', fontsize=16, fontweight = 'bold')
ax1.set_ylabel('Temperature, $T (^\circ$C)', fontsize=16, fontweight = 'bold',co
lor=color)
ax1.plot(t_vec, T_t, color=color, linestyle='-',linewidth=1)
ax1.tick_params(axis='y', labelcolor=color)
ax1.tick_params(axis='both', which='major', labelsize=14, direction = 'in')
ax1.xaxis.set_major_locator(ticker.MultipleLocator(75))
ax1.yaxis.set_major_locator(ticker.MultipleLocator(2))

ax2 = ax1.twinx()  # instantiate a second axes that shares the same x-axis
color = 'red'
ax2.set_ylabel('$R_0$', fontsize=16, fontweight = 'bold',color = color)
ax2.plot(t_vec, R0_vec, color=color,linewidth=2)
ax2.tick_params(axis='y', labelcolor=color)
ax2.tick_params(axis='both', which='major', labelsize=14, direction = 'in')
ax2.yaxis.set_major_locator(ticker.MultipleLocator(0.2))
plt.ylim((ax2.get_ylim()[0],R0_vec.max()+.01))

plt.savefig('Results/R0_fluct.pdf')
```

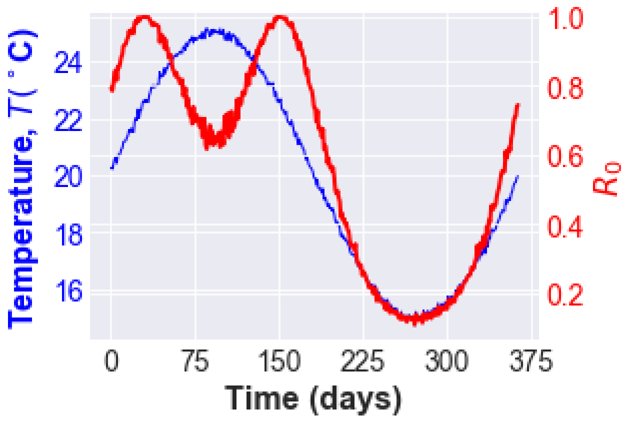


Thus, the trait-based VBD model predicts a widening of the temporal niche of disease transmission, with an early- as well as late-summer peak, with a dip in the warmest period of the summer season. The dip is more pronounced than abundance because of the additional environment-driven variation in transmission traits (biting rate, vectorial capacity and parasite development rate).

### Comparison with a phenomenological (non-trait-based) approach

Here we contrast the result of our trait-based abundance-$R_0$ model with a more phenomenological approach. In this approach (e.g., [Ng2018, Hoshi2014, Johnson2018]), Vector abundance $V$ is statistically associated with one or more environmental variables such as temperature, fitted using linear or non-linear lag (distributed or non-distributed) models (e.g., ARMAX; [Brockwell1991]). This results in a phenomenological, statistical temperature-abundance model. For example, the model may be a simple linear mapping of the temperature time series to abundance with time-lags:

$$V_t = \beta_0 + \beta_1 T_t + \beta_2 T_{t-1} + \ldots + \beta_n T_{t-n} \tag{35}$$

Where $T_t$ the temperature time series (generator), given by eqn 34, and the coefficients $\beta_0$ and $\beta_1$ and $\beta_2$ capture the numerical transformation of the temperature time series to the abundance time series as well as the lagged and unlagged effects of temperature on abundance.

In the case where the dominant time lag is the $j^{\text{th}}$ one, we can model the effect by manipulating the phase shift term in eqn. 34.

As an illustration, below we re-calculate $R_0$ by assuming a time-lag of two weeks between peak abundance and optimal temperature (approximately the combined effect of fecundity and development time delays in mosquitoes across 10-25˚C).

In [221]:

```
beta_0 = 5
beta_1 = 5
t_lag = -14 # lag, in days
np.random.seed(0)
T_t1 = (T_0 + A * sc.sin(2 * sc.pi * (t_vec+t_lag) / L)) + (sc.randn(len(t_vec))
*sigma)

#calculate lagged response

V_TS_vec = beta_0 + beta_1 * T_t1

fig, ax1 = plt.subplots(figsize=(4, 3))
color = 'blue'
ax1.set_xlabel('Time (days)', fontsize=16, fontweight = 'bold')
ax1.set_ylabel('Temperature, $T (^\circ$C)', fontsize=16, fontweight = 'bold',co
lor=color)
ax1.plot(t_vec, T_t, color=color, linestyle='-',linewidth=1)
ax1.tick_params(axis='y', labelcolor=color)
ax1.tick_params(axis='both', which='major', labelsize=14, direction = 'in')
ax1.xaxis.set_major_locator(ticker.MultipleLocator(75))
ax1.yaxis.set_major_locator(ticker.MultipleLocator(2))

ax2 = ax1.twinx()  # create a second fig that shares the same x-axis
color = 'red'
ax2.set_ylabel('Abundance, $V$', fontsize=16, fontweight = 'bold',color = color)
ax2.plot(t_vec, V_TS_vec, color=color,linewidth=2)
ax2.tick_params(axis='y', labelcolor=color)
ax2.tick_params(axis='both', which='major', labelsize=14, direction = 'in')
ax2.yaxis.set_major_locator(ticker.MultipleLocator(10))

plt.savefig('Results/V_TS_fluct.pdf')
```

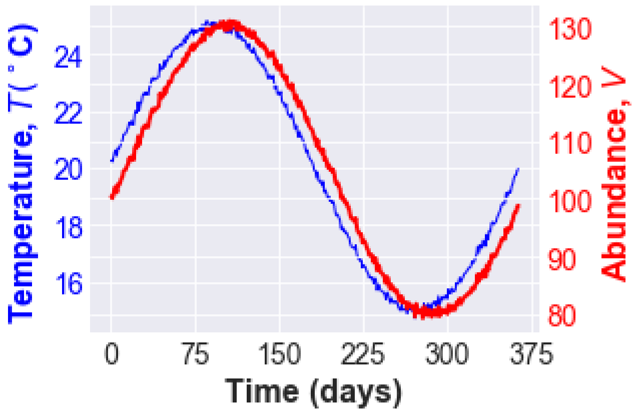


Now recalculate $R_0$, and rescale it to a max of 1 as in the case of th trait-based model above:

In [222]:

```
R0_vec = (V_TS_vec * a_par**2 * bc_par * sc.exp(-mu_par*(1/PDR_par)))/(H * d * m
u_par)
R0_vec = R0_vec/np.max(R0_vec)

fig, ax1 = plt.subplots(figsize=(4, 3))
color = 'blue'
ax1.set_xlabel('Time (days)', fontsize=16, fontweight = 'bold')
ax1.set_ylabel('Temperature, $T (^\circ$C)', fontsize=16, fontweight = 'bold',co
lor=color)
ax1.plot(t_vec, T_t, color=color, linestyle='-',linewidth=1)
ax1.tick_params(axis='y', labelcolor=color)
ax1.tick_params(axis='both', which='major', labelsize=14, direction = 'in')
ax1.xaxis.set_major_locator(ticker.MultipleLocator(75))
ax1.yaxis.set_major_locator(ticker.MultipleLocator(2))

ax2 = ax1.twinx()  # instantiate a second axes that shares the same x-axis
color = 'red'
ax2.set_ylabel('$R_0$', fontsize=16, fontweight = 'bold',color = color)
ax2.plot(t_vec, R0_vec, color=color,linewidth=2)
ax2.tick_params(axis='y', labelcolor=color)
ax2.tick_params(axis='both', which='major', labelsize=14, direction = 'in')
ax2.yaxis.set_major_locator(ticker.MultipleLocator(.1))
plt.ylim((ax2.get_ylim()[0],R0_vec.max()+.01))

plt.savefig('Results/R0_TS_fluct.pdf')
```

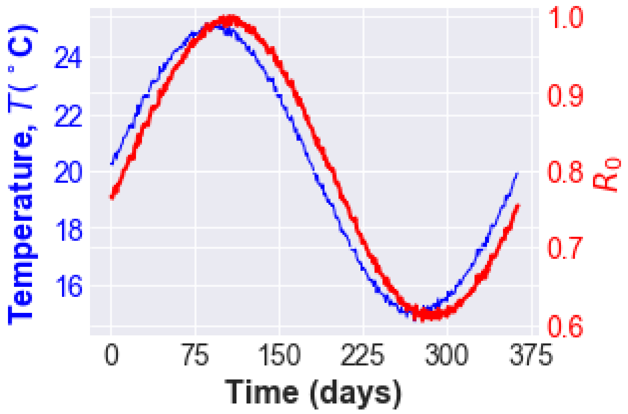


Thus a trait based approach provides new mechanistic insights and hypotheses compared to the phenomenological approach. In particular the trait-based model,

1. predicts a wider time period of high vector abundance and therefore transmission risk, while the phenomenological model predicts only a temporal shift in the risk due to inherent life history delays.
2. predicts a counter-intuitive decrease in transmission risk in the warmest period (but consistent with the results of Mordecai et al (2012) who used a plugging-in approach) to incorporate trait variation.
3. shows that environment-driven variation in underlying traits relative to the temperature regime matters, and why it matters.
4. reveals which traits matter the most (next section)

Ultimately, for more accurate predictions, the time delays included in the phenomenological model would also need to be incorporated into the trait-based model we have developed, which can be done by introducing time-delay functions into the trait-fitness parameter mappings.

### Trait sensitivity analyses

Here we show how a trait-based approach can also yield insights into the sensitivity of an emergent phenomenon such as vector abundance or transmission rate due to variation in underlying traits.

The general objective for such a trait sensitivity analysis is to mathematically or computationally (numerically) decompose the emergent phenomenon (e.g., transmission fitness $R_0$, abundance $V$, or population fitness $r_m$) into the contributions of underlying traits, through the trait-fitness mappings (trait $\rightarrow$ fitness parameter (including transmission parameters), trait variation $\rightarrow$ population fitness, fitness $\rightarrow$ abundance, and abundance $\rightarrow$ transmission).

An analytical way to tackle this problem, which is possible if the underlying mappings are mathematically differentiable (typically the case), is to decompose the the emergent variable using the chain rule of calculus. For example , we can use the chain rule to determine the relative contribution of the temperature dependence of each parameter to the temperature dependence of vector population abundance $V$ derived above as follows.

The relative contribution of a parameter can be expressed as the product of the partial derivative of population density ($V$) with respect to a parameter and the derivative of that parameter with respect to the environment driver (temperature in this case):

$$\frac{\mathrm{d}V}{\mathrm{d}T} = \frac{\partial V}{\partial b_{pk}}\frac{\mathrm{d}b_{pk}}{\mathrm{d}T} + \frac{\partial V}{\partial \alpha}\frac{\mathrm{d}\alpha}{\mathrm{d}T} + \frac{\partial V}{\partial \mu}\frac{\mathrm{d}\mu}{\mathrm{d}T} + \frac{\partial V}{\partial \mu_J}\frac{\mathrm{d}\mu_J}{\mathrm{d}T} + \frac{\partial V}{\partial \kappa}\frac{\mathrm{d}\kappa}{\mathrm{d}T} \tag{36}$$

Furthermore, as the partial derivative of $V$ with respect to each parameter can be expressed as:

$$\frac{\partial V}{\partial b_{pk}} = \frac{\partial V}{\partial r_m}\frac{\partial r_m}{\partial b_{pk}}, \; \frac{\partial V}{\partial \alpha} = \frac{\partial V}{\partial r_m}\frac{\partial r_m}{\partial \alpha}, \text{ etc.} \tag{37}$$

The relative contribution of each temperature driven variation in each trait to population density can be therefore be fully expressed as:

$$\frac{\mathrm{d}V}{\mathrm{d}T} = \frac{\partial V}{\partial r_m}\frac{\partial r_m}{\partial b_{pk}}\frac{\mathrm{d}b_{pk}}{\mathrm{d}T} + \frac{\partial V}{\partial r_m}\frac{\partial r_m}{\partial \alpha}\frac{\mathrm{d}\alpha}{\mathrm{d}T} + \frac{\partial V}{\partial r_m}\frac{\partial r_m}{\partial \mu}\frac{\mathrm{d}\mu}{\mathrm{d}T} + \frac{\partial V}{\partial r_m}\frac{\partial r_m}{\partial \mu_J}\frac{\mathrm{d}\mu_J}{\mathrm{d}T} + \frac{\partial V}{\partial r_m}\frac{\partial r_m}{\partial \kappa}\frac{\mathrm{d}\kappa}{\mathrm{d}T}$$

The above analytical approach can easily be extend to perform a trait sensitivity analysis on the transmission fitness ($R_0$) itself (e.g., [Mordecai2012, Johnson2015b]).

Depending on the type of trait variation considered ($p(z)$, $dz/dt$, or $dz/dE$) and the complexity of the trait-fitness mappings, numerical perturbation methods may need to be used instead of the above analytical method. below we illustrate such a numerical approach as well.

#### An example trait sensitivity analysis

To illustrate this approach, we assess the sensitivity of vector maximal growth rate (population fitness, $r_m$) through temperature dependent variation of the component traits. For this, we need to calculate each of the partial derivatives (the $\frac{\partial r_m}{\partial z}$'s):

In [223]:

```
dr_dbpk = diff(r_SP_app, b_pk); simplify(dr_dbpk)
```

Out[223]:

$$\frac{\kappa + \mu}{b_{pk} \left(\alpha\kappa + \alpha\mu + 1\right)} \tag{39}$$

In [224]:

```
dr_dalp = diff(r_SP_app, alpha); simplify(dr_dalp)
```

Out[224]:

$$\frac{1}{\left(\alpha\kappa + \alpha\mu + 1\right)^{2}} (\kappa + \mu) \left( \alpha\kappa\mu_J + \alpha\mu\mu_J - \mu_J \left(\alpha\kappa + \alpha\mu + 1\right) + \log \left( \right. \right.$$

In [225]:

```
dr_dmu = diff(r_SP_app, mu); simplify(dr_dmu)
```

Out[225]:

$$\frac{1}{\left(\alpha\kappa + \alpha\mu + 1\right)^{2}} \left( \alpha \left( \alpha\kappa\mu_J + \alpha\mu\mu_J + \log \left( \frac{(\kappa + \mu)^{\kappa+\mu}}{b_{pk}^{\kappa+\mu}} \right) \right) + (\alpha\kappa + \right.$$

In [226]:

```
dr_dmuJ = diff(r_SP_app, mu_J); simplify(dr_dmuJ)
```

Out[226]:

$$-\frac{\alpha (\kappa + \mu)}{\alpha\kappa + \alpha\mu + 1} \tag{42}$$

In [227]:

```
dr_dkap = diff(r_SP_app, kappa); simplify(dr_dkap)
```

Out[227]:

$$\frac{1}{\left(\alpha\kappa + \alpha\mu + 1\right)^{2}} \left( \alpha \left( \alpha\kappa\mu_J + \alpha\mu\mu_J + \log \left( \frac{(\kappa + \mu)^{\kappa+\mu}}{b_{pk}^{\kappa+\mu}} \right) \right) + (\alpha\kappa + \right.$$

For the TPC model, we ony need to calculate the derivative wrt $T$ for either the function $B$ (for parameters that are rates) or the 1/$B$ (for parameters that are times):

In [228]:

```
dBdT =  diff(B, T); simplify(dBdT)
```

Out[228]:

$$\frac{B_0 E}{T^2 k \left(E e^{\frac{E_D\left(T-T_{pk}\right)}{T T_{pk} k}} - E + E_D\right)^2}(E - E_D)\left(E_D e^{\frac{1}{T T_{pk} T_{ref} k}\left(E T_{pk}\left(T-T_{ref}\right)+E_D T_{ref}\left(T-\right.\right.}\right.$$

In [229]:

```
dB_invdT =  diff(B_inv, T); simplify(dB_invdT)
```

Out[229]:

$$\frac{E}{B_0 T^2 k}\left(e^{\frac{E_D}{T_{pk} k}-\frac{E_D}{T k}} - 1\right) e^{-\frac{E}{T_{ref} k}+\frac{E}{T k}} \quad (45)$$

Next we can evaluate (visualize the individual contributions/sensitivities):

In [230]:

```
alp_vec = B_inv_lam(B_0_alp,E_alp,T_pk_alp+273.15,T_ref_par,E_D_alp,k_par,T_vec)
bpk_vec = B_lam(B_0_bpk,E_bpk,T_pk_bpk+273.15,T_ref_par,E_D_bpk,k_par,T_vec)
mu_vec = B_inv_lam(B_0_mu,E_mu,T_pk_mu+273.15,T_ref_par,E_D_mu,k_par,T_vec)
muJ_vec = B_inv_lam(B_0_muJ,E_muJ,T_pk_muJ+273.15,T_ref_par,E_D_muJ,k_par,T_vec)
kap_vec = B_lam(B_0_kap,E_kap,T_pk_kap+273.15,T_ref_par,E_D_kap,k_par,T_vec)

r_m_vec = r_SP_app_lam(bpk_vec, muJ_vec, mu_vec, kap_vec, alp_vec)

#Get the lambdified pd functions

dr_dbpk_lam = lambdify((b_pk, alpha, mu, kappa), dr_dbpk, np)
dr_dalp_lam = lambdify((b_pk, alpha, mu, mu_J, kappa), dr_dalp, np)
dr_dmu_lam = lambdify((b_pk, alpha, mu, mu_J, kappa), dr_dmu, np)
dr_dmuJ_lam = lambdify((alpha, mu, kappa), dr_dmuJ, np)
dr_dkap_lam = lambdify((b_pk, alpha, mu, mu_J, kappa), dr_dkap, np)

dbpk_dT_lam = lambdify((B_0,E,T_pk,T_ref,E_D,k,T), dBdT, np)
dalp_dT_lam = lambdify((B_0,E,T_pk,T_ref,E_D,k,T), dB_invdT, np)
dmu_dT_lam =  lambdify((B_0,E,T_pk,T_ref,E_D,k,T), dB_invdT, np)
dmuJ_dT_lam =  lambdify((B_0,E,T_pk,T_ref,E_D,k,T), dB_invdT, np)
dkap_dT_lam =  lambdify((B_0,E,T_pk,T_ref,E_D,k,T), dBdT, np)

#Evaluate

dr_dbpk_vec = dr_dbpk_lam(bpk_vec, alp_vec, mu_vec, kap_vec)
dr_dalp_vec = dr_dalp_lam(bpk_vec, alp_vec, mu_vec, muJ_vec, kap_vec)
dr_dmu_vec = dr_dmu_lam(bpk_vec, alp_vec, mu_vec, muJ_vec, kap_vec)
dr_dmuJ_vec =  dr_dmuJ_lam(alp_vec, mu_vec, kap_vec)
dr_dkap_vec = dr_dkap_lam(bpk_vec, alp_vec, mu_vec, muJ_vec, kap_vec)

dbpk_dT_vec = dbpk_dT_lam(B_0_bpk,E_bpk,T_pk_bpk+273.15,T_ref_par,E_D_bpk,k_par,
T_vec)
dalp_dT_vec = dalp_dT_lam (B_0_alp,E_alp,T_pk_alp+273.15,T_ref_par,E_D_alp,k_par
,T_vec)
dmu_dT_vec =  dmu_dT_lam(B_0_mu,E_mu,T_pk_mu+273.15,T_ref_par,E_D_mu,k_par,T_vec
)
dmuJ_dT_vec =  dmuJ_dT_lam(B_0_muJ,E_muJ,T_pk_muJ+273.15,T_ref_par,E_D_muJ,k_par
,T_vec)
dkap_dT_vec =  dkap_dT_lam(B_0_kap,E_kap,T_pk_kap+273.15,T_ref_par,E_D_kap,k_par
,T_vec)
```

In [231]:

```
fig = plt.figure(); ax = fig.add_subplot(111)
ax.plot(T_vec-273.15, dbpk_dT_vec, "red");
ax.set_xlim([0,40])
ax.set_ylabel('($d b_{pk}/dT$)', fontsize=14, fontweight = 'bold')

fig = plt.figure(); ax = fig.add_subplot(211)
ax.plot(T_vec-273.15, dr_dbpk_vec, "red");
ax.set_xlim([0,40])
ax.set_xlabel('Temperature', fontsize=14, fontweight = 'bold');
ax.set_ylabel('($dr_m / db_{pk}$)', fontsize=14, fontweight = 'bold')

fig = plt.figure(); ax = fig.add_subplot(211)
ax.plot(T_vec-273.15, dalp_dT_vec, "blue");
ax.set_xlim([0,40])
ax.set_ylabel(r'($d\alpha/dT$)', fontsize=14, fontweight = 'bold')

ax = fig.add_subplot(212)
ax.plot(T_vec-273.15, dr_dalp_vec, "blue");
ax.set_xlim([0,40])
ax.set_xlabel('Temperature', fontsize=14, fontweight = 'bold');
ax.set_ylabel(r'($dr_m/d\alpha$)', fontsize=14, fontweight = 'bold')

fig = plt.figure(); ax = fig.add_subplot(211)
ax.plot(T_vec-273.15, dmu_dT_vec, "cyan");
ax.set_xlim([0,40])
ax.set_ylabel('($d\mu/dT$)', fontsize=14, fontweight = 'bold')

ax = fig.add_subplot(212)
ax.plot(T_vec-273.15, dr_dmu_vec, "cyan");
ax.set_xlim([0,40])
ax.set_xlabel('Temperature', fontsize=14, fontweight = 'bold');
ax.set_ylabel('($dr_m/d\mu$)', fontsize=14, fontweight = 'bold')

fig = plt.figure(); ax = fig.add_subplot(211)
ax.plot(T_vec-273.15, dmuJ_dT_vec, "orange");
ax.set_xlim([0,40])
ax.set_ylabel('($d\mu_J/dT$)', fontsize=14, fontweight = 'bold')

ax = fig.add_subplot(212)
ax.plot(T_vec-273.15, dr_dmuJ_vec, "orange");
ax.set_xlim([0,40])
ax.set_xlabel('Temperature', fontsize=14, fontweight = 'bold');
ax.set_ylabel('($dr_m/d\mu_J$)', fontsize=14, fontweight = 'bold')

fig = plt.figure(); ax = fig.add_subplot(211)
ax.plot(T_vec-273.15, dkap_dT_vec, "purple");
ax.set_xlim([0,40])
ax.set_ylabel(r'($d\kappa/dT$)', fontsize=14, fontweight = 'bold')

ax = fig.add_subplot(212)
ax.plot(T_vec-273.15, dr_dkap_vec, "purple");
ax.set_xlim([0,40])
ax.set_xlabel('Temperature', fontsize=14, fontweight = 'bold');
ax.set_ylabel(r'($dr_m/d\kappa$)', fontsize=14, fontweight = 'bold')
```

Out[231]:

```
Text(0,0.5,'($dr_m/d\\kappa$)')
```

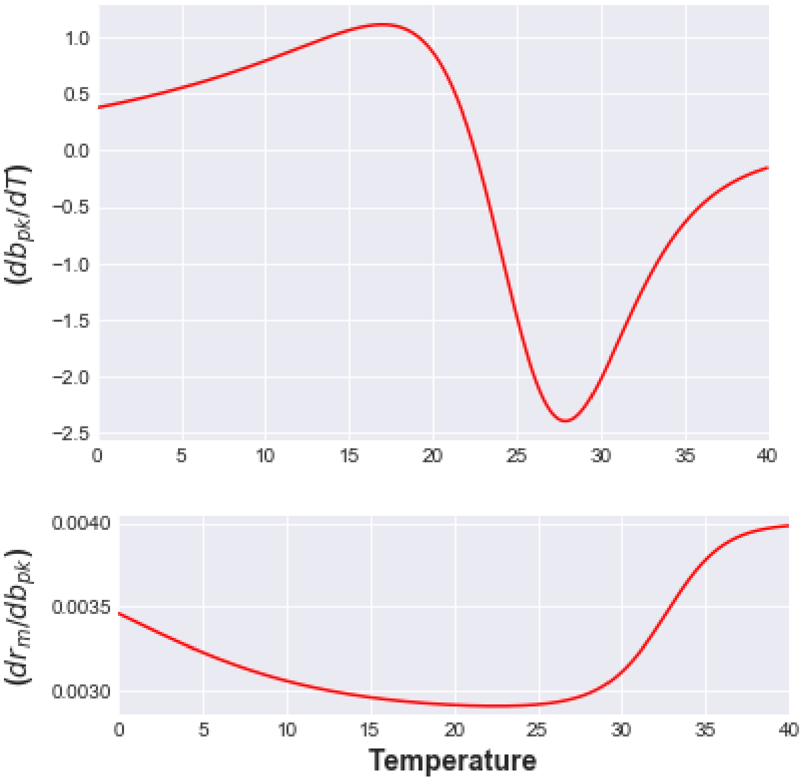


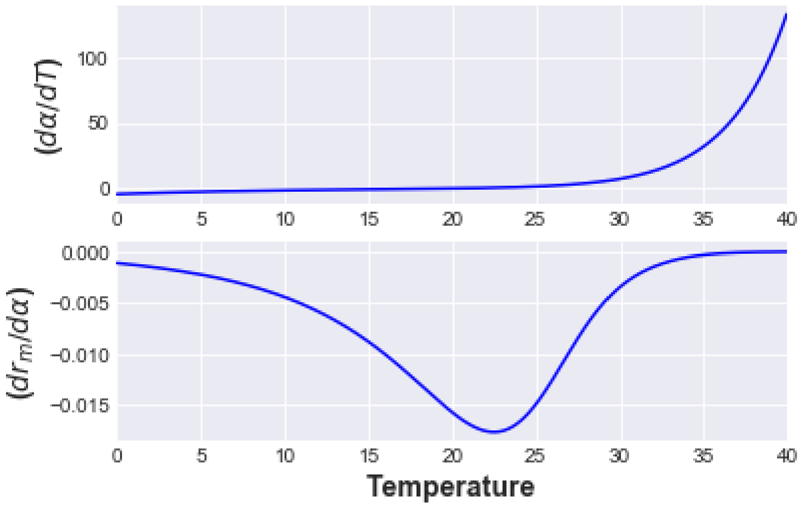

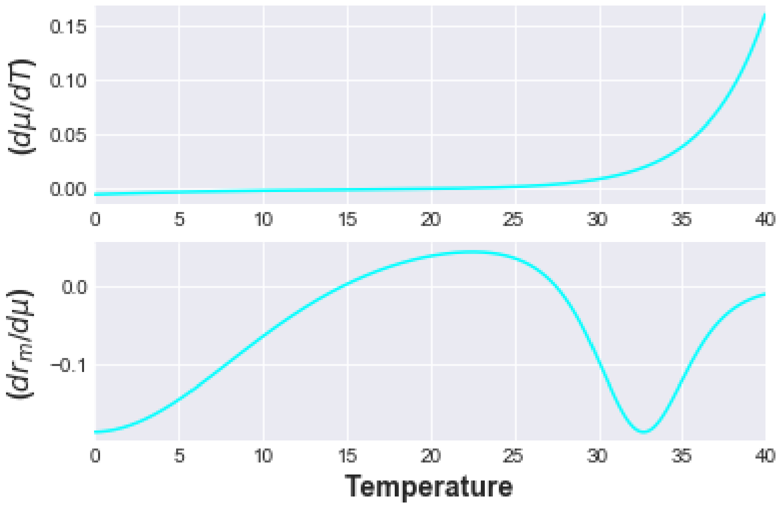
(dμ/dT)
(dr_m/dμ)
Temperature

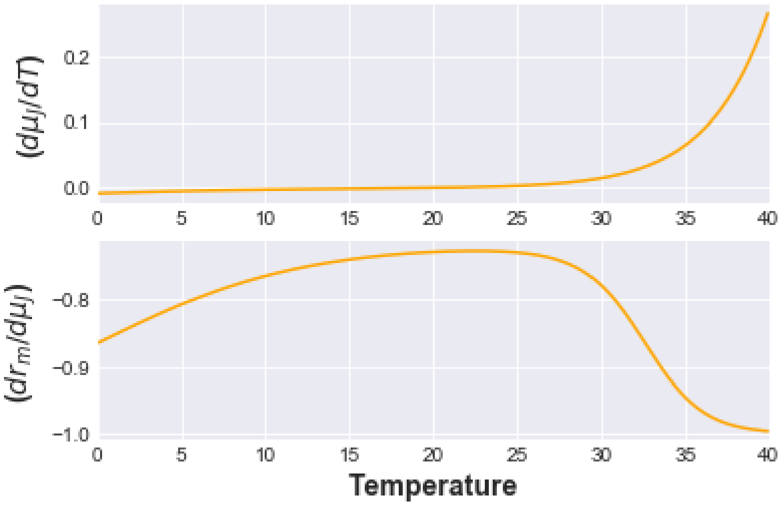
(dμ_l/dT)
(dr_m/dμ_l)
Temperature

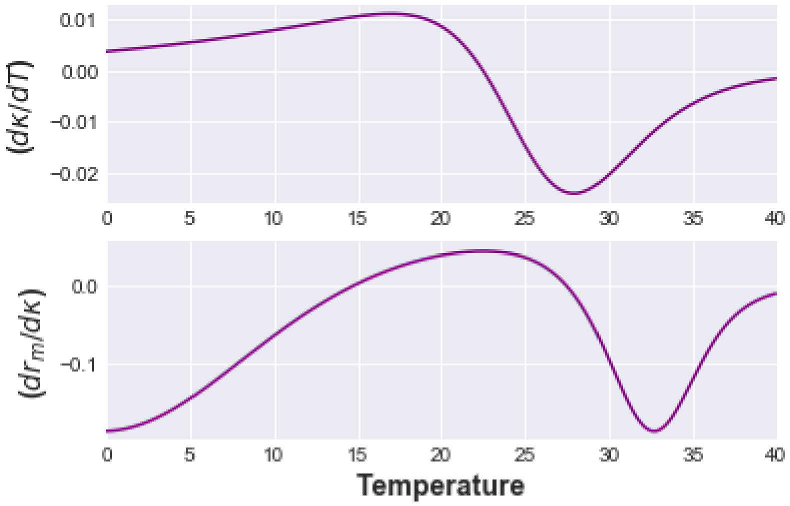
(dκ/dT)
(dr_m/dκ)
Temperature

In [232]:

```
dr_dbpk_dT_vec = dr_dbpk_vec * dbpk_dT_vec
dr_dalp_dT_vec = dr_dalp_vec * dalp_dT_vec
dr_dmu_dT_vec = dr_dmu_vec * dmu_dT_vec
dr_dmuJ_dT_vec = dr_dmuJ_vec * dmuJ_dT_vec
dr_dkap_dT_vec = dr_dkap_vec * dkap_dT_vec

dr_dT_vec = dr_dbpk_dT_vec + dr_dalp_dT_vec + dr_dmu_dT_vec + dr_dmuJ_dT_vec + d
r_dkap_dT_vec
dr_dT_A_vec = dr_dbpk_dT_vec + dr_dmu_dT_vec + dr_dkap_dT_vec
dr_dT_J_vec = dr_dalp_dT_vec + dr_dmuJ_dT_vec
```

Having evaluated the sensitivities, we can plot the relative contribution of each parameter to the TPC for $r_m$. The sensitivity of $r_m$ to the thermal dependence of each parameter can be assessed by deviation of each evaluated partial derivative with respect to each parameter (the differently colored lines) from zero.

Furthermore, traits can be combined (mathematically, summed) together by lifestage, with development rate and juvenile mortality categorised as juvenile traits and adult mortality, fecundity, and rate of loss of fecundity as adult traits. This reveals the relative contribution of the TPC of each lifestage to the TPC of of $r_m$, again with greater deviation from zero indicating a greater contribution.

In [233]:

```
fig = plt.figure(figsize=(10, 4))
ax = fig.add_subplot(121)
ax.set_title('A', fontsize=13, fontweight = 'bold')
ax.plot(T_vec-273.15,dr_dT_vec, 'black',linewidth=1.5)
ax.plot(T_vec-273.15,dr_dbpk_dT_vec, 'red')
ax.plot(T_vec-273.15,dr_dalp_dT_vec, 'blue')
ax.plot(T_vec-273.15,dr_dmu_dT_vec, 'cyan')
ax.plot(T_vec-273.15,dr_dmuJ_dT_vec, 'orange')
ax.plot(T_vec-273.15,dr_dkap_dT_vec, 'purple')
ax.plot([T_pk_par, T_pk_par], [ax.get_ylim()[0],ax.get_ylim()[1]], color='k', li
nestyle='--', linewidth=1)
ax.set_xlim([0,35])
ax.set_ylim([-0.06,0.02])
ax.set_xlabel('Temperature $(^\circ C)$', fontsize=14, fontweight = 'bold');
ax.set_ylabel(r'$\frac{dr_m}{dT}$', fontsize=14, fontweight = 'bold');
plt.legend(['Full', '$b_{pk}$', r'$\alpha$', '$\mu$', '$\mu_J$', r'$\kappa$'], l
oc='lower left') #Add kappa to legend if included in analysis

ax = fig.add_subplot(122)
ax.set_title('B', fontsize=13, fontweight = 'bold');
ax.plot(T_vec-273.15,dr_dT_vec, 'black',linewidth=1.5);
ax.plot(T_vec-273.15,dr_dT_A_vec, 'r--');
ax.plot(T_vec-273.15,dr_dT_J_vec, 'b--');
ax.plot([T_pk_par, T_pk_par], [ax.get_ylim()[0],ax.get_ylim()[1]], color='k', li
nestyle='--', linewidth=1)
ax.set_xlim([0,35])
ax.set_ylim([-0.06,0.02])
ax.set_xlabel('Temperature $(^\circ C)$', fontsize=14, fontweight = 'bold');
plt.legend(['Full', 'Adult', 'Juvenile'], loc='lower left')

plt.savefig('Results/r_sens1.pdf')
```

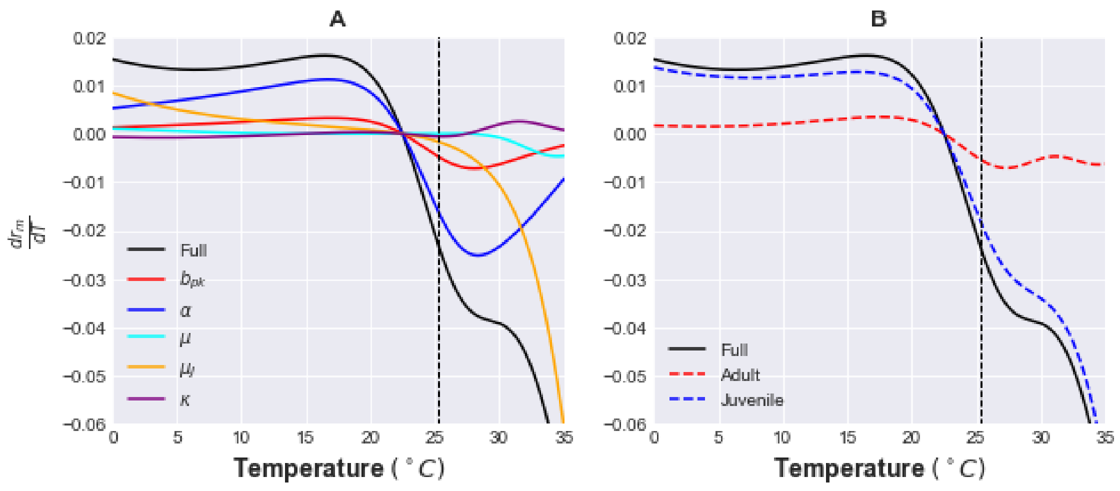


Note that in the above figure, sensitivity of $r_m$ to the thermal dependence of each parameter can be assessed by deviation of each evaluated partial derivative with respect to each parameter (the differently colored lines) from zero. The vertical dashed line is the thermal optimum of $r_m$. Thus juvenile traits make a larger contribution at temperatures below the thermal optimum.

#### A numerical sensitivity analysis

The sensitivity of the $r_m$'s TPC to the the thermal sensitivity of the traits can also be assessed by holding each trait's temperature response constant (no trait variation) in turn. This is essentially a form of non-infinitesimal numerical perturbation analysis (non-infinitesimal because each traits' thermal dependence is held constant instead of being perturbed infinitesimally). We proceed as follows, holding each parameter constant wrt temperature (at its reference value, $T_r$) :

In [234]:

```
r_bpk_cons_vec = r_SP_app_lam(B_0_bpk,muJ_vec, mu_vec, kap_vec, alp_vec)

r_alp_cons_vec = r_SP_app_lam(bpk_vec,muJ_vec, mu_vec, kap_vec, alp_par)

r_mu_cons_vec = r_SP_app_lam(bpk_vec,muJ_vec, mu_par, kap_vec, alp_vec)

r_muJ_cons_vec = r_SP_app_lam(bpk_vec,mu_J_par, mu_vec, kap_vec, alp_vec)

r_kap_cons_vec = r_SP_app_lam(bpk_vec,muJ_vec, mu_vec, kap_par, alp_vec)

r_J_cons_vec = r_SP_app_lam(bpk_vec, mu_J_par, mu_vec, kap_vec, alp_par)

r_A_cons_vec = r_SP_app_lam(b_pk_par,muJ_vec, mu_par, kap_par, alp_vec)
```

Now let's visualize the results:

In [235]:

```
fig = plt.figure(figsize=(10, 4.5))

ax = fig.add_subplot(121)
ax.set_title('A', fontsize=13, fontweight = 'bold');

ax.plot(T_vec-273.15,r_m_vec, 'black',linewidth=1.8);
ax.plot(T_vec-273.15,r_bpk_cons_vec, 'red');
ax.plot(T_vec-273.15,r_alp_cons_vec, 'blue');
ax.plot(T_vec-273.15,r_mu_cons_vec, 'cyan');
ax.plot(T_vec-273.15,r_muJ_cons_vec, 'orange');
ax.plot(T_vec-273.15,r_kap_cons_vec, 'purple');
ax.plot([T_pk_par, T_pk_par], [ax.get_ylim()[0],ax.get_ylim()[1]], color='k', li
nestyle='--', linewidth=1)
ax.set_xlim([0,40])
ax.set_ylim([-0.4,0.3])
ax.set_xlabel('Temperature $(^\circ C)$', fontsize=14, fontweight = 'bold');
ax.set_ylabel('$r_m$', fontsize=14, fontweight = 'bold');
plt.legend(['Full', '$b_{pk}$ const', r'$\alpha$  const', '$\mu$  const', '$\mu_
J$ const',r'$\kappa$ const'], loc='lower center',fontsize=11) #Add kappa to lege
nd if included in analysis

ax = fig.add_subplot(122)
ax.set_title('B', fontsize=13, fontweight = 'bold');
ax.plot(T_vec-273.15,r_m_vec, 'black',linewidth=1.5);
ax.plot(T_vec-273.15,r_J_cons_vec, 'b--');
ax.plot(T_vec-273.15,r_A_cons_vec, 'r--');
ax.plot([T_pk_par, T_pk_par], [ax.get_ylim()[0],ax.get_ylim()[1]], color='k', li
nestyle='--', linewidth=1)
ax.set_xlim([0,40])
ax.set_ylim([-0.4,0.3])
plt.legend(['Full', 'Juvenile traits constant', 'Adult traits constant'], loc='l
ower center',fontsize=11) #Add kappa to legend if included in analysis
ax.set_xlabel('Temperature $(^\circ C)$', fontweight = 'bold',fontsize=14)

fig.tight_layout()

plt.savefig('Results/r_sens2.pdf')
```

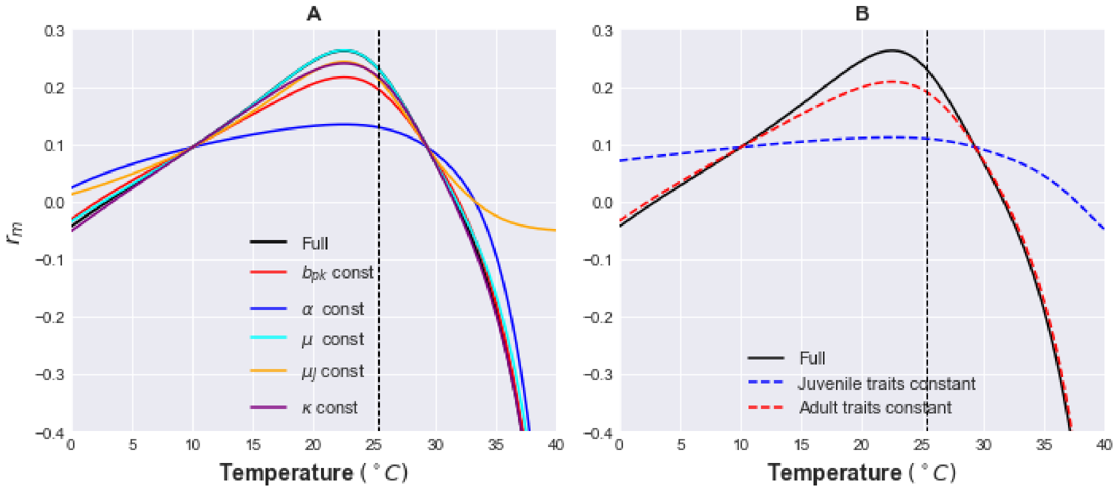

When thus plotting the thermal performance of $r_m$ with each parameter held constant, greater deviation from the full model (where all traits vary with temperature) indicates greater contribution from the thermal dependence of that trait. Again, traits may be grouped by life history stage to assess the combined contribution of thermal sensitivity of each lifestage as we did above. As expected, the results are qualitatively consistent (in terms of contribution of juvenile vs. adult traits to fitness) with the analytical sensitivity analysis above.

Thus, a trait sensitivity analysis within the context of a trait-based VBD framework reveals new, mechanistic insights. In this case, where we have investigated the relative contribution of juvenile and adult traits, it reveals that juvenile traits - traditionally ignored in classical VBD studies are expected to play a major role in determining vector fitness, and therefore abundance, and ultimately transmission.

In particular, the above trait sensitivity analysis adds further insight to why the trait-driven approach yields very different predictions for population abundance $R_0$ compared to a phenomenological approach: the thermal sensitivity of abundance ($V$) and the underlying population fitness ($r_m$) is driven by the variation of temperature-driven variation in larval stage traits. Predictions and insights provide, quantitative targets for validation using field data.

Traits sensitivity analyses also allow key traits can be identified, guiding further empirical and theoretical work on the contributions of traits to VBD system dynamics.

### References

[1] Pawar Samraat, Dell Anthony I. and M. Van, ``*Dimensionality of consumer search space drives trophic interaction strengths*'', Nature, vol. 486, number 7404, pp. 485--489, jun 2012. online (http://dx.doi.org/10.1038/nature11131)

[2] Dell Anthony I, Pawar Samraat and Savage Van M, ``*Temperature dependence of trophic interactions are driven by asymmetry of species responses and foraging strategy*'', J. Anim. Ecol., vol. 83, number 1, pp. 70--84, may 2014. online (http://www.ncbi.nlm.nih.gov/pubmed/23692182)

[3] Gilbert Benjamin, Tunney Tyler D., Mccann Kevin S. *et al.*, ``*A bioenergetic framework for the temperature dependence of trophic interactions*'', Ecol. Lett., vol. 17, number 8, pp. 902--914, 2014.

[4] Samraat Pawar, Anthony I. Dell and Van M. Savage, ``*From metabolic constraints on individuals to the dynamics of ecosystems*'', 2015.

[5] Rizzuto Matteo, Carbone Chris and Pawar Samraat, ``*Foraging constraints reverse the scaling of activity time in carnivores*'', Nat. Ecol. Evol., vol. 2, number 2, pp. 247--253, 2018. online (http://www.nature.com/articles/s41559-017-0386-1)

[6] Vucic-pestic Olivera, Rall C, Kalinkat Gregor *et al.*, ``*Allometric functional response model: Body masses constrain interaction strengths*'', J. Anim. Ecol., vol. 79, number 1, pp. 249--256, 2010.

[7] Dell Anthony I, Pawar Samraat and Savage Van M, ``*Systematic variation in the temperature dependence of physiological and ecological traits*'', Proc. Natl. Acad. Sci. U. S. A., vol. 108, number 26, pp. 10591--10596, jun 2011. online (http://www.ncbi.nlm.nih.gov/pubmed/21606358 http://www.pubmedcentral.nih.gov/articlerender.fcgi?artid=PMC3127911)

[8] Mordecai Erin A., Paaijmans Krijn P., Johnson Leah R. *et al.*, ``*Optimal temperature for malaria transmission is dramatically lower than previously predicted*'', Ecol. Lett., vol. 16, number 1, pp. 22--30, oct 2013.

[9] Mordecai Erin A., Cohen Jeremy M., Evans Michelle V. *et al.*, ``*Detecting the impact of temperature on transmission of Zika, dengue, and chikungunya using mechanistic models*'', PLoS Negl. Trop. Dis., vol. 11, number 4, pp. e0005568, 2017. online (http://dx.plos.org/10.1371/journal.pntd.0005568)

[10] Forster Jack, Hirst Andrew G and Atkinson David, ``*Warming-induced reductions in body size are greater in aquatic than terrestrial species*'', Proc. Natl. Acad. Sci. U. S. A., vol. 109, number 47, pp. 19310--4, nov 2012.

[11] Diekmann O, Heesterbeek Hans and Roberts M G, ``*The construction of next-generation matrices for compartmental epidemic models*'', J. R. Soc. Interface, vol. 7, number 47, pp. 873--885, 2010.

[12] May Robert M. and Anderson Roy M., ``*Population biology of infectious diseases: Part II*'', Nature, vol. 280, number 5722, pp. 455--461, 1979.

[13] Anderson R M and May R M, ``*Population biology of infectious diseases: Part I*'', Nature, vol. 280, number 5721, pp. 361--367, 1979.

[14] Beck-Johnson Lindsay M., Nelson William a., Paaijmans Krijn P. *et al.*, ``*The effect of temperature on Anopheles mosquito population dynamics and the potential for malaria transmission*'', PLoS One, vol. 8, number 11, pp. , 2013.

[15] Amarasekare Priyanga and Coutinho Renato M., ``*The intrinsic growth rate as a predictor of population viability under climate warming*'', J. Anim. Ecol., vol. 82, number 6, pp. 1240--1253, 2013.

[16] Johnson P T J, de Roode J C and Fenton A, ``*Why infectious disease research needs community ecology*'', Science (80-. )., vol. 349, number , pp. 1259504, 2015.

[17] Parham Paul Edward and Michael Edwin, ``*Modeling the effects of weather and climate change on malaria transmission*'', Environ. Health Perspect., vol. 118, number 5, pp. 620--626, may 2010.

[18] Frazier Melanie R., Huey Raymond B. and Berrigan David, ``*Thermodynamics constrains the evolution of insect population growth rates: "warmer is better"*'', Am. Nat., vol. 168, number 4, pp. 512--520, 2006. 19 ({\</p> <p>[<a id=)] Ng Ka-Chon, Chaves Luis, Tsai Kun-Hsien *et al.*, ``*Increased Adult Aedes aegypti and Culex quinquefasciatus (Diptera: Culicidae) Abundance in a Dengue Transmission Hotspot, Compared to a Coldspot, within Kaohsiung City, Taiwan*'', Insects, vol. 9, number 3, pp. 98, aug 2018.

[20] Hoshi Tomonori, Higa Yukiko and Chaves Luis Fernando, ``*Uranotaenia novobscura ryukyuana (Diptera: Culicidae) Population Dynamics Are Denso-Dependent and Autonomous From Weather Fluctuations*'', Ann. Entomol. Soc. Am., vol. 107, number 1, pp. 136--142, jan 2014.

[21] Johnson Leah R., Gramacy Robert B., Cohen Jeremy *et al.*, ``*Phenomenological forecasting of disease incidence using heteroskedastic Gaussian processes: A dengue case study*'', Ann. Appl. Stat., vol. 12, number 1, pp. 27--66, mar 2018.

[22] Peter J. Brockwell and Richard A. Davis, ``*Time Series: Theory and Methods*'', 1991.
